# A unique interplay of access and selection shapes peritoneal metastasis evolution in colorectal cancer

**DOI:** 10.1101/2024.09.25.614736

**Authors:** Emma CE Wassenaar, Alexander N Gorelick, Wei-Ting Hung, David M Cheek, Emre Kucukkose, I-Hsiu Lee, Martin Blohmer, Sebastian Degner, Peter Giunta, Rene MJ Wiezer, Mihaela G Raicu, Inge Ubink, Sjoerd J Klaasen, Nico Lansu, Emma V. Watson, Ryan B. Corcoran, Genevieve Boland, Gad Getz, Geert JPL Kops, Dejan Juric, Jochen K Lennerz, Djamila Boerma, Onno Kranenburg, Kamila Naxerova

## Abstract

Whether metastasis in humans can be accomplished by most primary tumor cells or requires the evolution of a specialized trait remains an open question. To evaluate whether metastases are founded by non-random subsets of primary tumor lineages requires extensive, difficult-to-implement sampling. We have realized an unusually dense multi-region sampling scheme in a cohort of 26 colorectal cancer patients with peritoneal metastases, reconstructing the evolutionary history of on average 28.8 tissue samples per patient with a microsatellite-based fingerprinting assay. To assess metastatic randomness, we evaluate inter- and intra-metastatic heterogeneity relative to the primary tumor and find that peritoneal metastases are more heterogeneous than liver metastases but less diverse than locoregional metastases. Metachronous peritoneal metastases exposed to systemic chemotherapy show significantly higher inter-lesion diversity than synchronous, untreated metastases. Projection of peritoneal metastasis origins onto a spatial map of the primary tumor reveals that they often originate at the deep-invading edge, in contrast to liver and lymph node metastases which exhibit no such preference. Furthermore, peritoneal metastases typically do not share a common subclonal origin with distant metastases in more remote organs. Synthesizing these insights into an evolutionary portrait of peritoneal metastases, we conclude that the peritoneal-metastatic process imposes milder selective pressures onto disseminating cancer cells than the liver-metastatic process. Peritoneal metastases’ unique evolutionary features have potential implications for staging and treatment.

## Introduction

The life history of metastases in humans remains poorly understood. Although recent advances in multi-region sequencing have uncovered important new insights into the dynamics of metastasis formation^1–5^, many foundational questions remain unanswered. One prominent unresolved question concerns the distribution of metastatic potential among primary tumor cells. Are all primary tumor cells similarly likely to become metastasis founder cells? Or do lineages with superior ability to execute at least one of the steps of the metastatic cascade exist^6^? Clonal lineages with variable, mitotically heritable metastatic potential have been demonstrated in mice^7,8^. In human cancer, the situation is less clear, as it has been challenging to identify molecular features that are enriched in metastases over primary tumors^9,10^. A principled search for possible molecular promoters of metastasis (mutations, copy number variants, epigenetic alternations, heritable changes in gene expression) would be greatly enabled if we could ascertain whether human metastases descend from cells with specialized attributes.

If metastases are in fact derived from cells that have acquired specialized pro-metastatic traits, then we would expect them to represent a non-random sample of primary tumor lineages. Conversely, if no specialized trait is required, all primary tumor subclones will be equally likely to give rise to metastases. To test the null hypothesis of metastatic randomness, two elements are required: a detailed picture of the clonal diversity in the primary tumor, and the genotypes of as many distinct metastases from the same patient as possible. If only a single metastasis is analyzed, it is difficult to judge whether the lesion was seeded by a lineage with increased metastatic capacity or whether it originated from a disseminated cell that managed to grow out by chance. If, on the other hand, multiple analyzed metastases all belong to the same lineage, we may suspect with greater confidence that some functional specialization has occurred.

Arguably the greatest hurdle for the study of metastatic randomness is the extreme difficulty of obtaining suitable tissue samples. Metastases are rarely surgically resected. Research autopsies represent valuable opportunities for metastasis collection, but the primary tumor has often been removed at the time of death and may not be available. Obtaining multi-region sampled primary tumors and matched metastases through prospective collection is a daunting task due to the rarity of such cases. Retrospective collection of formalin-fixed and paraffin-embedded (FFPE) samples is more feasible, but it can be technically challenging to work with these specimens, and a patient’s consent to broad sequencing – which exposes germline variants that in conjunction with other data types can be used to identify individuals^11^ – may not be available years after the fact.

Together, these challenges have resulted in a paucity of data that could effectively illuminate the question of metastatic randomness. We have previously circumnavigated some of these problems by reconstructing the evolutionary history of metastatic cancers with a scalable microsatellite fingerprinting assay that is suitable for archival FFPE samples^12–15^. Applying the method to a large cohort of patients with metastatic colorectal cancer, we made a surprising discovery: we found that liver metastases showed strong evidence of non-randomness, usually arising from a small subset of the lineages that were detected in the primary tumor^14^. In contrast, lymph node metastases were sampled from the primary tumor much more randomly. These results suggested that the evolutionary rules of metastasis formation differ from host organ to host organ. Liver metastases are formed by a privileged subset of primary tumor lineages, likely because the liver-metastatic process imposes severe selective constraints onto disseminating tumor cells, restricting successful colonization to only a few subclones with heritably increased liver-metastatic potential. Other environments, like the lymphatic system, seem to represent a friendlier milieu^16,17^ that can be successfully navigated by a larger fraction of primary tumor cells.

Here we investigate the host organ-specific properties of colorectal cancer metastases in one of the most important but largely overlooked sites: the peritoneum. Peritoneal metastasis is frequent – 5.7% of colon cancer patients have peritoneal metastases at the time of diagnosis, and 5.5% more will develop them in the course of their disease^18^, making the peritoneum the most frequent site of distant metastasis after the liver. Peritoneal metastasis patients have a poor prognosis; even if the peritoneum is the only affected site, median survival is a mere 16 months^19^. Despite its frequency, very little is known about the genetics of peritoneal disease in colorectal cancer, and new insights that could ultimately guide more effective treatment strategies are needed. Few comprehensive studies have examined this disease entity, and existing data are mostly gene expression-focused^20,21^. Not only do we not know to what degree peritoneal metastasis is non-random, we also do not know where peritoneal metastases originate and what their relationship to other locoregional and distant metastases is. This study aims to address these questions by leveraging a unique patient cohort with multi-region sampled primary tumors and matched peritoneal metastases.

## Results

### Polyguanine fingerprints reconstruct the evolutionary history of metastatic cancers

To study the evolution of peritoneal metastasis in human colorectal cancer patients, we assembled a retrospective cohort suitable for high density multi-region sampling. To be included in the study, patients had to be diagnosed with colorectal adenocarcinoma, and FFPE resection specimens of the primary tumor and at least one (but ideally multiple, spatially distinct) peritoneal metastases had to be available. For all patients that met these criteria (*n*=24, detailed clinical information in **Supplementary Table 1**), we made every effort to sample *all* surgically resected cancer components as comprehensively as possible. This approach resulted in dense sample coverage: on average, we successfully analyzed 28.8 tissue samples per patient (9.1 primary tumor areas, 10.5 peritoneal lesions, 5.2 locoregional metastases, 3.0 non-peritoneal distant metastases, and one normal tissue control, **Fig. 1a**, sample information in **Supplementary Table 2**). All but two patients had multiple peritoneal metastases. Within the constraints imposed by their specific characteristics (e.g. stromal content), we sampled primary tumors proportionately to their size (**Supplementary Figure 1**). We also acquired high-resolution images of hematoxylin and eosin (H&E) stained sections of all 445 FFPE tissue blocks from these patients, providing detailed microanatomical maps for all samples. We supplemented the cohort with two previously analyzed patients with multi-region sampled primary tumors and multiple peritoneal metastases (C38, C89).^13,14^

**Figure 1.**
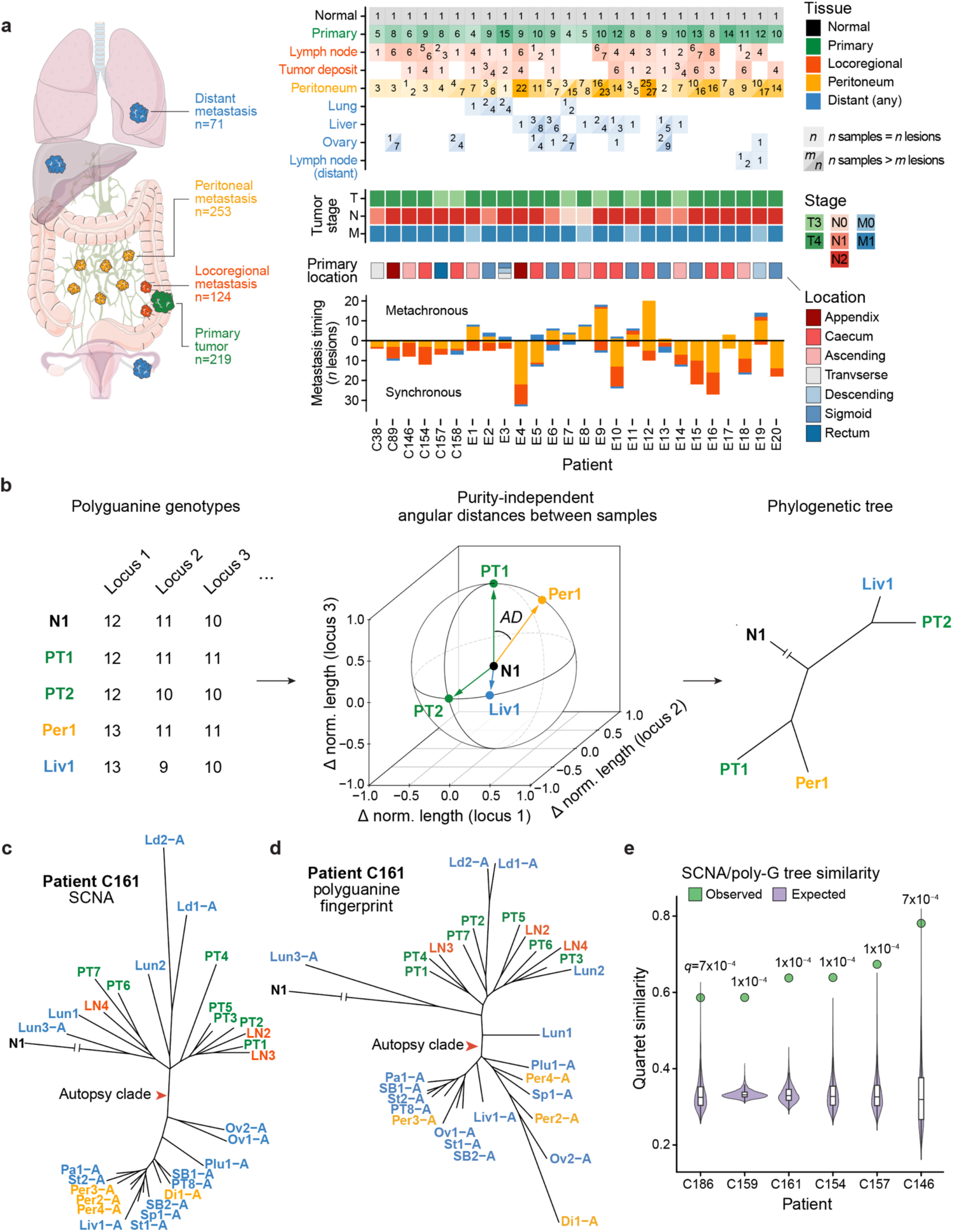
Polyguanine fingerprints reconstruct cancer evolution. **a,** Patient cohort overview. *Left*, Types and total numbers of samples analyzed in this study. *Right*, Overview of samples analyzed per patient, as well as AJCC Tumor, Node, and Metastasis (TNM) staging at diagnosis, primary tumor location, number of analyzed synchronous and metachronous metastatic lesions. **b,** Reconstruction of tumor evolution from polyguanine fingerprints. Each tumor sample is genotyped at approximately 31 polyguanine loci (here, only three are shown for simplicity). Polyguanine fingerprints are created by subtracting the germline genotype from each tumor sample’s vector of mean polyguanine lengths. The angular distance (*AD*) between two unit length-normalized polyguanine fingerprints (e.g. PT1 and Per1) is defined as arccos (*PT*1 · *Per*1). Phylogenetic trees are built from angular distance matrices with the neighbor-joining algorithm. The normal tissue sample is attached post hoc to the last internal node created. **c-d,** Comparison of phylogenetic trees reconstructed from somatic copy number alterations (SCNAs) **(c)** or polyguanine fingerprints **(d)** for patient C161. *PT*, primary tumor; *LN*, lymph node metastasis; *Per*, peritoneal metastasis; *Liv*, liver metastasis; *Lun*, lung metastasis; *Ld*, distant lymph node metastasis; *Di*, diaphragm metastasis; *Ov*, ovarian metastasis; *Pa*, pancreas metastasis; *Plu*, pleural metastasis; *SB*, small bowel metastasis; *Sp*, splenic metastasis; *St*, stomach metastasis. **e,** Quartet similarity between polyguanine and SCNA-based phylogenetic trees for 6 patients. *Green*, observed similarity. *Purple*, similarity expected by chance based on 1,000 random permutations of tree tip labels. Permutation-based *p*-values corrected for multiple-hypothesis testing by Holm’s method (*q*-values).

To reconstruct the evolutionary history of each patient’s cancer, we used polyguanine fingerprinting, a somatic lineage tracing method that relies on detection of insertion and deletion mutations in polyguanine microsatellites^12–15^. Polyguanine repeats mutate at very high rates (μ ≈ 10^-4^ per division per allele^15^) and thus act as natural molecular ‘barcodes’ that record a somatic lineage’s history with high efficiency. In colorectal cancer, interrogation of merely a few dozen tracts is typically sufficient to generate statistically well-supported phylogenetic trees that represent the lineage relationships between the cell populations in each sample^13^. Polyguanine fingerprints are generated by multiplex PCR and fragment analysis without the need for sequencing, providing several important advantages: first, the simplicity of the method allows it to perform robustly even on partially degraded DNA from older clinical FFPE specimens. Second, a research participant’s identity cannot be inferred from the data since no single nucleotide polymorphisms are captured, and no population databases of polyguanine sequences exist. These properties allow us to apply polyguanine fingerprinting to archival clinical samples that are typically off limits to other genetic analysis methods.

A disadvantage of polyguanine fingerprints is that precise tumor purity is difficult to estimate from the data. Prior studies using the methodology therefore had to be restricted to (manually macrodissected) samples of high purity^12–14^. This approach was not feasible for peritoneal metastases, which are often diffusely infiltrated with stromal cells^22^. To enable accurate phylogenetic reconstruction from samples of variable purity, we developed a purity-robust genetic distance. It takes advantage of the fact that the *direction* of a cancer sample’s polyguanine fingerprint (defined as the vector of mean allele lengths for all loci, minus the patient-matched germline genotype) is independent of tumor purity, although its magnitude is not. We therefore use the angle between two cancer sample’s polyguanine fingerprints (the “angular distance”) as a purity-independent measure of genetic divergence (**Fig. 1b**). In a **Supplementary Note**, we offer a detailed mathematical explanation of the angular distance in the context of a random walk model of polyguanine evolution, show simulations to confirm that the angular distance between two tumors with purities greater than 15% approximates the angular distance under optimal conditions (*i.e.*, 100% purity in each tumor) and provide additional details regarding phylogenetic reconstruction, including post-hoc attachment of the normal tissue root.

With the new, purity-robust angular distance integrated into our existing analysis pipeline, which involves extensive quality control, minimum purity filtering, and normalization of polyguanine fingerprints^13,14^ (**Methods**), we next sought to validate the new approach. We collected 30 spatially and temporally distinct samples from a patient with metastatic colorectal cancer, comprising the primary tumor and its associated locoregional lymph node metastases, two surgically resected lung metastases, as well as 19 samples from widely disseminated disease at the time of death (labeled with suffix-A in **Fig. 1c**). We performed low-pass whole genome sequencing (lpWGS, ∼1x depth –of coverage) of all samples and used the resulting data to estimate tumor purity and call somatic copy number alterations (SCNAs) (**Methods**). As expected, purity was variable and ranged between 0.16 and 0.44 (**Supplementary Table 3**). A phylogeny constructed based on purity-corrected SCNA profiles showed an informative topology (**Fig. 1c**): the primary tumor diversified early and locoregional lymph node metastases resembled it closely, a pattern we have described previously^14,15^. Two lung metastases that were resected while the patient was still alive clustered with the primary tumor as well. In contrast, almost all metastases collected at rapid autopsy – except for lung metastasis Lun3-A and distant lymph node metastases Ld1-A and Ld2-A – formed a distinct, homogeneous clade, indicating that widely disseminated disease at the time of death was the result of a recent clonal expansion. The phylogeny based on angular distances among polyguanine fingerprints recovered the same main features (**Fig. 1d**). The trees agreed on all major points, including the significant difference between the primary tumor and most autopsy specimens, the clustering of locoregional metastases with the primary, and the separation of Ld1/2-A and Lun3-A from the rest of the autopsy samples. One difference was a closer relationship between lung metastasis Lun1 and the autopsy sample clade in the polyguanine tree. A comparison of the two phylogenies using quartet similarity, a widely used measure of tree resemblance^23^, showed highly significant concordance (p<1×10^-4^, see **Methods** for details). To establish generality, we repeated the comparison between trees constructed from lpWGS-derived SCNAs and polyguanine fingerprinting for five more patients and 119 samples of variable purity (**Supplementary Table 3**) and observed the same high level of reproducibility in all cases (summary in **Fig. 1e**, all polyguanine- and SCNA-based phylogenies are shown in **Supplementary Fig. 2**). Overall, these data suggest that angular distance trees handle purity differences adequately and provide robust phylogenetic information.

### Peritoneal metastases are genetically more diverse than liver metastases

We reconstructed angular distance-based phylogenies for all patients in the peritoneal metastasis cohort and examined their topologies (**Supplementary Fig. 3**). The first question we hoped to answer concerned metastatic randomness. Do anatomically distinct peritoneal metastases generally resemble each other more than they resemble the primary tumor (low inter-metastatic diversity, **Fig. 2a**)? Or are their genotypes divergent, matching different primary tumor areas more than each other (high inter-metastatic diversity, **Fig. 2b**)? The first scenario would be indicative of peritoneal metastasis formation by only one (or a small subset) of the lineages that are present in the primary tumor, while the second scenario would be consistent with many different primary tumor subclones colonizing the peritoneum in parallel. If only a small number of subclones contributes to peritoneal disease, we can conclude that either access to the peritoneal cavity must be restricted to limited portions of the primary tumor, or that stringent selection prevents the outgrowth of all but a few (specialized) lineages.

**Figure 2.**
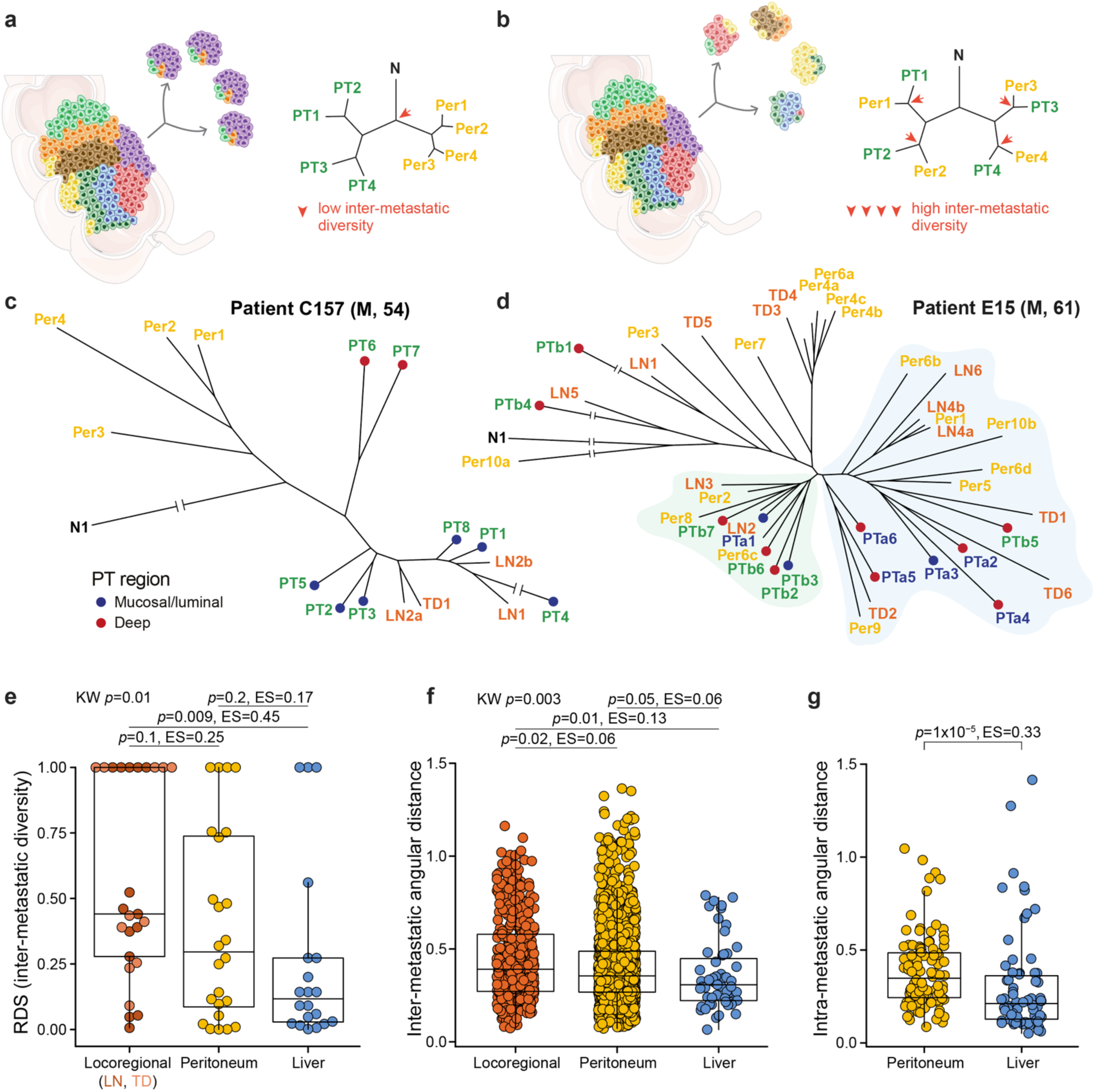
Peritoneal metastases exhibit intermediate inter-metastatic diversity. **a-b,** Schematic illustrating low **(a)** or high **(b)** inter-metastatic diversity in two hypothetical patients. Colored cells represent distinct lineages originating in the primary tumor. **c-d,** Phylogenetic trees for patients C157 and E15, illustrating low **(c)** and high **(d)** inter-metastatic diversity of peritoneal lesions. *Td*, tumor deposit; all other sample type abbreviations as in Figure 1. Spatial localization of primary tumor samples (deep-invasive or mucosal/luminal) is indicated in blue and red. In **(d)**, clades enriched for two spatially distinct primary tumors Pta and PTb are shaded in blue and green, respectively. **e,** Metastasis-specific root diversity scores (RDS) for locoregional, peritoneal, and liver metastases. Each point represents a patient. Lymph node metastases and tumor deposits are evaluated separately but plotted together as locoregional metastases. Kruskal-Wallace *p*-value is shown, along with Dunn’s test *p*-values for each pairwise comparison with Holm’s correction for multiple hypothesis testing. Effect sizes are based on Wilcoxon Rank Sum tests run independently for each pairwise comparison. **f,** Comparison of inter-metastatic diversity by pairwise angular distances. Each point is the angular distance between a pair of distinct metastatic lesions of the indicated type within a patient. Values in the locoregional category include all pairwise distances between lymph node metastases and tumor deposits. *P*-values and effect sizes as in **(c)**. **g,** Comparison of *intra*-metastatic diversity quantified by pairwise angular distances. Each point is the angular distance between a pair of spatially distinct samples taken from the same metastatic lesion. Only metastatic lesions with 2 or more sampled region are included. Wilcoxon rank sum test *p*-value and effect size.

Inspecting tumor phylogenies, we found examples of both high and low inter-metastatic heterogeneity in the peritoneum. For example, four synchronous, spatially distinct peritoneal metastases (Per1-4, located in the omentum and hemidiaphragm) from patient C157 had a recent common ancestor that clearly segregated away from the primary tumor and its associated locoregional lymph node metastases (**Fig. 2c**). The tree topology indicated that all peritoneal metastases had a relatively similar genetic composition and had descended from lineages that were not readily detectable in the primary tumor. Patient C157 was one of the few patients in our cohort who had provided informed consent for broad next-generation sequencing, we could thus additionally examine their cancer with lpWGS and deep whole exome sequencing. We found that all three methods agreed: peritoneal metastases were enriched for a unique clonal population that was not present at appreciable frequencies in any other analyzed samples (**Supplementary Figs. 2 and 4**).

In contrast, patient E15’s peritoneal metastases exhibited extensive inter-metastatic heterogeneity (**Fig. 2d**). Metastases in this case were also synchronous and had been resected along with two T4 stage primary tumors that grew 7 cm apart in the sigmoid colon. Using a recently established classification methodology^15^, we found that the two primaries had a common clonal origin (**Supplementary Fig. 5a** shows this classification for E15 and all other patients who had more than one primary tumor). Subclonal diversity in tumor PTb was significantly larger than in tumor PTa (**Supplementary Fig. 5b**), indicating that it was older^24^ and had likely seeded PTa, a hypothesis that was also consistent with PTb’s larger size (4.5 cm vs. 1.5cm for PTa). While 5 out of 6 PTa samples clustered in a common clade (along with area PTb5 which might have been involved in the genesis of PTa), the patient’s ten peritoneal metastases were randomly distributed throughout the tree, indicating that they were as clonally diverse as the two primary tumors.

To quantify inter-metastatic heterogeneity (i.e. genetic diversity between anatomically distinct metastases with respect to the multi-region sampled primary tumor) across all patients, we employed our previously developed ‘root diversity score’ (RDS) mathematical framework^14^. The RDS quantifies the probability of observing monophyletic clades containing *l* metastases of the same type on phylogenetic trees with *m* total metastases of that type and *k* other tumor samples. For example, patient C157’s tree contains *m*=4 peritoneal metastases and *k*=12 samples that are not peritoneal metastases (8 primary tumor samples and 4 locoregional metastasis samples, **Fig. 2c)**. All peritoneal metastases form a monophyletic clade (*l*=4). The probability that this 4-out-of-4 clustering of peritoneal metastases occurs by chance alone is given by the RDS and is 7.7×10^-4^ in this case. Thus, low RDS values indicate that metastases are genetically homogeneous with respect to the rest of the cancer, while high values indicate polyphyletic metastasis origins and high inter-metastatic diversity. Here, we calculated RDS values in a metastasis-specific manner: we constructed trees with only samples from the primary tumor and the metastasis type under investigation and compared the cohort-wide RDS distributions for locoregional, peritoneal and liver metastases (see **Methods** for more details, **Supplementary Table 5** for all metastasis-specific RDS values). Since few individuals in the this study had liver metastases (their presence was not among the selection criteria), we included data from 30 previously published^13,14^ liver metastasis patients in this analysis.

We observed that locoregional metastases – encompassing lymph node metastases and tumor deposits – were genetically diverse (high RDS values), while liver metastases were more homogeneous (low RDS values, **Fig. 2e**), as we had previously noted^14^. Peritoneal metastasis RDS values fell in-between, indicating intermediate diversity in the peritoneum. We obtained the same result when comparing pairwise inter-lesion angular distances between locoregional, peritoneal and liver metastases (**Fig. 2f**). Leveraging a considerable number of multi-region sampled metastatic lesions, we also compared *intra*-lesion angular distances between peritoneal and liver metastases and found that peritoneal metastases also contained greater clonal diversity *within* individual lesions (**Fig. 2g**). Although peritoneal metastasis is clinically viewed as a form of distant metastasis (defining stage IV disease), its progression mechanism has long been suspected to be more of a locoregional nature, potentially involving direct seeding through breaches in the serosal membrane^25,26^. It is therefore noteworthy that broadly, genetic diversity among peritoneal metastases is intermediate compared with locoregional and ‘true distant’ liver metastases, potentially indicating that seeding to farther organs imposes more severe selective pressures and thus reduces genetic heterogeneity.

### Genetic diversity is increased among metachronous peritoneal metastases exposed to systemic chemotherapy

Next, we wanted to dissect the possible influence of clinical history and treatment on genetic diversity across the different metastasis types. We had previously found that the stark disparity in inter-metastatic diversity between lymph node and liver metastases was most pronounced for synchronous, untreated lesions, indicating that these differences were driven by inherent biology rather than timing or treatment effects^14^. Subsetting root diversity scores to retain only synchronous, untreated lesions, we again found that the RDS distributions of locoregional and liver metastases remained significantly different, with no diminishment in effect size (**Fig. 3a**). Root diversity scores in the peritoneal group remained in their intermediate position but shifted towards lower values, prompting us to investigate explicitly whether inter-metastatic diversity in the peritoneum varied as a function of clinical history.

**Figure 3.**
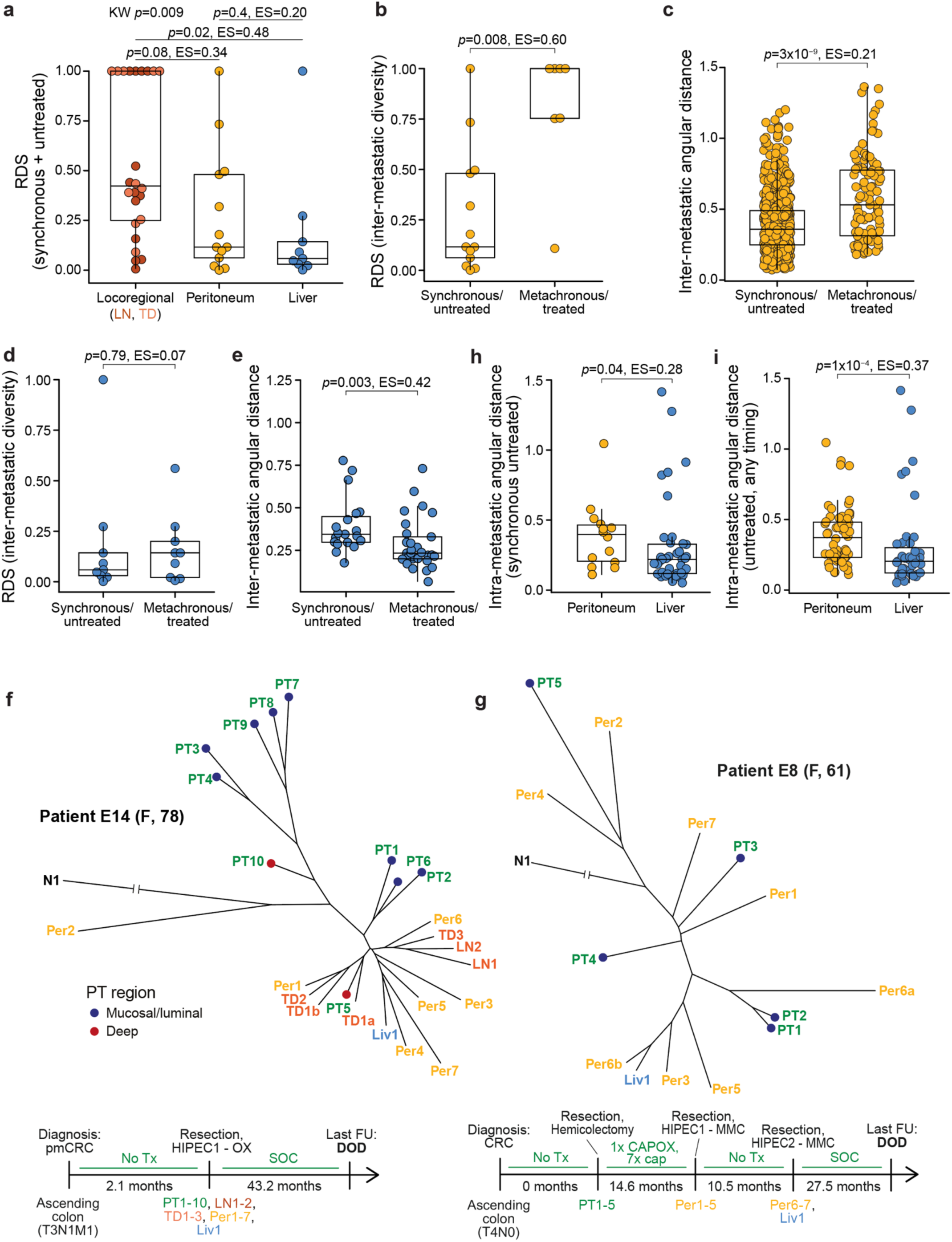
Inter-metastatic diversity varies by timing and treatment. **a,** Inter-metastatic diversity (RDS) of synchronous/untreated locoregional, peritoneal, and liver metastases. RDS calculations are based on a reduced phylogeny consisting only of the patient’s primary tumor, normal tissue, and synchronous/untreated metastases of the indicated tissue type. *P*-values and effect sizes as described in Fig. 2e. **b-c,** Inter-metastatic diversity among synchronous/untreated and metachronous/treated peritoneal metastases based on RDS **(b)** and pairwise inter-lesion angular distances **(c)**. **d-e,** As in (b-c) but for liver metastases. **f,** Phylogenetic tree and clinical timeline for patient E14. pmCRC, colorectal cancer with peritoneal metastasis; OX, oxaliplatin. **g,** Phylogenetic tree and clinical timeline for patient E8. MMC, mitomycin C; CAPOX, capecitabine and oxaliplatin. **h,** Intra-metastatic diversity in synchronous/untreated peritoneal and liver metastases. Intra-metastatic diversity is quantified as in Fig. 2g. **i,** As in (h), but including all untreated peritoneal and liver metastases regardless of timing. *P*-values and effect sizes for all comparisons between two groups (b-e, h, i) are from Wilcoxon rank sum tests.

The two main variables of interest – treatment history and metastasis timing – could not be studied separately since synchronous metastases were almost always untreated and metachronous metastases were almost always treated in this cohort. We did have a small number of synchronous/treated or metachronous/untreated metastases, but the number of patients in these categories never exceeded two, precluding meaningful statistical analysis (**Supplementary Table 4**). All subsequent comparisons therefore focus on metastases that were resected along with the primary tumor and had never been exposed to any kind of treatment (labeled “synchronous/untreated” from now on) and metastases that were resected in a separate surgery more than three months after primary tumor resection and had experienced systemic chemotherapy exposure (labeled “metachronous/treated”).

Comparing inter-lesion diversity between synchronous/untreated and metachronous/treated peritoneal metastases, we found large and significant differences for both RDS (**Fig. 3b**) and inter-lesion angular distances (**Fig. 3c**). The effect was specific to the peritoneum, as the RDSs of synchronous/untreated and metachronous/treated liver metastases were similar (**Fig. 3d**), and the inter-lesion angular distances even were diminished in the treated/metachronous setting (**Fig. 3e**). Locoregional metastases are almost always resected at the same time as the primary tumor, the comparison could thus not be performed for these lesions.

The cohort-wide patterns summarized in **Figs. 3b-c** were easily visible on the patient level. For example, the phylogenetic tree of patient E14 – whose primary tumor, locoregional metastases (*n*=5), peritoneal metastases (*n*=7) and liver metastasis (*n*=1) were resected synchronously with no neoadjuvant treatment – showed that most peritoneal metastases were closely related to each other (peritoneal metastasis-specific RDS=0.1, **Fig. 3f)** and resembled primary tumor area PT5. The remaining nine primary tumor regions located to other branches of the phylogenetic tree and did not appear to share any direct ancestry with the peritoneal metastases. Notably, in this patient, all locoregional metastases as well as the liver lesion also originated from that same metastatic lineage.

In contrast, peritoneal metastases were highly diverse in patient E8 who had undergone a primary tumor resection with curative intent and had then been treated with adjuvant systemic chemotherapy (**Fig. 3g**). Fourteen and 25 months after the initial surgery, the patient completed two cytoreductive surgeries and hyperthermic intraperitoneal chemotherapies (HIPECs) in which five and two peritoneal metastases were resected, respectively. In this patient, peritoneal metastases were intermixed with various primary tumor areas on the phylogenetic tree and there appeared to be no distinct peritoneal-metastatic lineage. Even multiple samples from the same metastatic lesion (Per6a,b) did not cluster together, indicating high degrees of subclonal diversity.

Treatment drives cancer evolution^27^, potentially explaining elevated levels of inter-metastatic diversity in metachronous/treated peritoneal metastases. However, we were surprised to find that the presumed treatment effect was host organ-specific, as no elevated diversity could be detected for metachronous/treated liver metastases. We hypothesized that this observation could be explained by exacerbation of *pre-existing* clonal diversity through chemotherapy-induced cell death and regrowth cycles which amplify heterogeneity via genetic drift. This reasoning is attractive because it relies on minimal assumptions, requiring only that cancer cells die as a consequence of chemotherapy. Pursuing this hypothesis, we returned to a closer examination of the baseline differences that exist between peritoneal and liver metastases. Focusing exclusively on lesions that were resected at the same time as the primary tumor and had not experienced any kind of treatment, we found that – as for all peritoneal and liver metastases, shown in **Fig. 2g** – *intra*-lesion heterogeneity was significantly higher in the peritoneum (**Fig. 3h**). The same was also true for all untreated lesions regardless of timing (**Fig. 3i**). Therefore, in their ‘natural’ untreated state, peritoneal metastases contain more intra-lesion clonal diversity than liver metastases.

We wondered whether the diversity gap between peritoneal and liver metastases was a general biological property that could be recapitulated in a mouse model of colorectal cancer. We transduced patient-derived organoids (PDOs) with multi-color lentiviral LeGO^28^ vectors to create artificial ‘color lineages’ that could be tracked by fluorescence microscopy. We implanted the cells in the caecum of immunocompromised mice and imaged spontaneously arising peritoneal and liver metastases (**Methods**). We observed high color heterogeneity within peritoneal metastases, while liver metastases were mostly uniform in color (**Supplementary Fig. 6**). Unbiased quantification of intra-lesion color diversity across all images via Simpson’s diversity index showed significantly higher diversity in the peritoneum.

Together, these data demonstrate that intra-lesion diversity is higher in the peritoneum than in the liver in the untreated setting, both in the mouse and in humans. To illustrate how chemotherapy might shape this baseline diversity, we considered a simple stochastic ‘toy’ model of cell death and regrowth (**Methods**). In this model, metastases are created and populated with subclones *in silico*. Metastases in one class are assigned high levels of intra-lesion diversity, while metastases in the other class are assigned low levels (**Supplementary Fig. 7**). Chemotherapy is then simulated as the random death of 80% (or 40%) of cells in each lesion. After therapy, each metastasis is regenerated through a random birth-death process and intra- and inter-lesion diversity are recorded. As predicted by the laws of genetic drift, we observe that chemotherapy-related death and regrowth fuel inter-lesion diversification among metastases. The most dramatic increases are seen for metastases that harbor more intra-lesion diversity at baseline (**Supplementary Fig. 7**), in line with our experimental data. Due to the universality of the underlying principle (namely: following the death of a fraction of cells, regrowth will invariably remodel clone frequencies through drift, with the greatest absolute impact on lesions with higher baseline heterogeneity), these results are qualitatively robust to large variation in all model parameters.

We conclude that repeated cycles of cell death and regrowth can exacerbate pre-existing genetic heterogeneity through neutral drift. While our simple model is potentially sufficient to explain why peritoneal metastases show greater post-treatment heterogeneity than liver metastases, we can of course not exclude the possibility of other, more complex factors at play, such as treatment-related selection that varies systematically across clones or host organs. Also, since our data cannot formally clarify whether treatment or metastasis timing is responsible for the high inter-lesion diversity among metachronous/treated peritoneal metastases, effects related to the timing of resection should be considered as well. For example, it is possible that seeding of metachronous metastases occurs by a distinct route (e.g. through release of cancer cells during surgery^29^), which could affect their clonal diversity.

### Clones on the deep-invading tumor edge are closely related to peritoneal metastases

From the data presented thus far, a first evolutionary portrait of colorectal cancer peritoneal metastases is emerging. In comparison with liver metastases, peritoneal metastases contain relatively high levels of intra-lesion genetic diversity (**Fig. 3h-i**). However, at least in the synchronous/untreated setting, anatomically distinct peritoneal metastases also tend to form monophyletic clades (**Fig. 3b**), meaning that they frequently resemble each other. Taken together, these two observations suggest that peritoneal metastases are typically composed of multiple subclones and that these subclones are distributed across peritoneal lesions in similar proportions, leading to relative uniformity *between* metastases (demonstrated for example for patient C157 in **Supplementary Fig. 4**). Many phylogenies further show that most primary tumor lineages are not directly related to peritoneal subclones in the synchronous/untreated setting (e.g. **Fig. 3f**).

These observations raise the question of what is special about the clones that form peritoneal metastases. One possibility is that these lineages acquired a novel trait that allowed them to colonize the peritoneum. Alternatively, their specialization could be of a spatial nature: they could be preferentially located on the deep-invading edge of the primary tumor and have increased access to the peritoneal cavity (**Fig. 4a**). Clinical data^30^ demonstrate that patients with T4 stage primary tumors, which by definition have breached the peritoneal lining, are particularly likely to have synchronous peritoneal metastases (**Fig. 4b**). In contrast, most patients with isolated synchronous liver metastases have T3 stage primary tumors. These data suggest a strong and specific association between peritoneal metastasis and local invasion. However, no direct evidence for a local seeding mechanism exists so far in humans. Since some extra-abdominal malignancies such as lung and breast cancer also metastasize to the peritoneum^26^, the possibility of hematogenous metastasis (and organotropism) cannot be excluded^25^. We reasoned that our dense multi-region sampling scheme might be able to pinpoint the spatial localization of peritoneal metastasis origins in the primary tumor.

**Figure 4.**
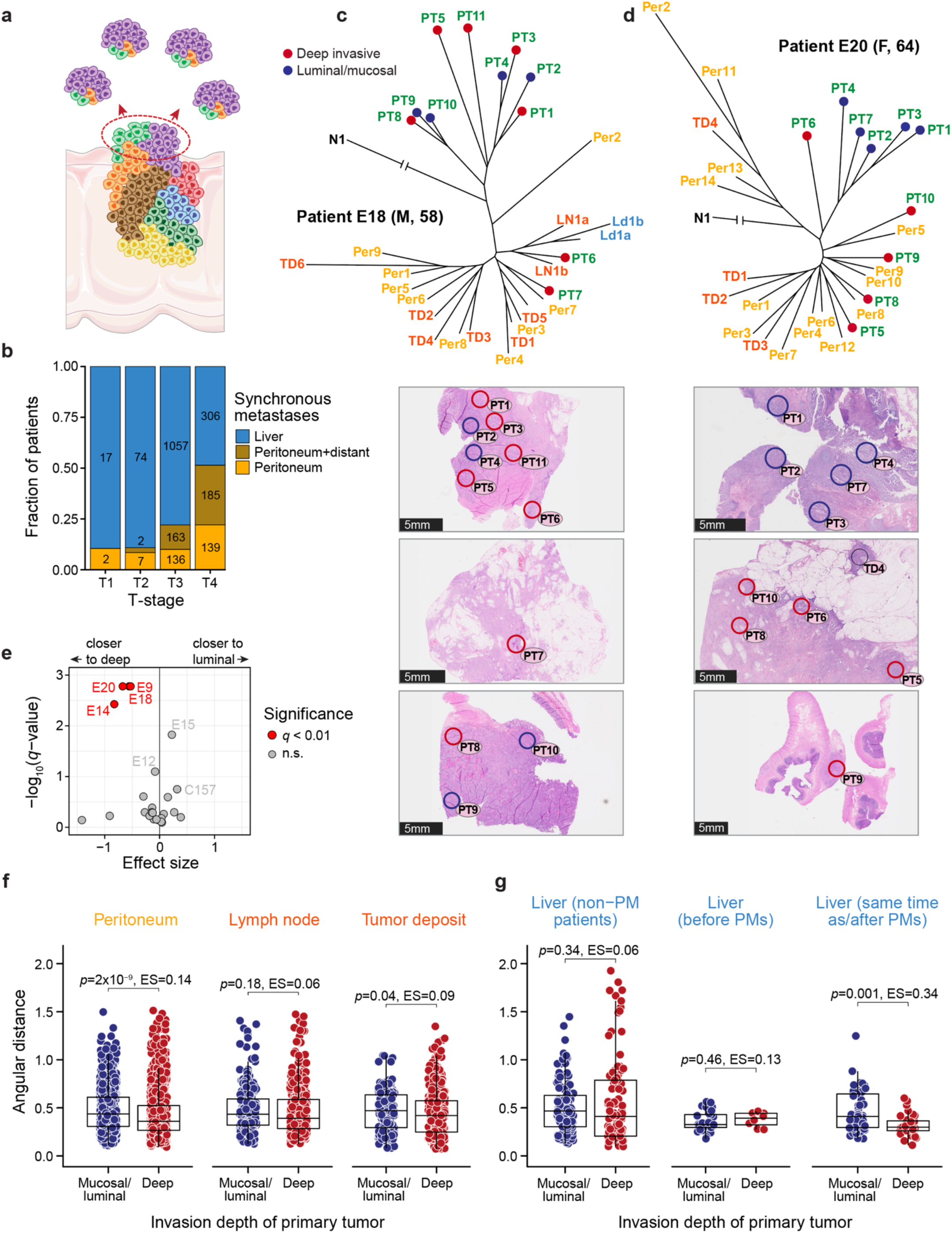
Peritoneal metastases associate with deep-invading primary tumor regions. **a,** Schematic of a T4 stage primary tumor breaching the peritoneal lining (red highlighted region) and seeding peritoneal metastases that are enriched for lineages that are in the breach area **b,** Metastasis types observed in stage IV patients stratified by T-stage. Bars are labeled with the number of patients in each category. Data adapted from Lemmens et al.^30^. **c-d,** Phylogenetic trees for patients E18 **(c)** and E20 **(d)** along with histological images showing the precise anatomical location of primary tumor samples. *Red circles*, deep-invading regions. *Blue circles*, luminal/mucosal regions. **e,** Association of peritoneal metastases (as a group) with deep-invading vs. luminal primary tumor for each patient. For each peritoneal metastasis, we calculate the ratio of its angular distances to the closest deep-invading and closest luminal/mucosal region (lesion-depth ratio). This value is then averaged across all lesions to quantify their overall proximity to deep-invading vs. luminal regions. x-axis: log_2_-ratio of the *observed* average lesion-depth ratio to the *expected* average lesion-depth ratio (median of 10,000 permutations of primary tumor regions’ invasion-depth labels within each patient). y-axis:-log_10_ *p*-values from two-sided permutation tests for each patient, with correction for multiple hypothesis testing (*q*-values). Patients with significant peritoneal metastasis similarity to either deep-invading or luminal/mucosal regions are highlighted in red. **f,** Pairwise angular distances between metastases and deep-invading vs. luminal/mucosal primary tumor regions. Each point is the angular distance between a metastasis and a primary tumor region of the indicated invasion depth. All unique combinations of metastases and primary tumor regions within the same patient are included. *p*-values and effect sizes from two-sided Wilcoxon rank sum tests. **g,** Pairwise angular distances between liver metastases and deep-invading vs. luminal/mucosal primary tumor regions, separated by liver metastasis timing with respect to the earliest diagnosed peritoneal metastasis (PM). *Left*, patients with no peritoneal metastases, only liver metastases. *Center,* liver metastases diagnosed at least 3 months before peritoneal metastases. *Right*, liver metastases diagnosed at the same time or after peritoneal metastases.

To determine whether deep-invading primary tumor regions were more likely to be related to peritoneal metastases than luminal areas, we returned to the H&E images annotated with precise sampling locations. A board-certified gastrointestinal pathologist reviewed the images and classified each primary tumor sample as belonging to the “luminal” or “deep-invading” edge. Several interesting observations emerged after we overlayed these annotations onto the phylogenetic trees. First, we noticed that in many patients who exhibited overall clear segregation of primary tumor and peritoneal metastases, a small subset of sampled primary tumor regions separated from the remainder and clustered among peritoneal metastases. Without exception, such ‘runaway’ primary tumor samples were from the deep-invading edge. For example, patient E14, who was already introduced in **Fig. 3f** as a representative example of low inter-metastatic heterogeneity, showed a strong association between deep-invading area PT5 and a metastatic lineage that was present across multiple host sites (peritoneum, liver, locoregional lymph nodes and tumor deposits). Examining this patient’s H&E images, we observed that PT5 was the only sampled primary tumor region that directly abutted and invaded into the pericolonic fat (**Supplementary Fig. 8**). It is therefore plausible that this region would be the ancestor of all seven synchronous peritoneal metastases in this patient.

Almost identical patterns were observed in several other cases. In patient E18, primary tumor areas PT6 and PT7 – the deepest-invading of all 11 sampled regions – were genetically clearly distinct from the rest of the primary tumor and perfectly matched a large metastatic clade which comprised 8 anatomically distinct peritoneal metastases, 7 locoregional metastases as well as a distant lymph node metastasis (**Fig. 4c**). In Patient E20, four deep-invading regions (three of them sampled from a front of primary tumor that was abutting and pushing into the pericolonic fat) clustered with 10 peritoneal metastases and three tumor deposits, while all five luminal samples formed an unrelated branch elsewhere on the phylogenetic tree (**Fig. 4d**).

To quantify the association between the deep-invading edge and clusters of peritoneal metastases more formally, we conducted a permutation-based statistical test which evaluated whether peritoneal metastases *as a group* were more closely related to the deep-invading or luminal primary tumor in each patient (**Fig. 4e**, with details in legend). This analysis confirmed a significant (q<0.01) clustering of peritoneal metastases with the deep-invading tumor edge in multiple patients, while no such clustering was observed for luminal primary tumor regions. Similarly, we found no significant association between the deep invading edge and groups of lymph node or liver metastases; one patient’s (E20) tumor deposits scored in the significant range (**Supplementary Fig. 9**).

Since the analysis above rewards scenarios in which *multiple* lesions of a given type (peritoneal, locoregional, liver) are non-randomly associated with either the deep-invading or luminal edge, we wanted to provide an additional quantification that directly assesses the proximity of *individual* metastatic lesions to different parts of the primary tumor. We therefore evaluated all pairwise angular distances from metastatic lesions to their matching deep-invading and luminal primary tumor regions. We found that also in this global view, peritoneal metastases had significantly shorter distances to deep-invading primary tumor samples (**Fig. 4f**). Lymph node metastases, on the other hand, were equally closely related to deep-invading and luminal areas, suggesting that these lesions do not preferentially originate at the invasive front. Tumor deposits showed an intermediate association with the deep edge. Liver metastases exhibited a noteworthy pattern. They were not preferentially associated with the luminal or deep-invading edge in patients who had no peritoneal metastases, or in patients in whom peritoneal metastases occurred after the liver lesions had already been resected. However, we did see an association between liver metastases and the deep-invading edge in patients who had prior or synchronous peritoneal metastases (**Fig. 4g)**. This observation raised the question whether liver metastases in these patients might have been seeded by peritoneal metastases – a plausible hypothesis given that the visceral peritoneum (which makes up 70% of the peritoneal surface), drains into the portal vein. Disseminating cancer cells that originated at the deep-invading edge of the primary tumor and colonized the visceral peritoneum would thus encounter the liver as their first capillary bed. We therefore turned to a closer examination of the lineage relationship between peritoneal and distant metastases.

### Most peritoneal metastases do not share a common evolutionary origin with distant metastases

Do peritoneal lesions have distinct subclonal origins from ‘true’ distant metastases in more remote organs like the liver or the lungs, or are all stage IV-defining metastatic lesions clonally related? Examining patient phylogenies, we found several cases in which peritoneal and distant metastases clearly did share a common subclonal origin. For example, peritoneal metastases were closely related to distant metastases in patient E7 who received surgery for a primary tumor in the caecum and synchronous metastases to both ovaries. The patient was disease-free until she relapsed almost 4 years later with metastases to the peritoneum and the lungs (detailed clinical timeline in **Supplementary Fig. 3**). The phylogeny showed a clear separation between the primary tumor and all metastases (**Fig. 5a**) and, in conjunction with the patient’s clinical history, suggested that the ovarian lesions could have been involved in seeding peritoneal metastases. This scenario seems particularly plausible because the primary tumor was T3 stage and thus had not visibly penetrated the peritoneal lining yet. In contrast, most of patient E5’s synchronous peritoneal metastases appeared clonally unrelated to their synchronous ovarian and metachronous liver metastases (**Fig. 5b**). The liver metastases consisted of three anatomically distinct lesions (Liv1-3) that were resected less than one year after the primary tumor. Although they were recovered from different liver segments, they clustered in a tight monophyletic clade, as we often observe for liver metastases (**Fig. 2e**). The ovarian metastasis was much more clonally diverse, with its different multi-region samples (OvH1a-d) mapping to different branches of the phylogenetic tree. Two peritoneal metastases (Per3-4) appeared clonally related to parts of the ovarian metastasis and deep-invading primary tumor region PT10, but most peritoneal lesions located to distinct clades that had no special relationship to any of the distant-metastatic samples.

**Figure 5.**
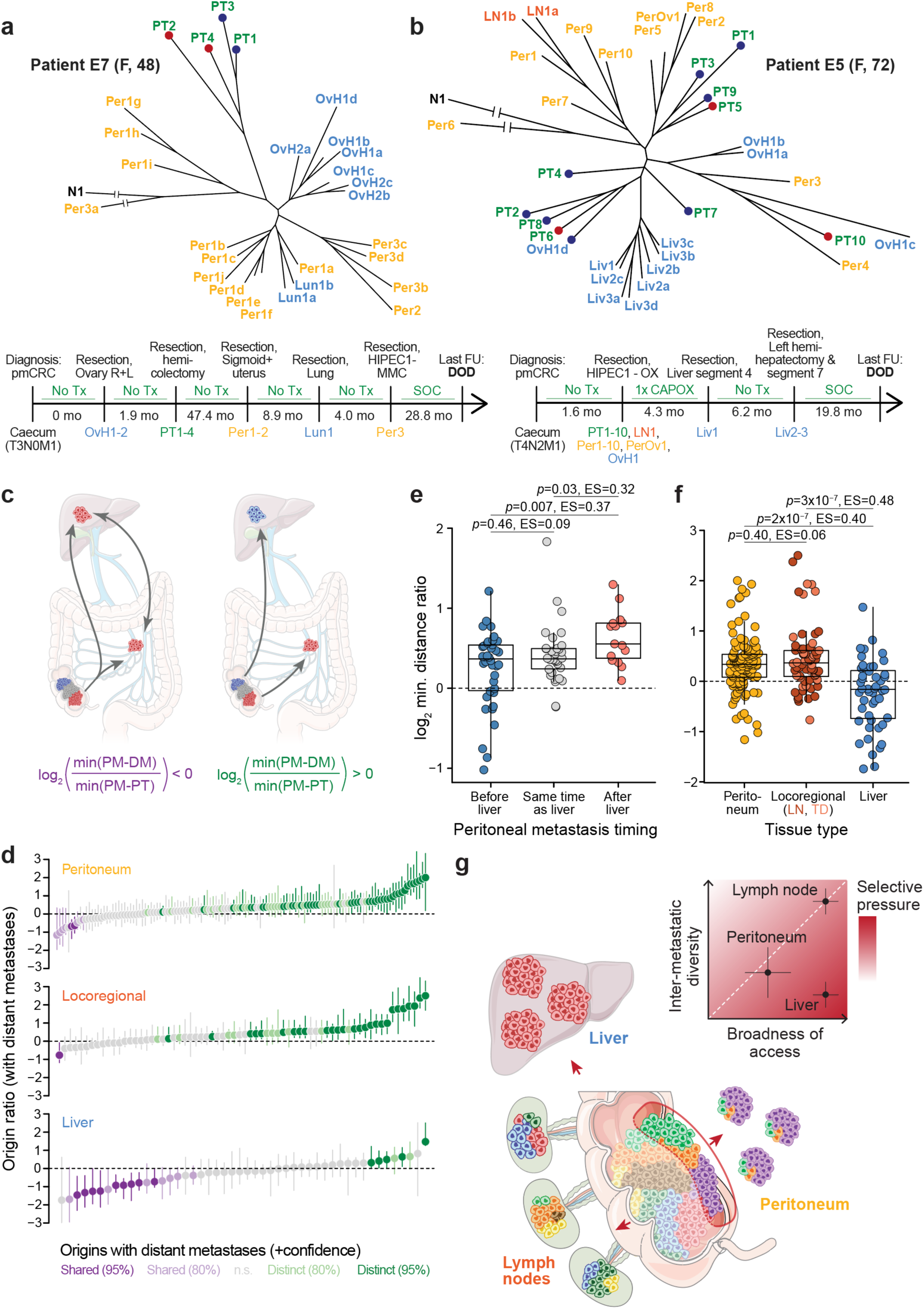
Peritoneal and distant metastases typically have distinct evolutionary origins. **a,** Phylogenetic tree and clinical timeline for patient E7. OvH, Ovarian metastasis of suspected hematogenous origin by pathological examination (tumor growth within the parenchyma but not on the ovarian surface). **b,** Phylogenetic tree and clinical timeline for patient E5. **c,** Schematic depicting two possibilities for the lineage relationship between peritoneal (PM) and distant metastases (DM). *Left*, peritoneal and distant metastases have a common subclonal origin. This could mean that they are both seeded from the same primary tumor lineage, or that they gave rise to each other. In this case, the origin ratio, defined as 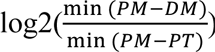, is expected to be smaller than 0. *Right*, peritoneal and distant metastases have distinct origins in the primary tumor. In this case, the origin ratio is expected to be larger than 0. **d,** Origin ratios for each peritoneal, locoregional, and liver metastases. Specifically, the origin ratio for peritoneal metastases is as described above, for locoregional metastases it is 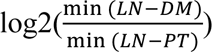 and for liver metastases it is 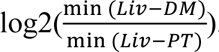. Bootstrapped 95% confidence intervals based on 1,000 iterations of randomly resampled polyguanine markers. Confidence values for origin classifications are based on the upper bound of 80% or 95% confidence intervals. **e,** Peritoneal-liver metastasis origin ratios 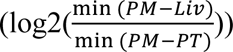 for peritoneal metastases arising before, synchronously with, or after liver metastases. *Left*, peritoneal metastases arising at least 3 months prior to the earliest detected liver metastasis; *center*, within 3 months of the earliest liver metastasis; *right*, at least 3 months after the earliest liver metastasis. **f,** Direct comparison of origin ratios for peritoneal, locoregional and liver lesions metastases (same data as in (d)). Locoregional point color differentiates lymph node metastases and tumor deposits. *P*-values and effect sizes based on independent pairwise comparisons using two-sided Wilcoxon rank sum tests. **g,** Summary schematic. Genetic diversity from the primary tumor (colored cells) is transferred most efficiently to locoregional metastases, less efficiently to peritoneal metastases, and least efficiently to liver metastases, resulting in decreasing inter-metastatic diversity across these host sites (inset). The broadness of tumor cell access to the relevant migration routes is high for cells undergoing lymphatic or hematogenous metastasis, and more restricted for cells undergoing peritoneal metastasis. By jointly evaluating broadness of access and inter-metastatic diversity, we deduce that selective pressures are highest during liver metastasis (see discussion for details).

To identify larger trends in the relationship between peritoneal and distant metastases across the whole cohort, we recorded the genetic distance of each peritoneal metastasis to its closest distant metastasis (*min*(*PM* − *DM*)), as well as its closest primary tumor sample (*min*(*PM* − *PT*), **Fig. 5c**). The log_2_ ratio of these two values (hereafter “origin ratio” for simplicity) is negative whenever a peritoneal metastasis associates closely with a distant metastasis while being further removed from the primary tumor. This can either indicate that both the peritoneal and the distant metastasis arose from related lineages in the primary tumor, or that they seeded each other. Conversely, the ratio is positive if the peritoneal metastasis is more closely related to the primary tumor than to any distant metastasis, indicating that the two lesions likely arose from distinct primary tumor subclones.

Examining origin ratios across *n*=129 peritoneal lesions that arose in patients who also had distant metastases, we noticed a clear skewing towards positive values, meaning that most peritoneal metastases were more closely related to the primary tumor than to a distant metastasis (**Fig. 5d**). These results were reminiscent of a prior study in which we had investigated the lineage relationships between lymph node and distant metastases and found that in most (65%) colorectal cancer patients, lymph node metastases did not share a subclonal origin with distant metastases but originated directly in the primary tumor^13^. Other studies subsequently reached the same conclusions^31^. Notably, we found that the small subset of peritoneal lesions that were more closely related to a distant metastasis than to the primary tumor (purple bars in the top panel of **Fig. 5d**) was enriched for patients who were diagnosed with liver metastases at the same time or after their peritoneal metastasis resection (**Fig. 5e**). That was the same patient group that had exhibited uncharacteristically close associations between the deep-invading edge and liver metastases in **Fig. 4g**, providing further support for the idea that peritoneal-to-liver seeding is relatively common when peritoneal metastases precede or coincide with liver metastases.

Calculating origin ratios for locoregional metastases (lymph nodes and tumor deposits) in the peritoneal metastasis cohort, we found a distribution that resembled our previous findings (76% of locoregional metastases associated more closely with the primary tumor while 24% resembled distant metastases more closely). In a direct comparison, we found no difference between the origin ratio distributions of locoregional and peritoneal metastases (**Fig. 5f**), suggesting that both metastasis types had the same close relationship to the primary tumor and the same comparatively weak link to distant metastases. Importantly, when we calculated origin ratios for liver metastases, we observed a much more pronounced skewing towards negative values (**Fig. 5d**), indicating that liver metastases typically arose from the same subclonal lineage as other distant metastases^14^. The difference between the origin ratios of peritoneal/lymph node metastases and liver metastases was highly significant (**Fig. 5f**). A multiple linear regression which included the ratio between the number of sampled primary tumor areas and number of distant metastases (a potential confounder) as an additional independent variable also recovered highly significant differences between origin ratios for liver metastases on the one hand and peritoneal and locoregional metastases on the other hand (peritoneum vs. liver *p*=1.4×10^-5^; locoregional vs. liver *p*=3×10^-6^; ratio between the number of sampled primary tumor areas and number of distant metastases *p*=0.55). We conclude that just like locoregional metastases, peritoneal metastases typically originate in the primary tumor and mostly do not share a common evolutionary origin with distant metastases.

## Discussion

In this study, we have traced the origins of hundreds of colorectal cancer peritoneal metastases, locating them on a detailed genetic map with the primary tumor and other locoregional and distant metastases as landmarks for comparison. Our exhaustive sampling scheme allowed us to make several novel observations. First, we were able to assess metastatic randomness in peritoneal metastases in direct comparison with locoregional and liver metastases. We observed that peritoneal metastases showed intermediate levels of inter-metastatic heterogeneity: they were less diverse than locoregional metastases but more diverse than liver metastases. Second, we found that peritoneal – but not lymph node or liver – metastases tended to associate with lineages (or subclones) that were located on the deep-invading edge of the primary tumor. This result is consistent with decades of clinical experience showing that patients whose primary tumor has breached the serosal lining face a much-increased risk of peritoneal metastasis. Finally, we found that peritoneal metastases typically have distinct evolutionary origins from metastases in distant organs like the liver or lungs. This surprising discovery can perhaps be explained by their preferential seeding from the deep-invading front, a predilection that is not shown by other types of metastases. Patients who develop liver metastases following peritoneal metastases appear to be an exception: we often observe a shared subclonal origin of metastatic lesions in these cases, and we suspect that cancer cells may reach the liver from the peritoneum in these cases.

What can these observations in their totality teach us about the cells that seed peritoneal metastases in humans? Is a specialized trait required for peritoneal metastasis formation? The data suggest that the peritoneal-metastatic process exerts *less* stringent selective pressures on disseminating tumor cells than the liver-metastatic process, resulting in a greater variety of subclones that can successfully complete it. Hence, there is a relative lack of requirement for specialized traits, at least in comparison with the liver. Specifically, the following line of argument can be considered: The degree of metastatic diversity in a host organ is determined by i) the broadness of tumor cell access to that organ and by ii) the strength of selective pressures encountered by tumor cells during dissemination and colonization. If access is broad and plentiful, and selective pressures are low, many subclones will successfully complete the process and metastases will exhibit high inter- and intra-lesion diversity. Lymph node metastasis appears to be a paradigm for this type of metastasis. Lymphatic vessels are typically broadly distributed in and around colorectal cancers^32^, giving most lineages access to this migration route (although lymphatic transport can sometimes be hampered by the physical forces generated during tumor growth^33^). In line with this observation, lymph node metastases are equally likely to originate in luminal or deep tumor areas in our data. Once cells enter the lymphatics, shear and oxidative stresses are low and beneficial lipids that protect cells from ferroptosis abound^16^. The broadness of access to lymphatics in combination with low selective pressures in the environment result in high levels of genetic diversity in and among lymph node metastases. Tumor cells require few specialized traits to succeed.

Liver metastasis seems to present a much greater challenge. Access to the relevant migration route – the blood vessel – is likely to be just as broad as for the lymphatics, as all viable tumor areas must have at least a minimal vascular supply. Liver metastases in patients who do not (yet) have peritoneal metastases do not preferentially associate with luminal or deep areas. However, since liver metastases show striking levels of genetic homogeneity, both within and among lesions, we must conclude that the migration and/or colonization process exerts strong selective pressures onto cancer cells, resulting in a severe “thinning” of lineages that can survive the process. Again, this is consistent with many functional studies that have shown the systemic circulation to be a highly challenging environment for cancer cells^17^. Liver metastases probably arise from select lineages that have acquired a trait that confers a selective advantage in this context.

Extending the same logic to the peritoneum, we can cautiously conclude a few points: First, access to the peritoneal cavity is likely to be more restricted than access to the lymphatics or the blood. Only subclones that have completely breached the colon wall can easily reach the peritoneum and seed metastases – for all other subclones, the journey is far and the seeding rate probably negligible. This unique spatial restriction should dramatically restrict the diversity of peritoneal metastases. However, metastatic diversity in the peritoneum trumps the liver, although access to the organ is much less broad. We believe that this can only be explained by relatively lax selective pressures: many of the cells that do have access to the peritoneal cavity will be able to grow there, resulting in higher diversity despite restricted access (summary cartoon in **Fig. 5g**). It is possible that cancer cells benefit from being able to reach the peritoneum without having to expose themselves to the systemic circulation – a particularly harsh environment as discussed above.

If cells that form peritoneal metastases indeed have not yet evolved to withstand the severe selective pressures that are exerted upon them during systemic dissemination, we may perhaps look hopefully toward future treatment options for patients with this disease. One possible reason that even a single resectable liver metastasis portends a poorer prognosis than many (equally resectable) lymph node metastases might be that the occurrence of the liver metastasis signals the presence of a very dangerous lineage. This species can travel in the blood stream and grow even in unfriendly environments, darkening the outlook for the patient. If peritoneal-metastatic cells – much like locoregional metastases – indeed represent a less evolved precursor of this lethal cell type, long-term disease control could perhaps still be achieved if we had more effective local management tools. For example, if more complete resections could be achieved through advanced intra-operative imaging, outcomes might improve further. Finally, the distinct genetic properties of peritoneal metastases, which in some respects recapitulate features of locoregional metastases, raise the question whether peritoneal metastases should be staged separately from metastases in other distant organs.

## Methods

### Tissue samples

#### Peritoneal metastasis patient cohort

Tumor specimens from two institutions (The St. Antonius Hospital, Nieuwegein, the Netherlands and Massachusetts General Hospital, Boston, USA) were included in this study after approval from each hospital’s institutional review board (MGH IRB and Medical Research Ethics Committees United in Nieuwegein), and in accordance with the Declaration of Helsinki. Since polyguanine fingerprinting is a very limited interrogation of non-coding genomic regions, it is performed under a waiver of consent. We searched internal databases for patients who had undergone resection of at least one peritoneal metastasis from colorectal cancer. Search results were then subset for cases with an available primary tumor. We reviewed hematoxylin & eosin (H&E)-stained tissue sections for candidate patients to identify those that had tumors of sufficient size and purity. For example, cases were excluded from consideration if they had very small peritoneal metastases consisting of only a small number of scattered tumor cells, or if primary tumors exhibited extensive necrosis and treatment effects. In this manner, we identified 20 suitable patients from St. Antonius (E1-E20) and 4 patients from Massachusetts General Hospital (C146, C154, C157, C158). In all these cases, both primary tumor and metastases consisted of high-quality tumor tissue. This cohort of 24 patients was supplemented with two patients (C38 and C89) with colorectal cancer and peritoneal metastases from previously published studies^13,14^.

#### Additional tumor samples

To compare the genetic properties of peritoneal metastases to distant lesions in other organs, we included in various analyses throughout the paper polyguanine fingerprints from n=25 additional patients with metastatic colorectal cancer to the liver from previously published studies^13,14^. As in Reiter et al.^14^, we also included 5 patients with colorectal cancer and liver metastases whose phylogenies were generated from whole-exome sequencing data (patients CRC1-5)^34^. These 30 retrospective patients contributed only to analyses involving liver metastases. The 5 whole-exome phylogenies CRC1-5 were only included in analyses involving root diversity scores (RDS), which can be calculated independently of the underlying data type if a phylogenetic tree is available. Finally, 3 previously unpublished patients with metastatic colorectal cancer are described in this study which are not part of the peritoneal metastasis cohort. These patients (C159, C161, C186) either had no peritoneal metastases (C186), or only had peritoneal metastasis samples recovered at autopsy (C159 and C186), a biologically distinct scenario which we did not want to mix with the main cohort which consisted exclusively of surgical resection specimens. These patients were only used to compare polyguanine vs. low-pass whole genome sequencing-derived phylogenies in **Fig. 1c-e** and **Supplementary Fig. 2**; their data are not included in any other analyses.

#### Metastasis characteristics and naming conventions

Anatomically distinct metastases were given distinct numbers, in line with our previously used naming convention^13,14^. For example, Per1 and Per2 represent two spatially separated peritoneal lesions. Multiple samples from the same metastatic lesion are additionally labeled with letters, e.g. Per1a and Per1b. Metastases were classified as synchronous if they were present at the time of primary tumor resection, and metachronous if they were resected more than 3 months after that. This study distinguishes between lymph node metastases (locoregional metastases that retain clearly identifiable lymphoid tissue) and tumor deposits (locoregional metastases that show no evidence of lymphoid tissue). These categories are thought to be potentially biologically distinct^35^. For patients C38 and C89 who were included from previous studies, histological images were no longer available, and we could not make this distinction. Therefore, in analyses that distinguish between lymph node metastases and tumor deposits (as opposed to treating them as one category), locoregional metastases from C38 and C89 were excluded. In analyses of treatment effects, we generally considered systemic chemotherapy separately from hyperthermic intraperitoneal chemotherapy (HIPEC), as the latter is a local intervention. We refer to distant metastases in the liver, lungs, ovaries or distant lymph nodes as “untreated” if they were never exposed to systemic chemotherapy. Peritoneal metastases, however, were only considered untreated if they had experienced neither chemotherapy nor HIPEC. Patient E13’s liver metastasis (Liv1) arose after treatment with HIPEC only, we therefore retain it in the untreated category.

#### Histology and tissue processing

A board-certified gastrointestinal pathologist (J.K.L) reviewed H&E slides or images and carefully annotated tumor regions. Bulky tumor was sampled with 1.5-2 mm core biopsies. For smaller regions of interest, 8 µm tissue sections were macrodissected under the microscope. Tissues were deparaffinized with xylene and digested with proteinase K overnight. DNA was extracted with phenol-chloroform and precipitated with ethanol and sodium acetate, as previously described^13,14^.

### Polyguanine fingerprinting

Detailed descriptions of the primer sequences and PCR protocol for amplification of polyguanine repeats have been previously published^13,14^. We genotyped between 18 and 45 markers per patient (mean = 31.3 markers/patient). Genotypes for each marker and tumor sample were acquired in triplicate for a total of 73,859 individual PCRs that contributed to the data set. Peak information for each sample, polyguanine marker, and replicate was extracted from GeneMapper 4.0 and used as input to a previously described automated analysis pipeline^13^. We will briefly explain the main analysis steps here, but a more detailed description can be found in the original publication. In a first pre-processing step, we removed all reactions that did not show robust amplification, i.e. where fragment fluorescence intensity was less than 10% of the average intensity for that polyguanine marker and patient. After intensity filtering, we proceeded to examine the three PCR replicates for each marker and sample and chose the best (“representative”) replicate for further analysis, as described in detail in Naxerova et al.^13^. Briefly, for each sample, we calculated the Jensen-Shannon distance (JSD, the square root of the Jensen-Shannon divergence^36^) among all replicate pairs. Two replicates were classified as identical if their JSD was less than 0.11. Among the pairs of replicates that were classified as identical, we chose the pair with the lowest JSD and selected the replicate with the higher intensity to be the representative replicate for the sample. If no two replicates were classified as identical, either the sample was excluded, or the marker was excluded across all samples of a given subject. Because our phylogenetic reconstruction method is intolerant to missing data, we removed polyguanine markers from a patient’s data set if they failed to amplify for too many samples, or we removed tumor samples if they had too many missing genotypes (e.g. due to suboptimal DNA quality). Balancing the dual goals of retaining as many markers and as many tumor samples as possible for each patient, we chose to remove between 1-15% of polyguanine marker genotypes per patient to obtain a valid genotype across all samples. We ran this entire pipeline once at the beginning of the analysis stage for all samples and then executed an impurity exclusion step to remove samples that did not meet minimum purity standards. Due to the introduction of the purity-robust angular distance method, we were able to slightly relax the stringency of impurity exclusion in comparison with previous studies^13,14^. The adjusted impurity exclusion approach was identical to the published version^13^, except that for each sample, we now fitted a linear regression, with the ‘calibrator distances to normal reference’ as response and ‘sample distances to normal reference’ as explanatory variables. If the slope was smaller than 0.35, the sample was excluded from the analysis. After impurity exclusion, the entire pipeline was re-run on the remaining set of sufficiently pure samples to generate the final data set on which all subsequent analyses were based. This data set is freely available at https://github.com/agorelick/peritoneal_metastasis. The automated pipeline is implemented as an installable R library and can be downloaded at https://github.com/agorelick/polyG.

### Phylogenetic reconstruction and analysis of evolutionary trees

#### Phylogenetic reconstruction from polyguanine fingerprints

For each patient, we constructed a distance matrix containing pairwise, purity-robust angular distances between samples. More details on this distance measure are provided in the accompanying **Supplementary Note**. In brief, the stutter distribution derived from each polyguanine locus in each sample was simplified to its mean fragment length^15^. Each tumor sample is thus represented by a vector of length *n*, with *n* being the number of measured polyguanine markers. We subtracted from this vector the mean lengths measured in the normal tissue control, thus creating the “polyguanine fingerprint” which encodes the sample’s somatic mutations across all polyguanine loci. Next, all polyguanine fingerprints were normalized to unit length. The angular distance between two unit length polyguanine fingerprints x and y is then defined as arccos (*x* · *y*). Phylogenies were then constructed from the angular distance matrices using the neighbor-joining method (implemented in the *ape* R package^37^). The normal germline sample was attached post-hoc to each patient’s phylogeny by connecting it to the last internal node created by the neighbor-joining algorithm. Please note that the branch *length* leading to the normal tissue control sample has no meaning in this context (to highlight this, we introduce a break in the branch leading to the normal tissue samples in all phylogenies). Phylogenies were annotated with confidence values based on 1,000 bootstrap replicates constructed by randomly re-sampling polyguanine markers with replacement.

#### Pre-processing for inter-lesion heterogeneity analyses

In some cases, we sampled individual metastatic lesions multiple times. These samples were useful to assess intra-metastatic heterogeneity. However, for all analyses of inter-metastatic diversity (described below), we had to choose *one* sample to represent the metastasis and thus make it comparable to other metastases that were only sampled once. Therefore, for all analyses involving pairwise inter-lesion angular distances or root diversity scores, we generated a “collapsed” version of each patient’s phylogenetic tree. In the collapsed tree, each distinct metastatic lesion is represented by only one sample. For simplicity, we chose the sample that received the letter “a” in the initial sample processing workflow, resulting in a random choice from a biological point of view.

#### Metastasis-specific root diversity score (RDS)

To assess the phylogenetic diversity of multiple, anatomically distinct metastases with respect to the multi-region sampled primary tumor, we calculated the root diversity score (RDS), defined as the probability that at least *l* out of *m* metastases form a monophyletic clade in a tree with *n* = *k* + *m* tumor samples^14^. The RDS reflects the probability that a tree with an equal or more extreme clustering of metastases occurs by chance alone. In this study, we calculated “metastasis-specific” RDS values by constructing collapsed phylogenetic trees that only contain primary tumor samples and samples from a metastasis group of interest (peritoneal, lymph node, tumor deposit, liver). This approach avoids that clustering of different metastasis subtypes (e.g. clustering of both lymph node and liver metastases away from the primary tumor) inflates RDSs. Metastasis-specific RDS values thus quantify the degree of separation *between the primary tumor and a specific metastasis type of interest* (unencumbered by the position *other* metastases on the phylogenetic tree). R code that can be used to calculate RDS values for any phylogenetic tree has been packaged into an installable R library available at https://github.com/agorelick/rds.

### Analysis of somatic copy number alterations (SCNAs)

#### Low-pass whole genome sequencing

Low-pass whole genome sequencing (lpWGS, ∼1x) libraries were prepared from genomic DNA using the NEBNext Ultra DNA Library Prep Kit for Illumina. Quality of raw sequencing output was verified using FASTQC^38^ (v0.12.1) and sequencing reads were aligned to the human reference genome (version humanG1Kv37) with the BWA-MEM algorithm^39^ (v0.7.15) with soft-clipping enabled.

#### SCNA-based trees from lpWGS data

lpWGS-based genomic copy number profiles were generated using the QDNASeq^40^ and ACE^41^ R packages to obtain copy number estimates in 1Mb genomic bins. For each patient, the QDNAseq R package was used to generate read counts in 1Mb bins for each BAM file, based on reference human genome hs37d5. To correct read counts for bins’ variable GC content and mappability, we estimated correction factors using *estimateCorrection* function with default parameters after first excluding sex chromosomes, as well as blacklisted and low mappability (< 25) regions. To retain sex chromosomes in the corrected read counts (X and Y are excluded in this output by default by QDNAseq), as per the authors’ instructions, the filter on sex chromosomes was then removed and the corrections were applied to bins from all chromosomes. The corrected read counts were then normalized, outliers were smoothed, bin counts were segmented, and copy number segments were normalized, following instructions in the QDNAseq documentation with default parameters. For each sample, the ACE package was then used to obtain purity and ploidy-corrected total copy number.

Briefly, given an overall average ploidy value (usually 2 or 4), ACE exploits the notion that clonal copy number alterations create segments with integer-valued total copy number to find purity values that maximize the number of segments with integer values (indicating that the tumor purity and ploidy have correctly been adjusted for). For a given patient, we manually reviewed and fine-tuned the purity/ploidy fits from ACE for each sample to best-align the likely-clonal segments, such that they had the same integer copy number values in each sample (while also preserving their overall similarity across all tumor samples). The 1Mb copy number bins were then corrected for these purity/ploidy values, and the resulting total copy number profiles were used to calculate Euclidean distances between each pair of samples. Finally, the neighbor-joining algorithm was used to construct phylogenetic trees for each patient’s copy number data. Confidence values for copy number-based phylogenetic trees were generated based on 1,000 bootstrap replicates constructed by randomly sampling entire chromosomes with replacement from the 1Mb bin data.

#### Tests for concordance between polyguanine and lpWGS-based phylogenetic data

The similarity between two phylogenetic trees representing the same set of samples was quantified using the quartet similarity (implemented in the *Quartet* R library, which employs the tqDist software^42^). Given two unrooted, bifurcating trees of size *n* with the same set of samples, the quartet similarity is defined as the fraction of all possible 4-sample subtrees that are common between them. To test whether two phylogenetic trees were more similar than expected by chance, we used a permutation-based approach, comparing the true quartet similarity between two trees to a null distribution generated by randomly permuting the tip labels on one of the two trees 10,000 times. A one-sided *p*-value was calculated as the fraction of random permutations with a quartet similarity at least as great as the similarity observed, with a pseudocount of 1 added to both the numerator and denominator. A similar permutation-based approach was used to test the similarity between two distance matrices for the same set of samples. Here, the rows and columns of the matrices were both ordered identically, and Spearman’s correlation coefficient was calculated between the values in each matrix’s upper triangle (excluding the diagonal). To compare this to a null distribution, we randomly permuted the row and column labels in one of the matrices 10,000 times, each time reordering its rows and columns to match the non-permuted matrix and calculating a new Spearman correlation. A one-sided *p*-value for the significance of the matrices’ similarity was calculated as described above.

### Clonal evolution inference for patient C157

#### Whole exome sequencing data processing

Whole exome sequencing libraries were prepared from genomic DNA using the Human Core Exome kit by Twist Bioscience and sequenced to an average depth of 184x. Quality of raw sequencing output was verified using FASTQC^38^, and adapter sequences were trimmed using Cutadapt^43^ (v4.1). Sequencing reads were pre-processed following the GATK Best Practices workflow^44^ for variant discovery. Briefly, sequencing reads were aligned to the human reference genome (version humanG1Kv37) with the BWA-MEM algorithm (v0.7.15). Duplicate reads were removed using GATK (v4.1.9.0) *MarkDuplicatesSpark*. Base scores were recalibrated using GATK *BaseRecalibrator* and *ApplyBQSR* using all polymorphic sites from dbSNP (build 151) as the list of known sites for exclusion. After pre-processing sequencing data, somatic mutations were called using *Mutect2* in multi-sample mode using a matched normal sample, a panel of normal provided by GATK, and germline allele frequencies provided by gnomAD^45^ using default arguments. To reduce potential false positives due to formalin fixation, orientation bias artifact priors were learned using *LearnReadOrientationModel* and the resulting variants were then filtered using *FilterMutectCalls.* MAF files were created from VCF using vcf2maf (v1.6.21). Mutations were annotated for known/predicted oncogenic effects using OncoKB^46^ (v3.3, accessed Dec 5, 2023) and considered potentially oncogenic if they were annotated by OncoKB as “Oncogenic” or “Likely Oncogenic”, or if the variant was classified as “Loss-of-function” or “Likely Loss-of-function”. The FACETS software^47^ (v0.6.2) was used to estimate purity and ploidy and generate allele-specific copy number profiles for each sample. Mutations were then annotated with estimated cancer cell fractions (CCFs) as previously described^48,49^. Mutations with any detected mutant reads were considered subclonal if the CCF’s 95% confidence interval’s upper bound was less than 0.9, and clonal otherwise. This process resulted in 3,507 mutation calls. We next applied a series of filters to exclude mutations which were likely induced by formalin. In addition to only using variants with “PASS” filter status based on GATK *FilterMutectCalls*, we removed SNVs that met all of the following 3 criteria: (1) had the dominant formalin-associated mutation signature^50^ C>T (or G>A); (2) were not likely to be oncogenic based on OncoKB annotation; and (3) were detected in only 1 sample, subclonally. We additionally removed any variant for which we suspected the possibility of false negative calls for any sample. Briefly, for every mutation, we checked if any sample that had no reads supporting the mutant allele may have been a false negative due to insufficient power to detect the mutation (e.g., due to insufficient sequencing depth at that position). To this end, we calculated the probability of detecting at least one read with the mutant allele in each sample, given its tumor purity and read depth at that position, and assuming the mutation had a CCF of 20% and arose on one mutant copy. If this probability was below 90% in any sample with 0 mutant reads, we considered this mutation call a potential false negative and therefore removed this mutation from all samples. After applying these filters, 2,385 protein-coding missense, nonsense, splice-site, translation start site mutations, silent mutations, or small in-frame and frame-shift insertions and deletions remained for downstream phylogenetic analyses.

#### Clone detection and phylogeny inference

Mutations that were clonal in at least one sample were clustered into subclones using PyClone-vi^51^ and their phylogenetic relationships were inferred using Pairtree^52^ and Orchard^53^. Here, we conservatively excluded sample LN2b, as this sample had an unexpectedly large number of variants compared to other samples (3.2x the median number of clonal mutations, 2.7x the median number of variants overall) and was therefore suspicious for being enriched for false positives. (However, LN2b was retained in the overall WES mutation phylogeny). After removing variants that according to Pairtree violate the infinite sites assumption using r*emovegarbage*, we input the remaining variants into PyClone-vi, which was run using a maximum of 20 clones. PyClone-vi identified 7 mutation clusters, which after manual curation to split up clusters with high CCF variance and remove four additional variants with ambiguous clustering resulted in 10 clusters (*i.e.*, clones) comprising 86 mutations. Finally, Orchard was used infer a phylogeny from the 10 clones and estimate the proportions of each clone in each tumor sample.

### Simulated effects of chemotherapy on metastasis heterogeneity

To illustrate the effects of systemic chemotherapy on inter-lesion heterogeneity, we simulated 100 patients, each with four high-diversity “peritoneal” lesions and four low diversity “liver” lesions. All lesions initially consisted of 1 million cells that were randomly sampled from 5 subclones (A, B, C, D, E). Peritoneal lesions were created by sampling from each subclone with equal probability, translating into approximately 20% frequency for each clone. Liver lesions were created by sampling from clone A with probability 0.95 and from the remaining clones with probability 0.0125 each. Chemotherapy was then simulated as the random death of 80% of cells in each metastasis. The surviving cells regrew according to a discrete time birth-death process with birth rate 0.25 and death rate 0.24^14^ until the lesions contained 100 million cells. At this point, inter-lesion heterogeneities for peritoneal and liver lesions were quantified as the median pair-wise Euclidean distance between lesions of each type.

### Simulated angular distance between tumors with variable purities

To explore the effects of tumor purity on angular distance, we simulated a phylogeny with two ancestrally related tumor samples of perfect purity. Specifically, we first generated a germline genotype (“c0”) which consisted of 100 polyguanine markers of random lengths between 10-20 nucleotides. From this germline genotype, we constructed a genotype for the patient’s first cancer cell (“c1”): each marker mutated independently with a constant probability of 5×10^-4^ per generation; each mutation was either an insertion or a deletion of 1 basepair with equal probability; and 1,000 generations passed between the germline genotype and c1. Next, we simulated two tumor genotypes (“c2” and “c3”) that evolved from the ancestral c1 genotype over another 1,000 generations. The resulting genotypes c2 and c3 thus represented the genotypes of two 100% pure, ancestrally related tumors. We then numerically admixed the pure tumor genotypes c2 and c3 each with c0 (the germline genotype) at various proportions to create impure tumor genotypes (“t1” and “t2”). The resulting genotypes c0, t1, and t2 represented the *idealized* genotypes of the patient’s germline and tumor samples. Technical noise affects measurements from any assay and has a greater effect when the magnitude of the signal is small. We therefore simulate *measured* genotypes by adding random noise to each marker length to represent technical error. As a surrogate measurement for technical noise, we used the precision in marker length measurements. Briefly, for each normal sample in our patient cohort, length distributions for each polyguanine marker were sampled with replacement 200 times, and the coefficient of deviation of their mean lengths (their standard deviation normalized by their mean) was recorded. Across all markers and patients, the median coefficient of variation in marker length was 0.002. Random noise was then simulated with a normal distribution of mean 0 and standard deviation equal to the true length of the marker multiplied by 0.002, and added to every marker length in c0, t1, and t2. Finally, the angular distance between t1 and t2 was calculated (see **Supplementary Note**) using the measured c0 as the genotype of the patient’s matched normal sample. This angular distance was then compared to an “optimal” value assuming no technical error and 100% purity of each tumor (**Supplementary Note Fig. 1**).

### Animal experiments

#### Patient-derived organoids and organoid culture

All human experiments were approved by the ethical committee of University Medical Center Utrecht (UMCU). Written informed consent from the donors for research use of tissue in this study was obtained prior to acquisition of the specimen. Tumor patient-derived organoids (PDOs) were embedded in ice-cold Matrigel® (Corning) mixed with a CRC culture medium in a 3:1 ratio. The medium contained advanced DMEM/F12 medium (Invitrogen), HEPES buffer (Lonza, 10 mM), penicillin/streptomycin (Gibco, 50 U/ml), GlutaMAX (Gibco, 2 mM), R-spondin-conditioned medium (20%), Noggin-conditioned medium (10%), B27 (Thermo/Life Technologies, 1×), nicotinamide (Sigma-Aldrich, 10 mM), N-acetylcysteine (Sigma-Aldrich, 1.25 mM), A83-01 (Tocris, 500 nM), EGF (Invitrogen/Life Technologies, 50 ng/ml) and SB202190 (Gentaur, 10 μM). For passaging, the PDOs were dissociated with TrypLE Express (Gibco) for 5–10 min at 37°C and re-plated in a pre-warmed 6-well plate. Rho-associated kinase (ROCK) inhibitor Y-27632 (Tocris, 10 μM) was added to culture medium upon plating for two days.

#### Multicolor Marking

The PDOs were simultaneously transduced with three RGB constructs (each encoding for a color) according to a previously published protocol^28^. The following lentiviral gene ontology (LeGO) vectors were used: LeGO-C2 (27339), LeGO-V2 (27340), and LeGO-Cer2 (27338) (Addgene). Briefly, lentiviral production was performed by a calcium phosphate transfection protocol in human embryonic kidney 293T cells using the transfer plasmid (15 µg), pMD2.G (12259, 7.5 µg) and psPAX2 (12260, 7.5 µg). The following day, medium was replaced by advanced DMEM/F12 medium (Invitrogen) supplemented with HEPES buffer (Lonza, 10 mM), penicillin/streptomycin (Gibco, 50 U/ml), and GlutaMAX (Gibco, 2 mM). The next day, 50,000 single cells of PDOs were resuspended in the virus medium (which was filtered through 0.45 μm polyethersulfone filter), supplemented with Polybrene (Sigma-Aldrich, 8 µg/ml), N-acetylcysteine (Sigma-Aldrich, 1.25 mM) and ROCK-inhibitor Y-27632 (Sigma-Aldrich, 10 μM), and incubated overnight 37°C, 5% (vol/vol) CO_2_ on non-adherent plates (ultra-low attachment surface, Sigma-Aldrich). After 24h incubation, cells were washed twice in PBS (Sigma-Aldrich) and cultured as described above. The PDOs were sorted based on YeGr2-A (mCherry), blue 1-A (Venus), and Violet1-A (Cerulean) expression at least two passages after transduction on a Fluorescence Activated Cell Sorting (FACS) Aria II (BD Biosciences) machine.

#### Orthotopic caecum-implantation model

This study was approved by Utrecht University’s Animal Welfare Body, the Animal Ethics Committee and licensed by the Central Authority for Scientific Procedures on Animals. To evaluate the spontaneous metastatic capacity of the tumor PDOs, we made use of the murine orthotopic caecum-implantation model^54^. In summary, one day before implantation, RGB-PDOs were dissociated into single cells and 2.5×10^5^ cells were plated in 10 μL drops of neutralized Rat Tail High Concentrated Type I Collagen (Corning). The PDOs were allowed to recover overnight at 37°C, 5% (vol/vol) CO_2_. NOD.Cg-Prkdc^SCID^ Il2rg^tm1Wjl^/SzJ (NSG) male mice, between 8-10 weeks of age, were treated with a subcutaneous dose of analgesic Carprofen (5 mg/kg, Rimadyl^TM^) 30 min before surgery and were subsequently sedated by isoflurane inhalation anesthesia [2% (vol/vol) isoflurane/O2 mixture]. The caecum was exteriorized through a midline abdominal incision and a single collagen drop containing the RGB-PDOs was micro-surgically transplanted in the cecal submucosa. Carprofen was given 24h post-surgery. The endpoint for all animals reached after five weeks. Caecum, liver and peritoneum were harvested for further analysis.

#### Tissue processing and image analysis

Mouse organs containing multicolor, endogenously fluorescent primary tumors and metastases were fixed using 4%-paraformaldehyde in PBS solution overnight at 4°C followed by tissue preservation in a 20% sucrose solution for 12h at 4°C. Samples were cut into 4 μm-thick frozen tissue sections and covered with ProLong Gold Antifade Mountant (Thermo Fisher Scientific). For high-resolution imaging, sections were scanned with an LSM 700 confocal microscope (Carl Zeiss) using an EC Plan-Neofluar 10x/0.30 or 20x/0.50 M27 objective. Fluorescence images were preprocessed by downsampling to 1.25 µm/pixel and Gaussian filtering (scikit-image, scipy-ndimage, Python). For all images and image channels, auto-fluorescence of the background tissue was individually contrast corrected by intensity clipping at appropriate, manually chosen, thresholds (**Supplementary Table 6**) followed by rescaling pixel values to 0.6% oversaturation. Lesions were manually highlighted as regions of interest (Labkit, Fiji) and images were split into separate images for each lesion. Pixel values were filtered for pixels in which the sum of RGB values was at least 50 and converted to the HSV scale. For each image, Hue values were binned into 10 equally sized bins and counted before calculating the Simpson diversity index (*vegan*, R) (approach adapted from Coffey et al.^55^). The average diversity was calculated over a total of 10 repetitions, each time shifting the bin positions. Differences in SDI values between caecum, peritoneum, and liver regions were assessed using a Kruskal-Wallis test followed by Dunn’s test with Holm’s correction for multiple hypothesis testing.

### Statistical analyses and figures

All statistical analyses were performed with R (version 4.1.1) unless otherwise noted. In boxplots comparing two groups, statistical differences were tested with unpaired two-sided Wilcoxon rank sum tests and effect sizes were calculated using the *rstatix* R library. For boxplots with pairwise-comparisons of three or more groups, a Kruskal-Wallace test was used followed by a post-hoc Dunn’s test with Holm’s correction for multiple hypothesis testing as implemented in the *rstatix* library, and Wilcoxon rank sum test effect sizes are provided for each comparison. *p*-values from independent tests were corrected for multiple hypothesis testing with the Benjamini-Hochberg method and reported as *q*-values method where appropriate. In all boxplots, boxes show the median value with lower and upper hinges corresponding to the 25th and 75th percentile values, respectively. Boxplot whiskers extend from the lower/upper hinges to the smallest/largest values no further away than 1.5 times the inter-quartile range from the hinge. Figures were generated with either base-R or the *ggplot2* library. Phylogenetic trees were visualized using the *ggtree* library^56^.

## Data availability

Easily machine readable raw polyguanine marker data are available at https://github.com/agorelick/peritoneal_metastasis. Processed data files used as input for analyses in this study, including angular distance matrices, lpWGS binned read counts and full clinical data are available at the same repository. As for all our previous studies, the raw data has also been submitted to Data Dryad and is ready to download for reviewers under: https://datadryad.org/stash/share/jYnd55JoQ5F6gRYI0LwHd3RZ83TQv7W3jFGN-B0bts0. Deposition of raw lpWGS and whole exome sequencing data to dbGAP has been initiated; the full data set will be available at the time of publication (preliminary accession ID: phs003722.v1.p1).

## Code availability

Code and instructions to regenerate all data-based figures is available at https://github.com/agorelick/peritoneal_metastasis. Angular distance matrices can be regenerated from marker files by running the complete polyguanine PCR assay data-processing pipeline, which is available at https://github.com/agorelick/polyG.

## Acknowledgements

This work was supported by funding from the NIH/NCI (R37CA225655, R01CA279054, R01CA26928 to K.N.), an American Association for Cancer Research NextGen Grant for Transformative Cancer Research, an Emerging Leader Award from the Mark Foundation for Cancer Research, and the St. Antonius Research fund. Alexander Gorelick is the MacMillan Family Foundation Awardee of the Life Sciences Research Foudnation.

