## Supplementary Information for "A unique interplay of access and selection shapes peritoneal metastasis evolution in colorectal cancer"

Emma CE Wassenaar, Alexander N Gorelick, Wei-Ting Hung, David M Cheek, Emre Kucukkose, I-Hsiu Lee, Martin Blohmer, Sebastian Degner, Peter Giunta, Rene MJ Wiezer, Mihaela G Raicu, Inge Ubink, Sjoerd J Klaasen, Nico Lansu, Emma V. Watson, Ryan B. Corcoran, Genevieve Boland, Gad Getz, Dejan Juric, Geert JPL Kops, Jochen K Lennerz, Djamila Boerma, Onno Kranenburg, Kamila Naxerova

**Table of Contents**

Supplementary Figures

| | |
|---|---|
| Supplementary Fig. 1 | Correlation between primary tumor size and number of sampled regions. |
| Supplementary Fig. 2 | Comparison of SCNA- and polyguanine fingerprint-based phylogenies for patients C146, C154, C157, C159, C161, C186 |
| Supplementary Fig. 3 | Clinical timeline, mutation heatmap, phylogeny for all patients |
| Supplementary Fig. 4 | Clonal evolution in patient C157 |
| Supplementary Fig. 5 | Polyguanine fingerprints distinguish independent vs. clonally related primary tumors |
| Supplementary Fig. 6 | Intra-lesion heterogeneity visualized *in vivo* using optical barcoding |
| Supplementary Fig. 7 | Simulated effects of chemotherapy on inter-lesion heterogeneity |
| Supplementary Fig. 8 | Hematoxylin and eosin stains of primary tumor regions from patient E14 |
| Supplementary Fig. 9 | Association of lymph node metastases, tumor deposits and liver metastases with deep-invading vs. luminal primary tumor areas in each patient. |

Supplementary Note “Angular distance from Microsatellite Data”
Includes Supplementary Note Fig1. Simulated effects of tumor impurity on angular distance

Supplementary Tables

| | |
|---|---|
| Supplementary Table 1 | Clinical information for patients |
| Supplementary Table 2 | Detailed sample information |
| Supplementary Table 3 | Purity and ploidy for all lpWGS samples |
| Supplementary Table 4 | Treatment and timing of metastasis resection |
| Supplementary Table 5 | Metastasis-specific root diversity scores |
| Supplementary Table 6 | Channel threshold for mouse image analysis |

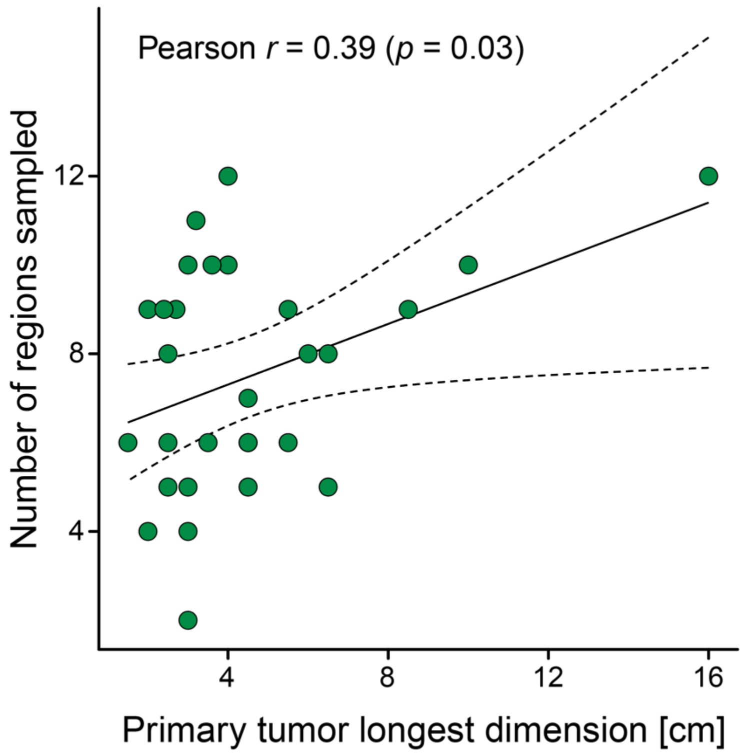


**Supplementary Figure 1. Correlation between primary tumor size and number of sampled regions.** X-axis, length of the longest dimension of each primary tumor in centimeters. Y-axis: number of regions sampled from that tumor. Pearson correlation coefficient and *p*-value as shown.

**Generalized legend for Supplementary Figure 2: Comparison of SCNA- and polyguanine fingerprint-based phylogenies for patients. a,** Heatmap of segmented total copy number in each sample. Black dots indicate segments with non-integer copy number consistent with subclonal SCNAs. Segments with copy number greater than or equal to 5 are visualized in the same color. Gray regions indicate segments for which copy number could not be inferred. Left, neighbor-joining phylogenetic tree based on pairwise Euclidean distances (calculated from purity/ploidy-corrected total copy number in 1Mb bins). **b,** Side-by-side comparison of SCNA and polyguanine distance matrices. Right, statistical test to assess their similarity. Spearman's correlation coefficient between the matrices (red line) is compared to a null distribution generated from 10,000 random permutations of sample names in the SCNA matrix (histogram). **c,** Side-by-side comparison of SCNA and polyguanine trees. Right, permutation-based test to assess their similarity. Similarity is measured by quartet distance (red line) and compared to a null distribution based on 10,000 permutations of tip labels in the SCNA tree (histogram).

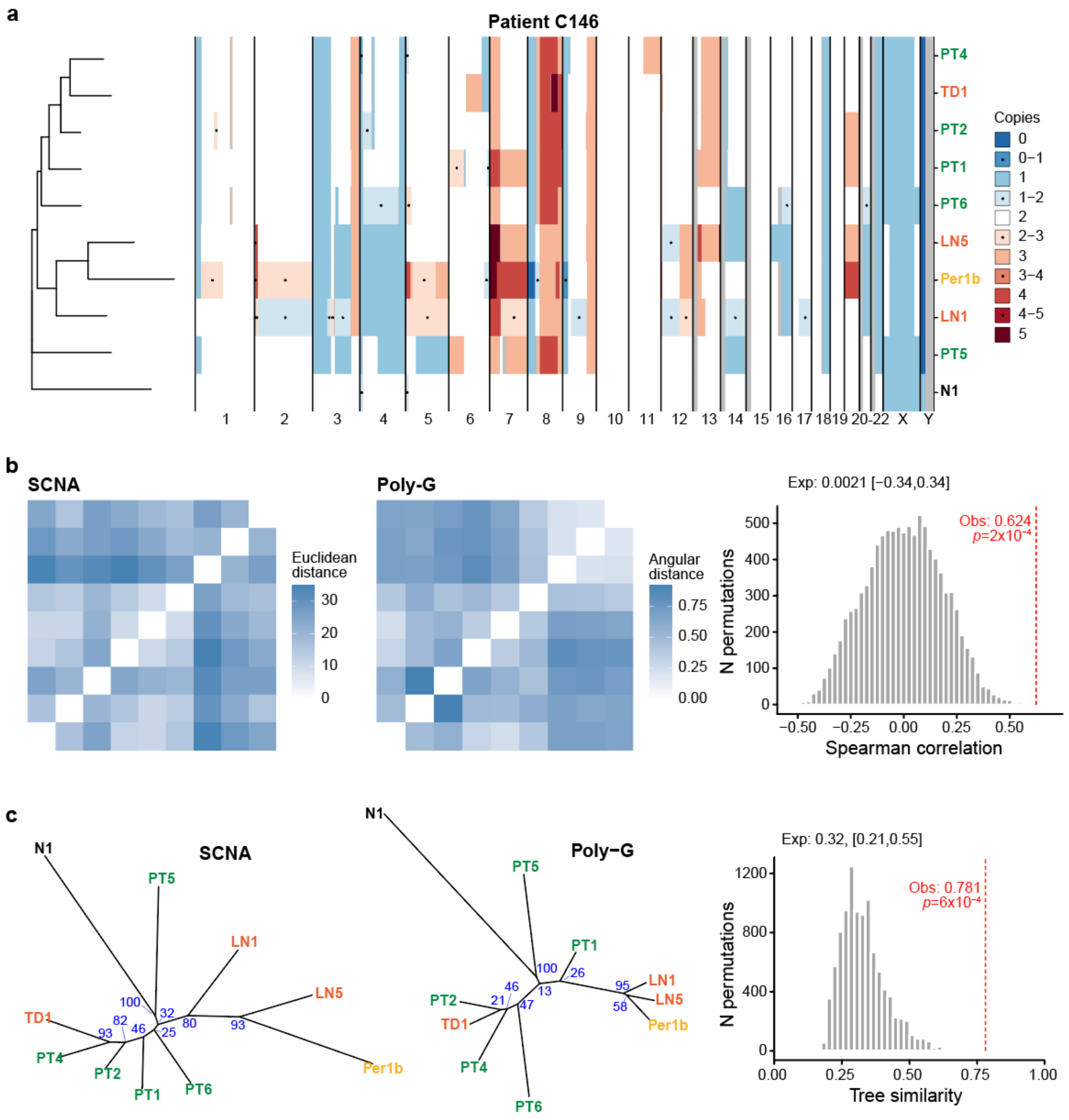


**Supplementary Figure 2.** Comparison of SCNA- and polyguanine-based phylogenies for patient C146.

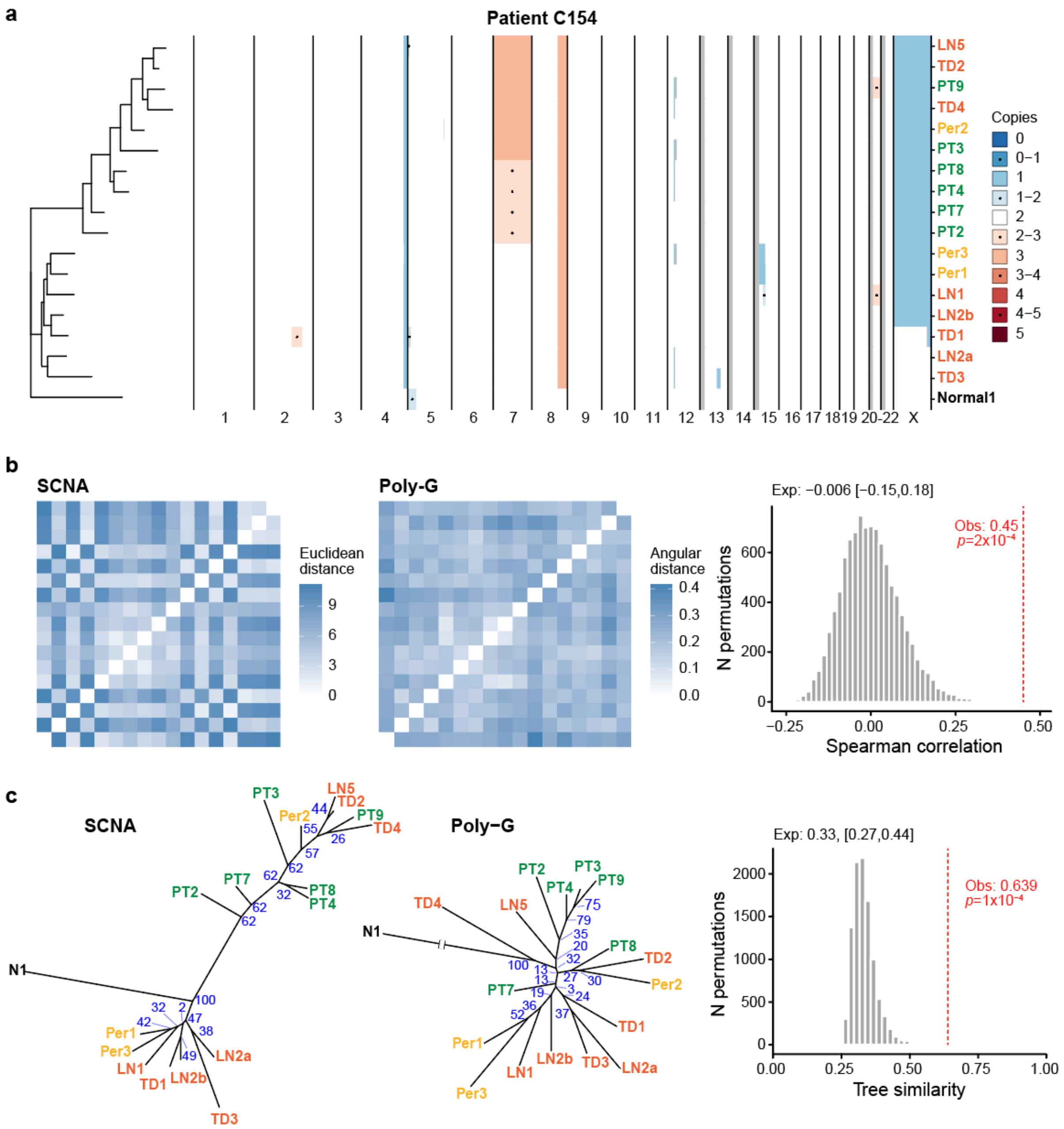


**Supplementary Figure 2.** Comparison of SCNA- and polyguanine-based phylogenies for patient C154.

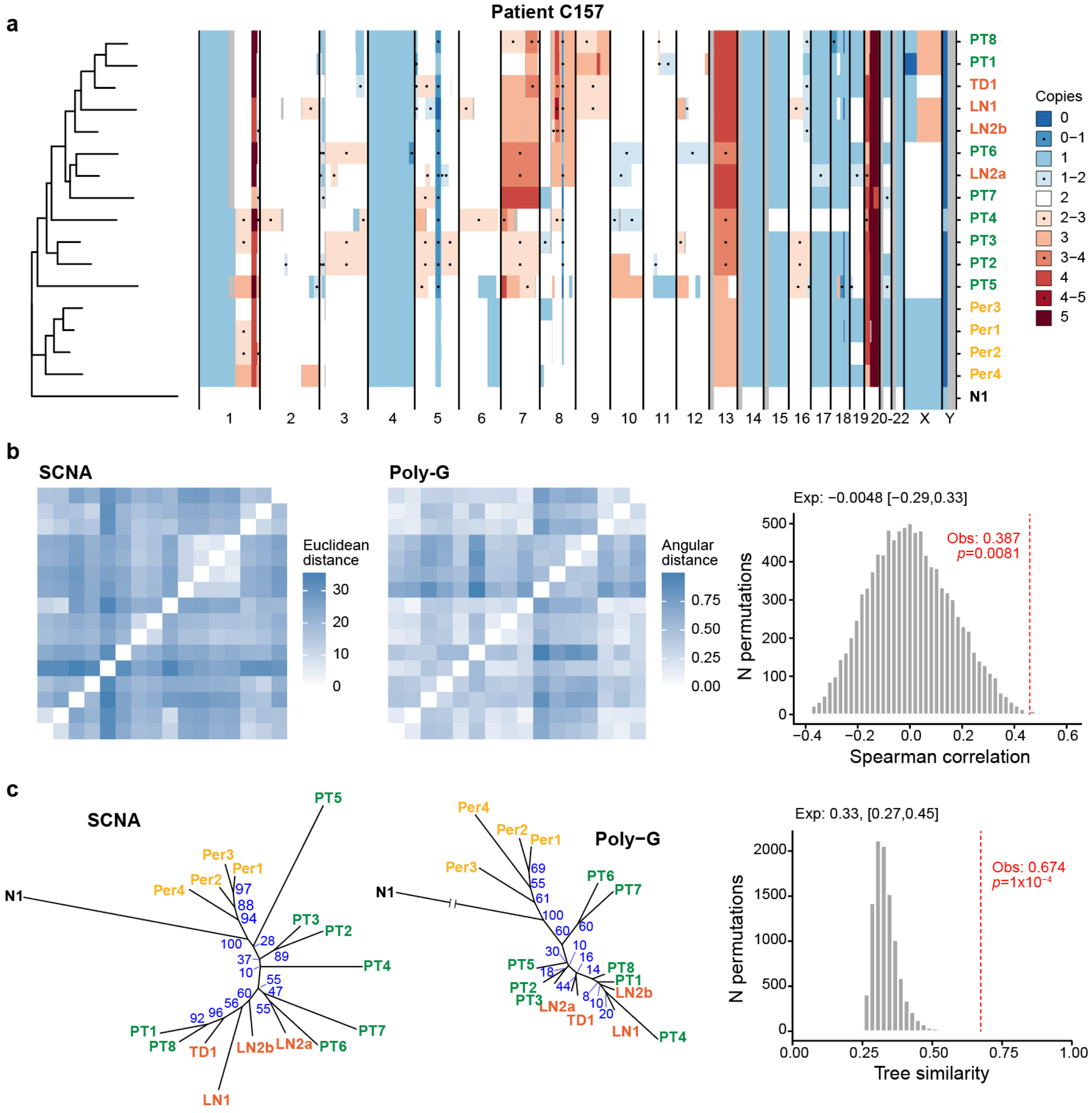


**Supplementary Figure 2.** Comparison of SCNA- and polyguanine-based phylogenies for patient C157.

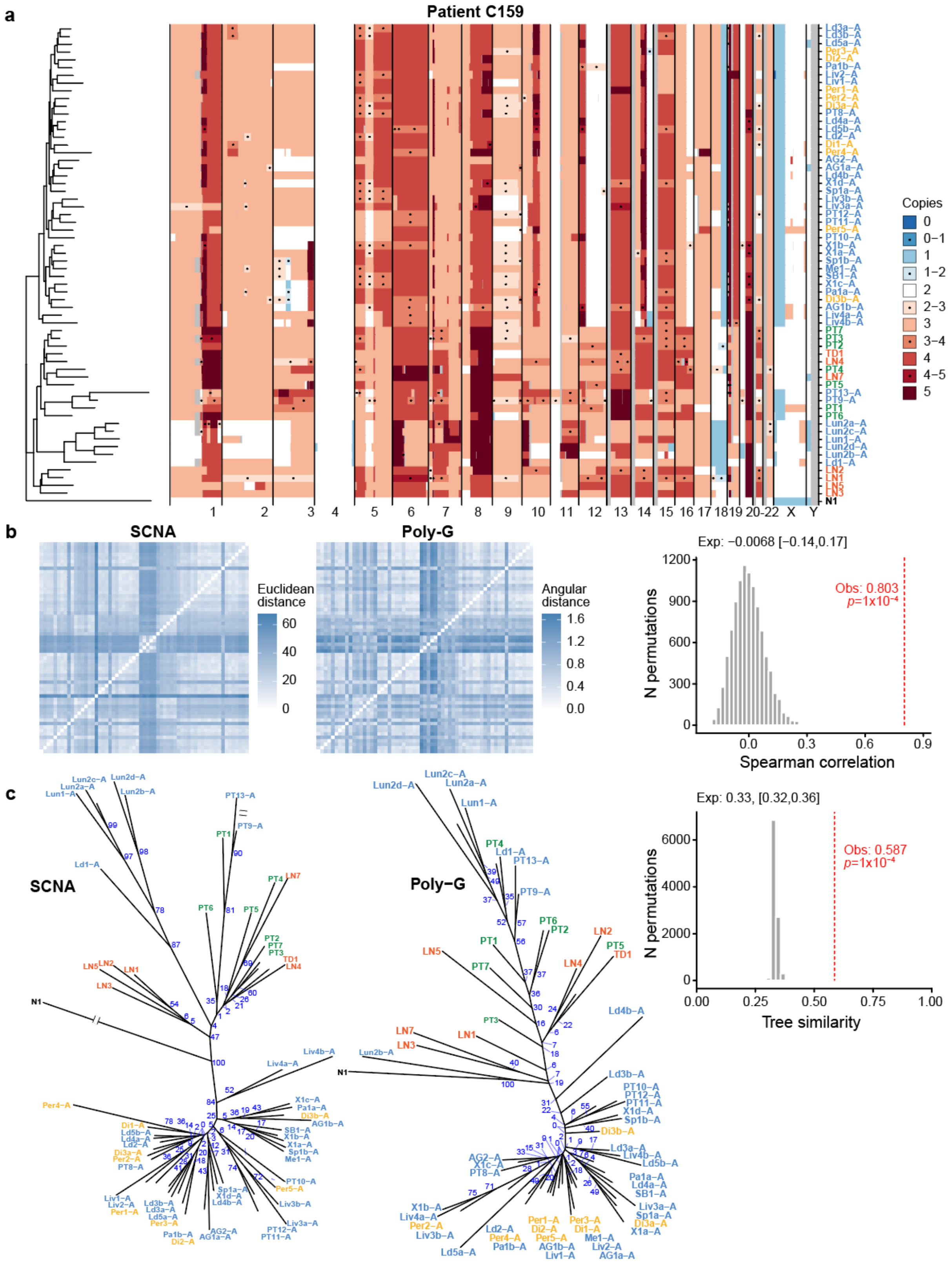


**Supplementary Figure 2.** Comparison of SCNA- and polyguanine-based phylogenies for patient C159.

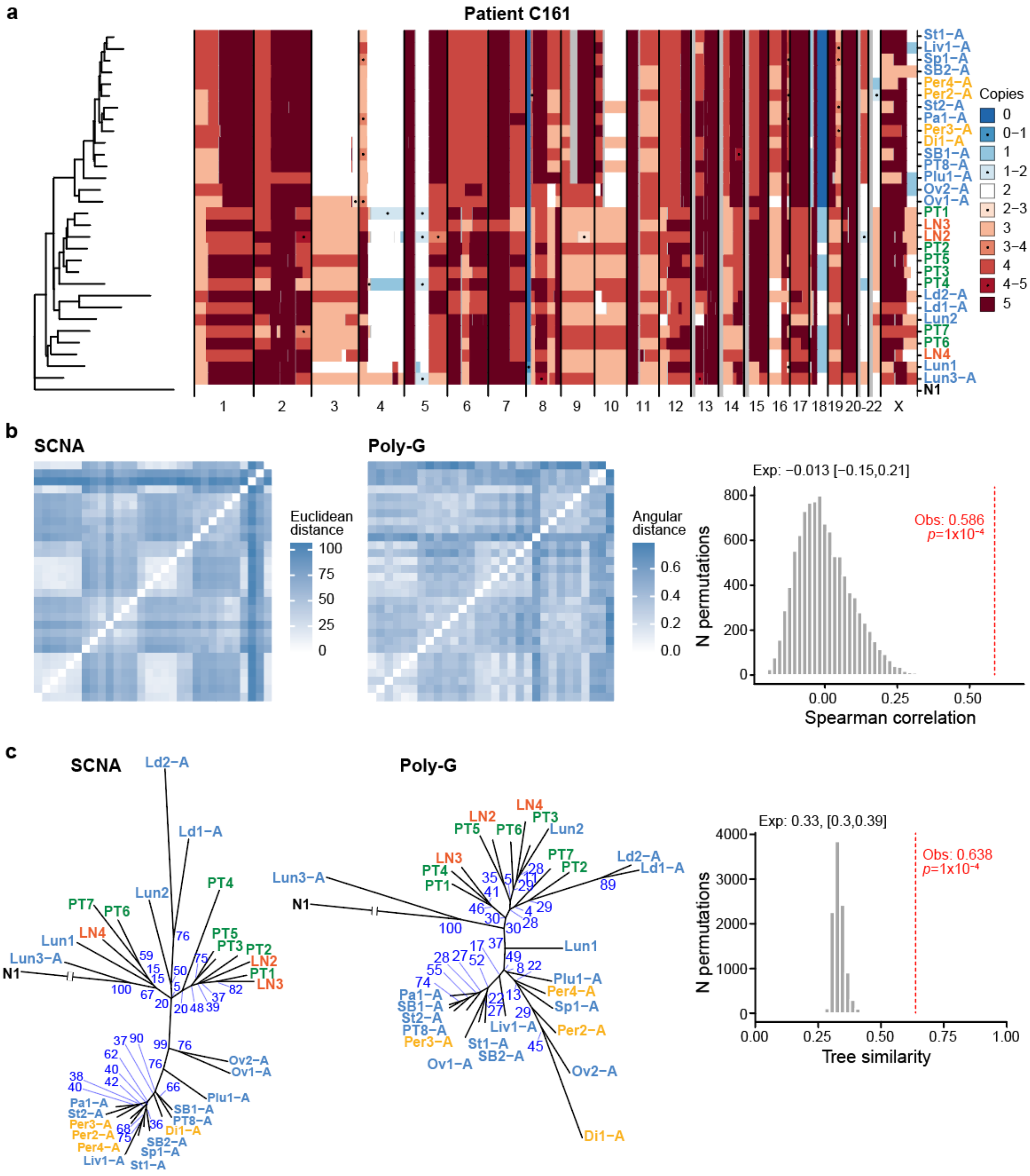


**Supplementary Figure 2.** Comparison of SCNA- and polyguanine-based phylogenies for patient C161.

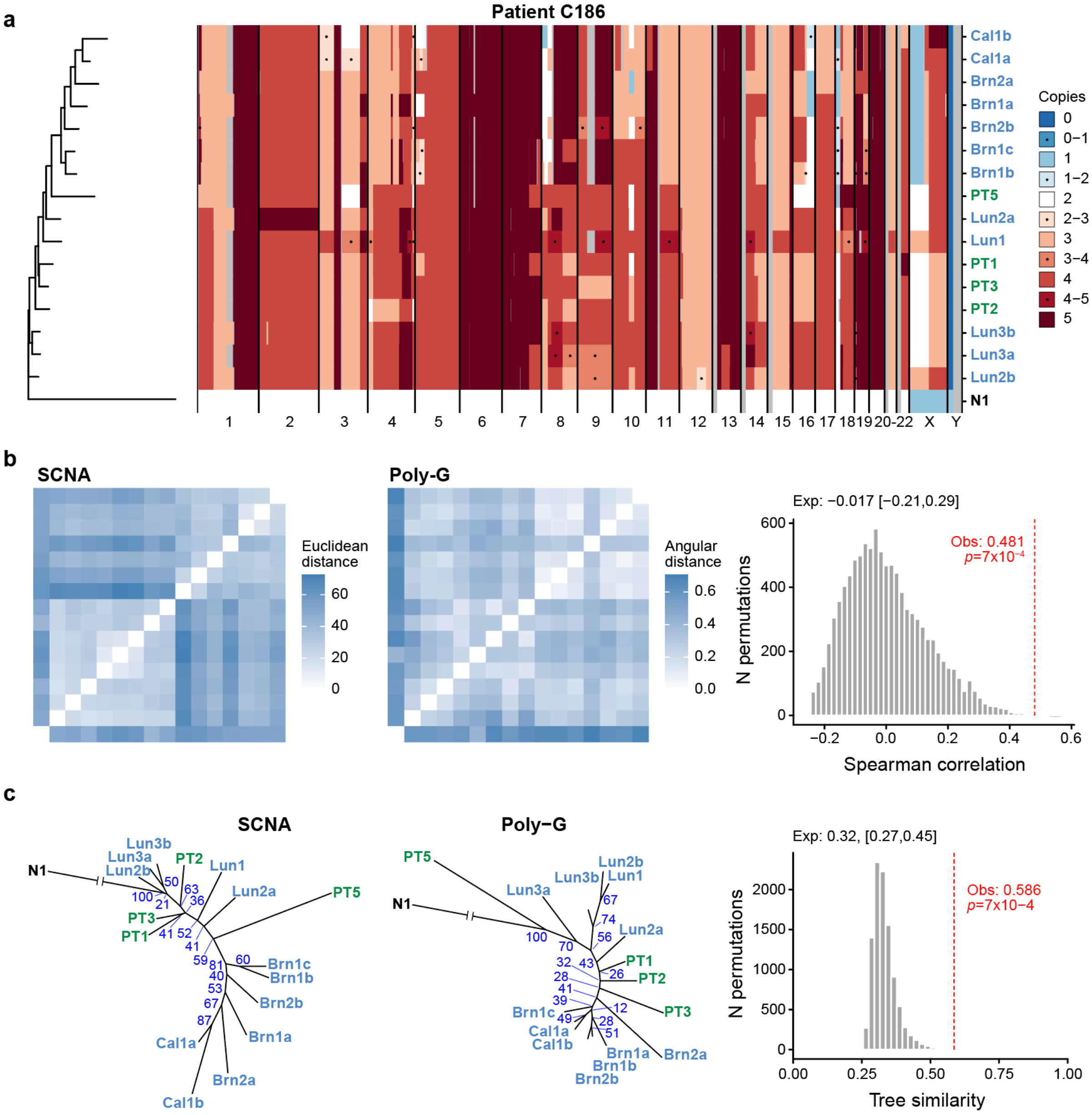


**Supplementary Figure 2.** Comparison of SCNA- and polyguanine-based phylogenies for patient C186.

**Generalized legend for Supplementary Figure 3: Additional information for each patient in the peritoneal metastasis cohort. a,** Clinical information and treatment timeline. **b,** Anatomical diagram indicating the primary tumor site and the locations and quantities of analyzed metastatic lesions. **c,** Representative histological images for select lesions. Circled regions indicate sampling locations. **d,** Visualization of polyguanine genotypes. The heatmap shows the mean lengths difference between a tumor sample and the normal reference for each marker. Rows are normalized to unit length. Green, deletions; purple; insertions. **e,** Angular distance-based phylogenetic tree, including confidence values (blue text) based on 1,000 bootstrap replicates. Primary tumor samples taken from deep-invading and luminal/mucosal regions are indicated with red and blue dots, respectively. Metastases are annotated according to their timing relative to the primary tumor surgery (open circles, synchronous; crosses, metachronous). **f**, Metastasis-specific RDS scores.

Abbreviations used in clinical timelines: OX = Oxaliplatin; MMC = Mitomycin C; CAPOX = Capecitabine + Oxaliplatin; FOLFOX = Leucovorin + Fluorouracil + Oxaliplatin; FOLFIRINOX = Leucovorin + Fluorouracil + Irinotecan + Oxaliplatin; RTx = radiotherapy; 5-FU = Fluorouracil; Cap mono = Cabecitabine monotherapy; SOC = standard of care; DOD = death of disease

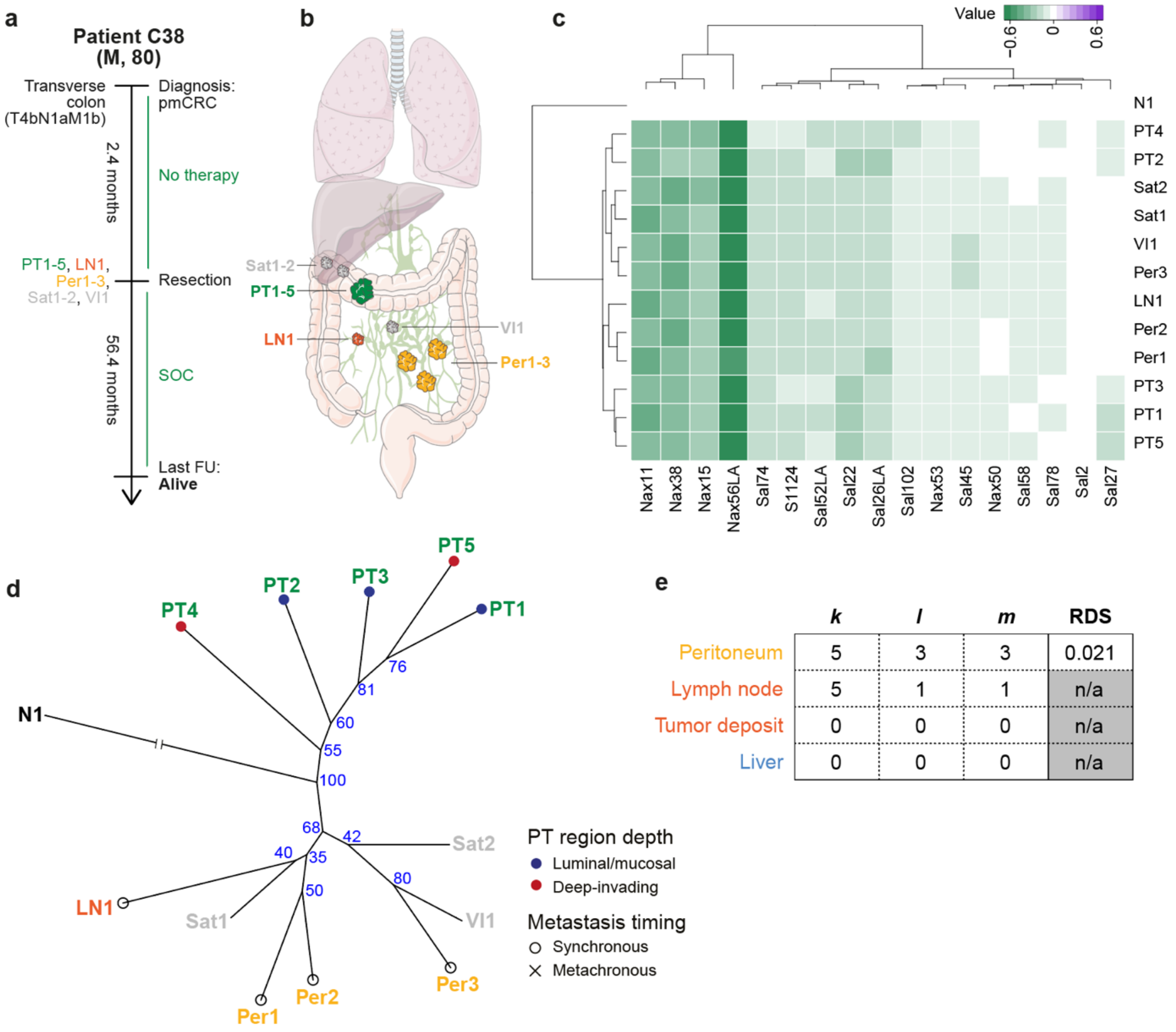


| | *k* | *l* | *m* | RDS |
|---|---|---|---|---|
| Peritoneum | 5 | 3 | 3 | 0.021 |
| Lymph node | 5 | 1 | 1 | n/a |
| Tumor deposit | 0 | 0 | 0 | n/a |
| Liver | 0 | 0 | 0 | n/a |

**Supplementary Figure 3.** Additional information for patient C38. (Please note, this figure does not contain histological images, therefore general legend caption d), e), f) applies to c), d), e) here).

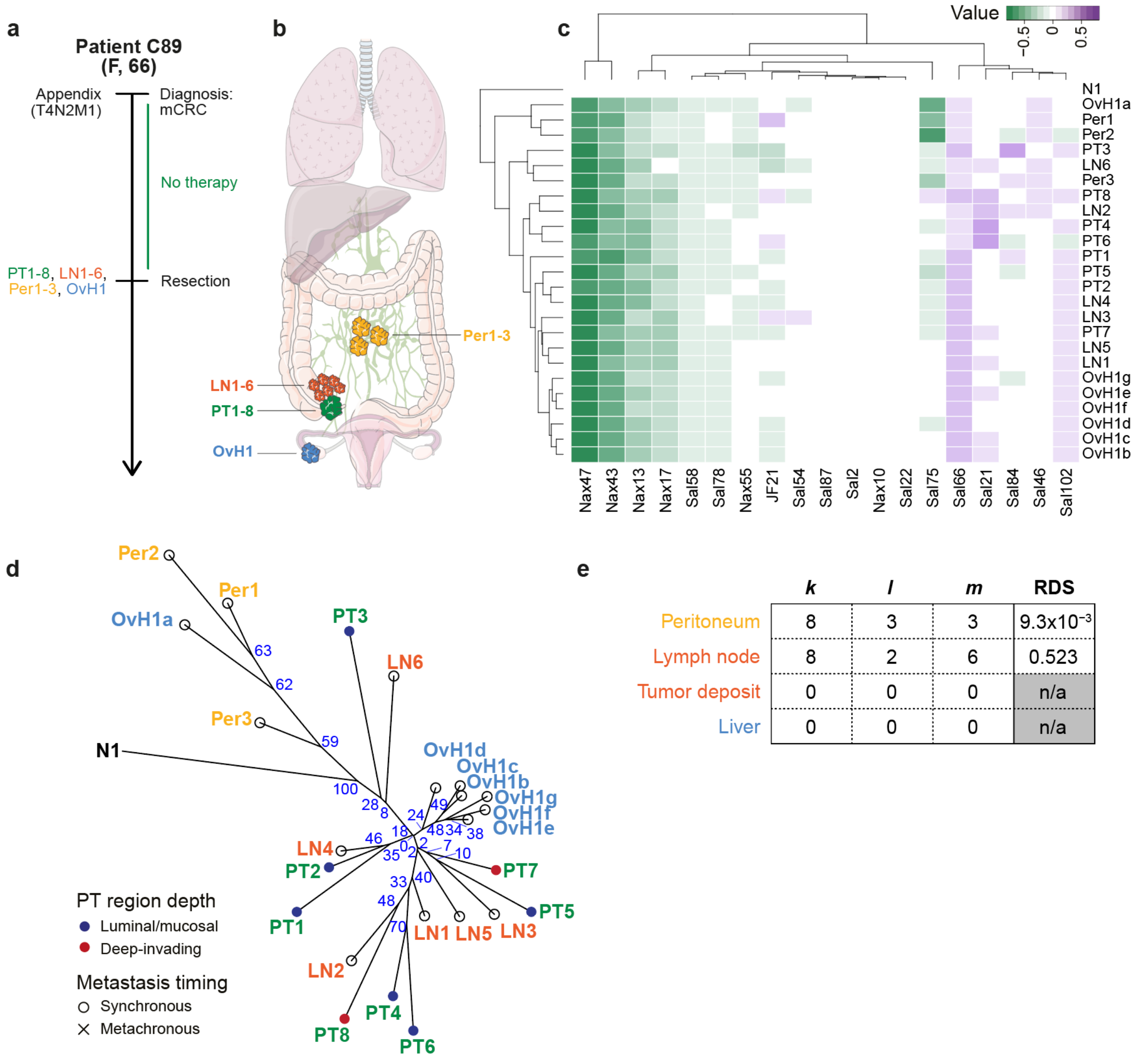


| | *k* | *l* | *m* | RDS |
|---|---|---|---|---|
| Peritoneum | 8 | 3 | 3 | $9.3 \times 10^{-3}$ |
| Lymph node | 8 | 2 | 6 | 0.523 |
| Tumor deposit | 0 | 0 | 0 | n/a |
| Liver | 0 | 0 | 0 | n/a |

**Supplementary Figure 3.** Additional information for patient C89. (Please note, this figure does not contain histological images, therefore general legend caption d), e), f) applies to c), d), e) here).

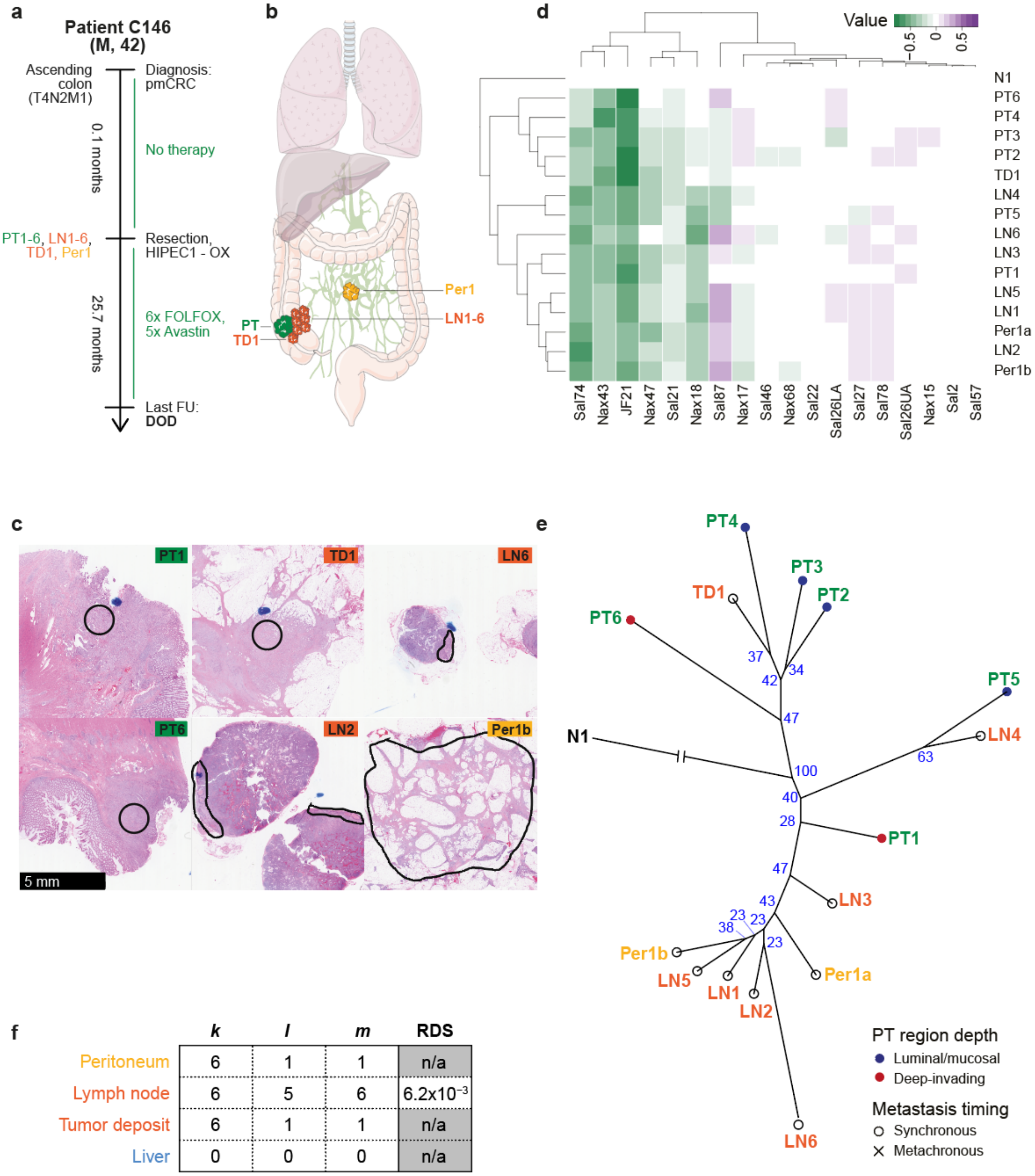


| | *k* | *l* | *m* | RDS |
|---|---|---|---|---|
| Peritoneum | 6 | 1 | 1 | n/a |
| Lymph node | 6 | 5 | 6 | $6.2\times10^{-3}$ |
| Tumor deposit | 6 | 1 | 1 | n/a |
| Liver | 0 | 0 | 0 | n/a |

**Supplementary Figure 3.** Additional information for patient C146.

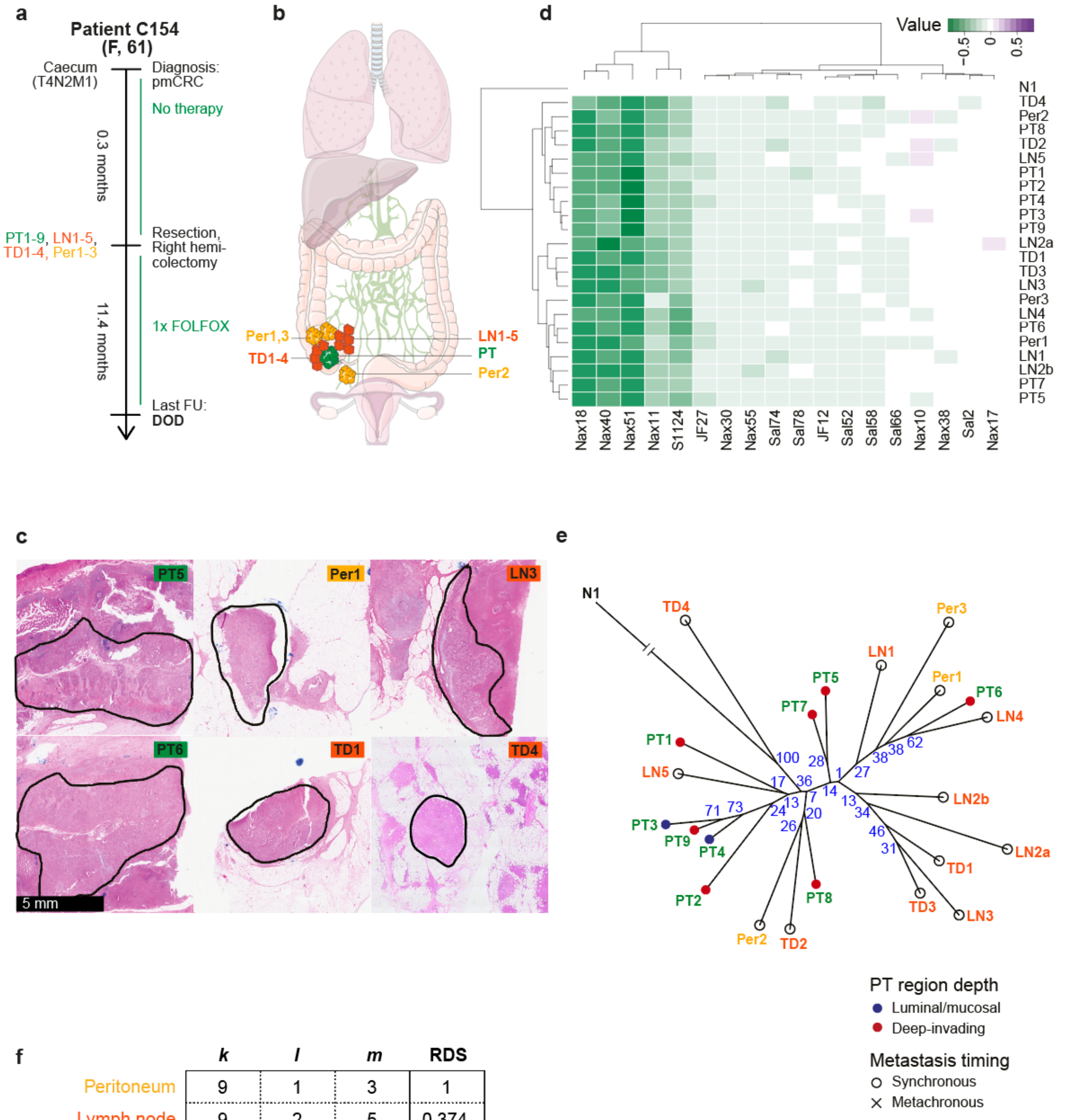


**f**

| | *k* | *l* | *m* | RDS |
|---|---|---|---|---|
| Peritoneum | 9 | 1 | 3 | 1 |
| Lymph node | 9 | 2 | 5 | 0.374 |
| Tumor deposit | 9 | 2 | 4 | 0.255 |
| Liver | 0 | 0 | 0 | n/a |

**Supplementary Figure 3.** Additional information for patient C154.

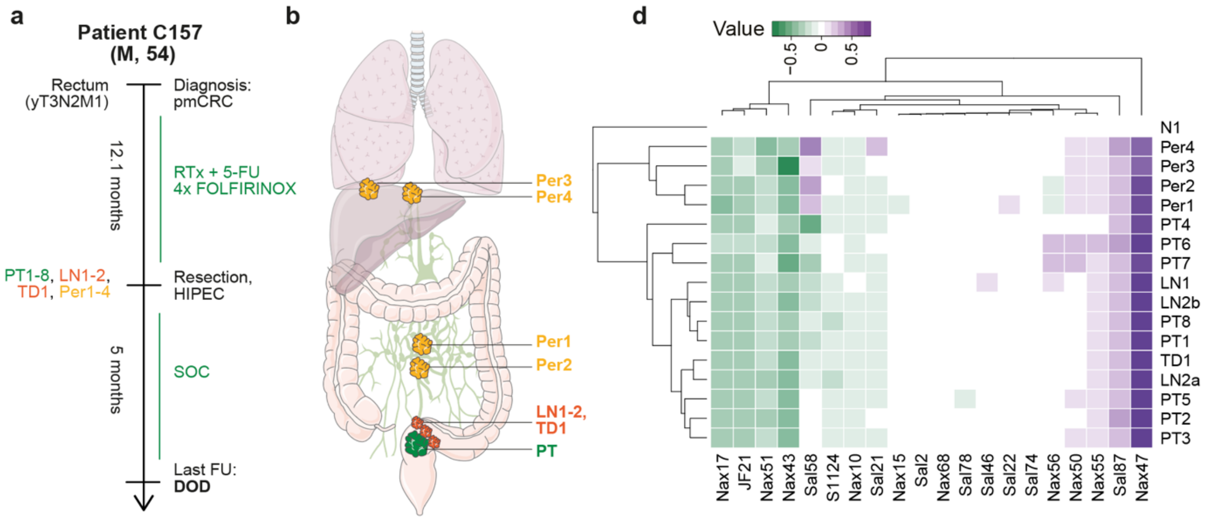


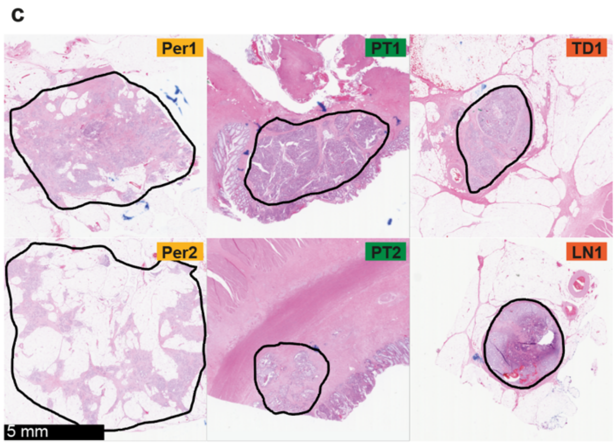


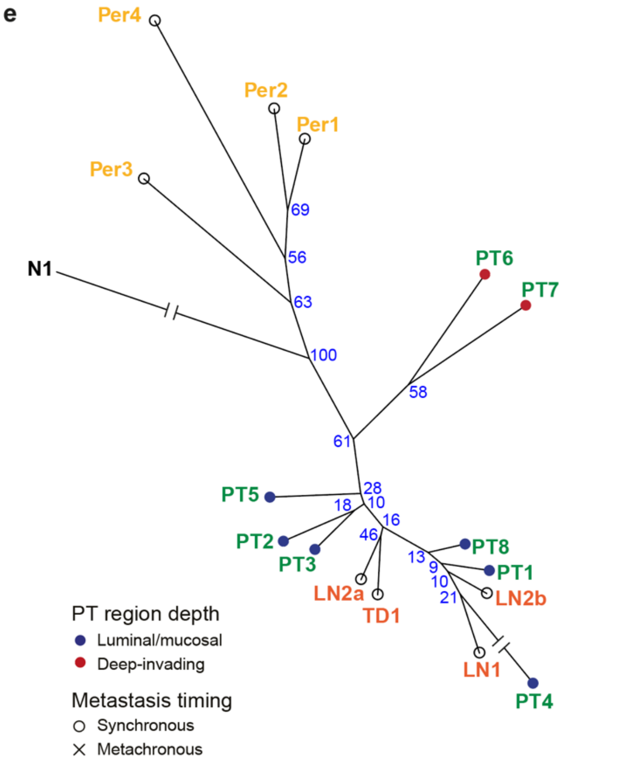


f

| | *k* | *l* | *m* | RDS |
|---|---|---|---|---|
| Peritoneum | 8 | 4 | 4 | $2.2 \times 10^{-3}$ |
| Lymph node | 8 | 1 | 2 | 1 |
| Tumor deposit | 8 | 1 | 1 | n/a |
| Liver | 0 | 0 | 0 | n/a |

**Supplementary Figure 3.** Additional information for patient C157.

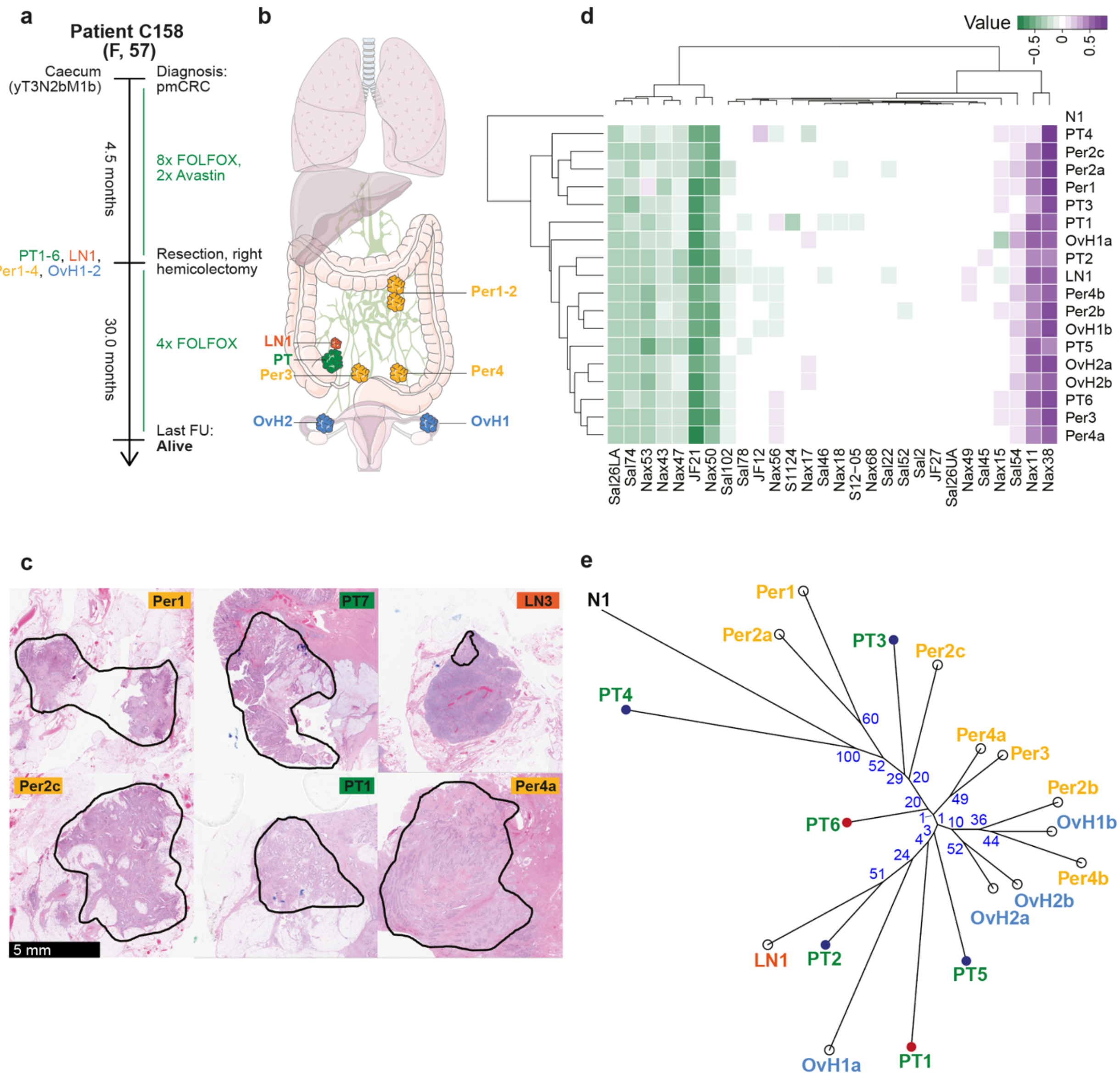


f

| | *k* | *l* | *m* | RDS |
|---|---|---|---|---|
| Peritoneum | 6 | 2 | 4 | 0.341 |
| Lymph node | 6 | 1 | 1 | n/a |
| Tumor deposit | 0 | 0 | 0 | n/a |
| Liver | 0 | 0 | 0 | n/a |

**Supplementary Figure 3.** Additional information for patient C158.

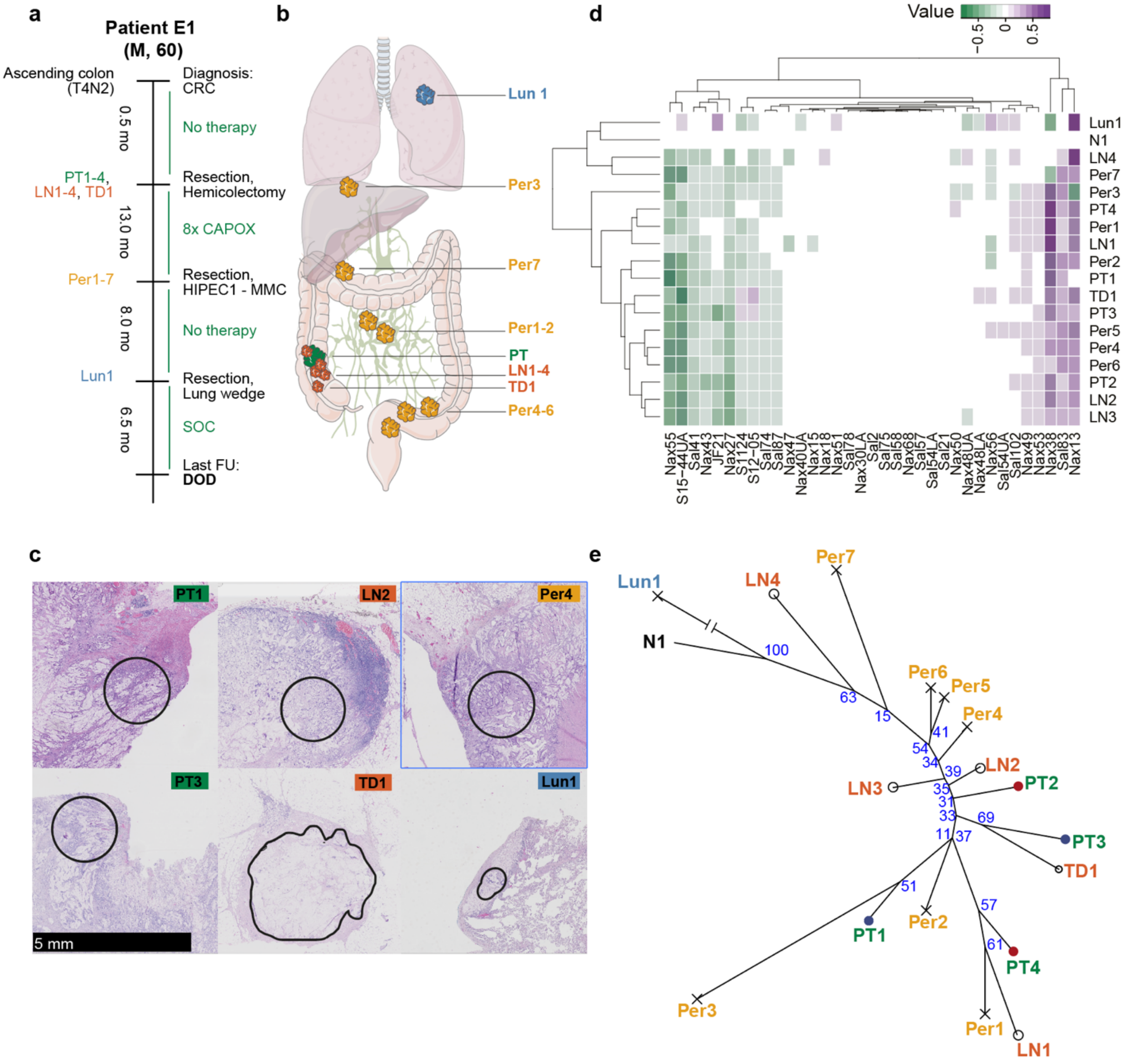


| f | *k* | *l* | *m* | RDS |
|---|---|---|---|---|
| Peritoneum | 4 | 4 | 7 | 0.108 |
| Lymph node | 4 | 2 | 4 | 0.441 |
| Tumor deposit | 4 | 1 | 1 | n/a |
| Liver | 0 | 0 | 0 | n/a |

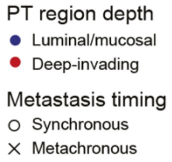


**Supplementary Figure 3.** Additional information for patient E1.

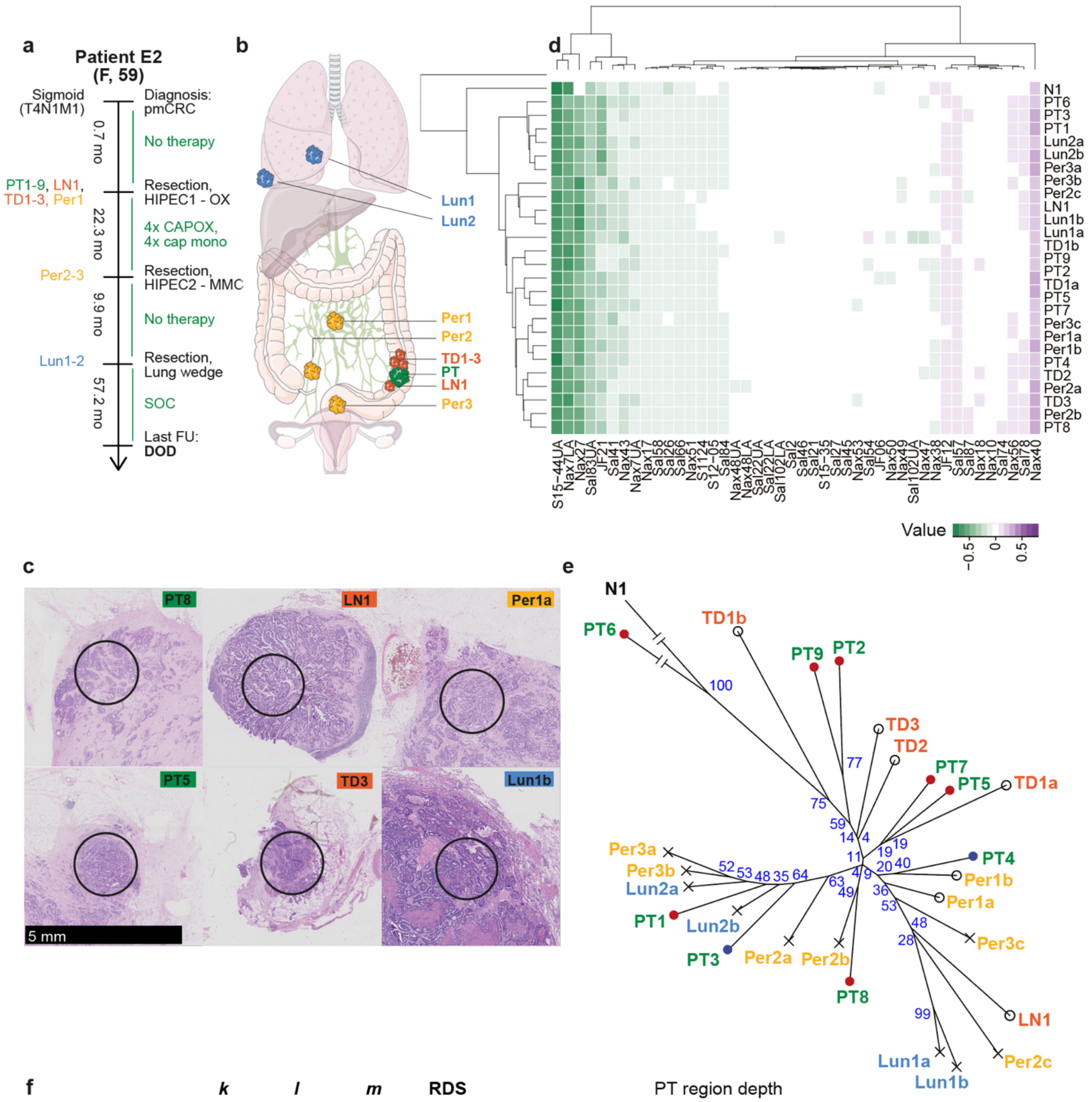


f

| | *k* | *l* | *m* | RDS |
|---|---|---|---|---|
| Peritoneum | 9 | 1 | 3 | 1 |
| Lymph node | 9 | 1 | 1 | n/a |
| Tumor deposit | 9 | 1 | 3 | 1 |
| Liver | 0 | 0 | 0 | n/a |

**Supplementary Figure 3.** Additional information for patient E2.

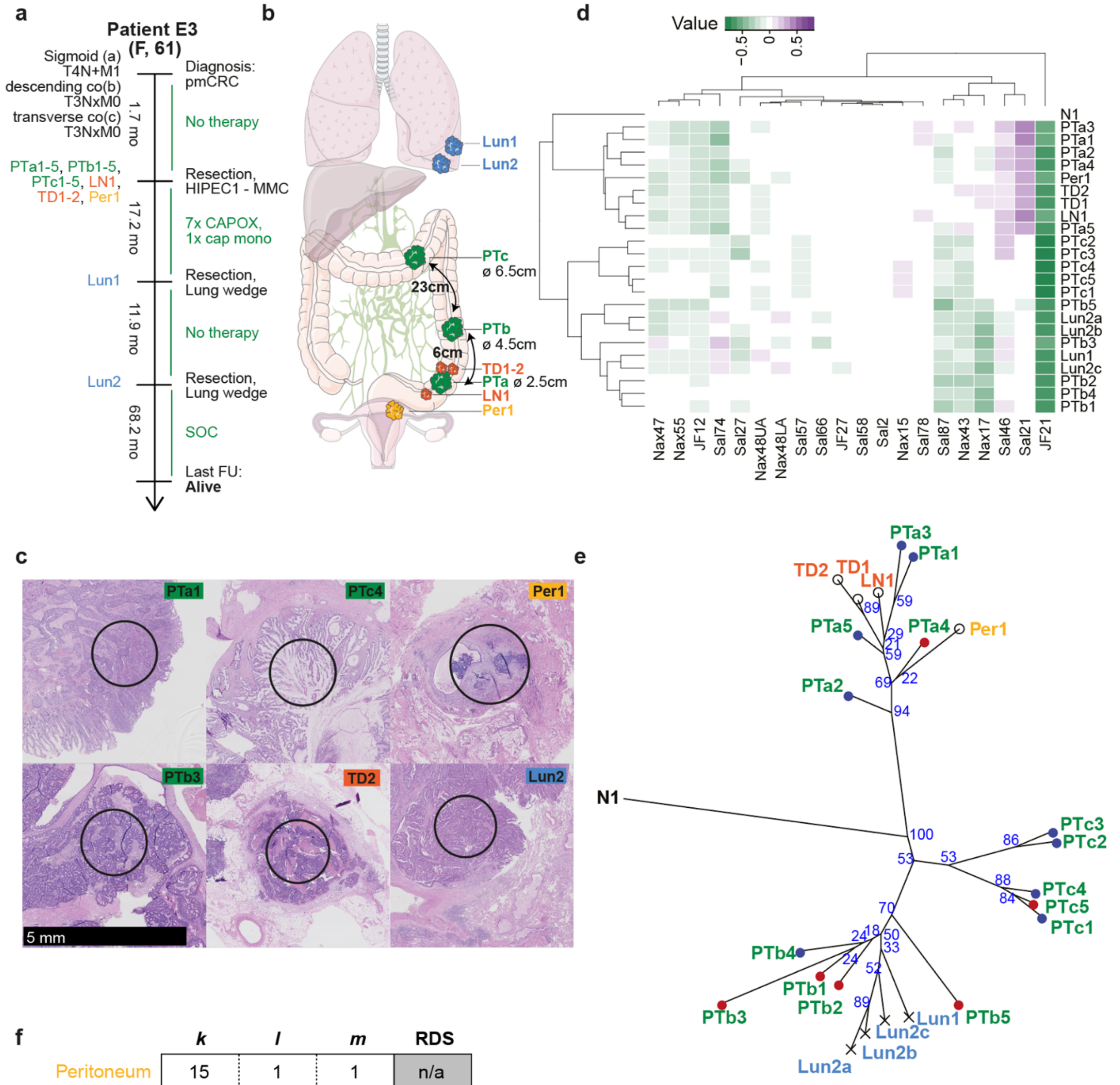


f

| | *k* | *l* | *m* | RDS | |
|---|---|---|---|---|---|
| Peritoneum | 15 | 1 | 1 | n/a | All samples |
| Lymph node | 15 | 1 | 1 | n/a | |
| Tumor deposit | 15 | 2 | 2 | 0.032 | |
| Liver | 0 | 0 | 0 | n/a | |
| Peritoneum | 5 | 1 | 1 | n/a | PTa clade only |
| Lymph node | 5 | 1 | 1 | n/a | |
| Tumor deposit | 5 | 2 | 2 | 0.091 | |
| Liver | 0 | 0 | 0 | n/a | |

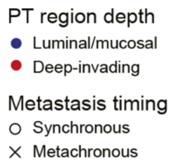


**Supplementary Figure 3.** Additional information for patient E3.

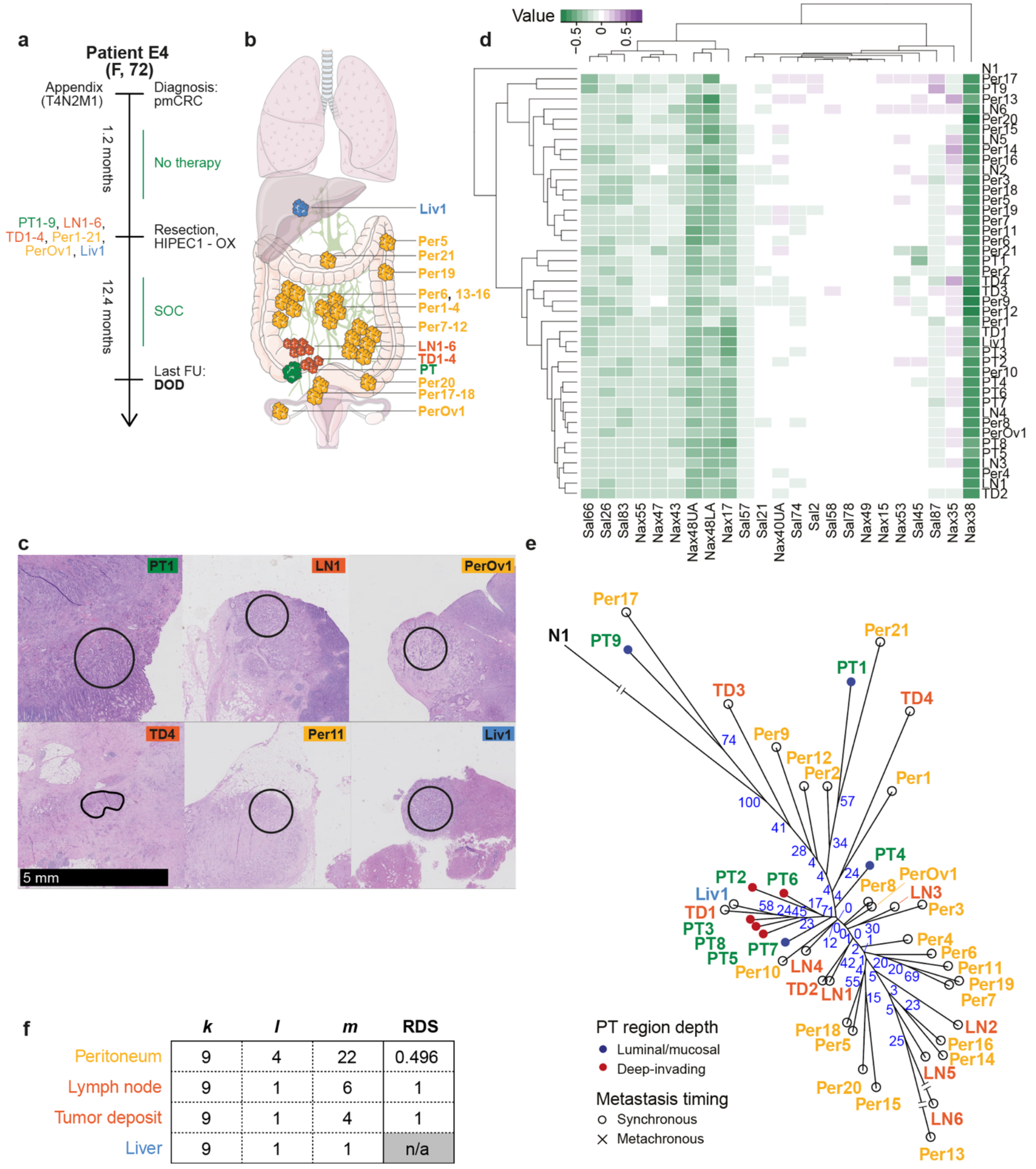


| | *k* | *l* | *m* | RDS |
|---|---|---|---|---|
| Peritoneum | 9 | 4 | 22 | 0.496 |
| Lymph node | 9 | 1 | 6 | 1 |
| Tumor deposit | 9 | 1 | 4 | 1 |
| Liver | 9 | 1 | 1 | n/a |

**Supplementary Figure 3.** Additional information for patient E4.

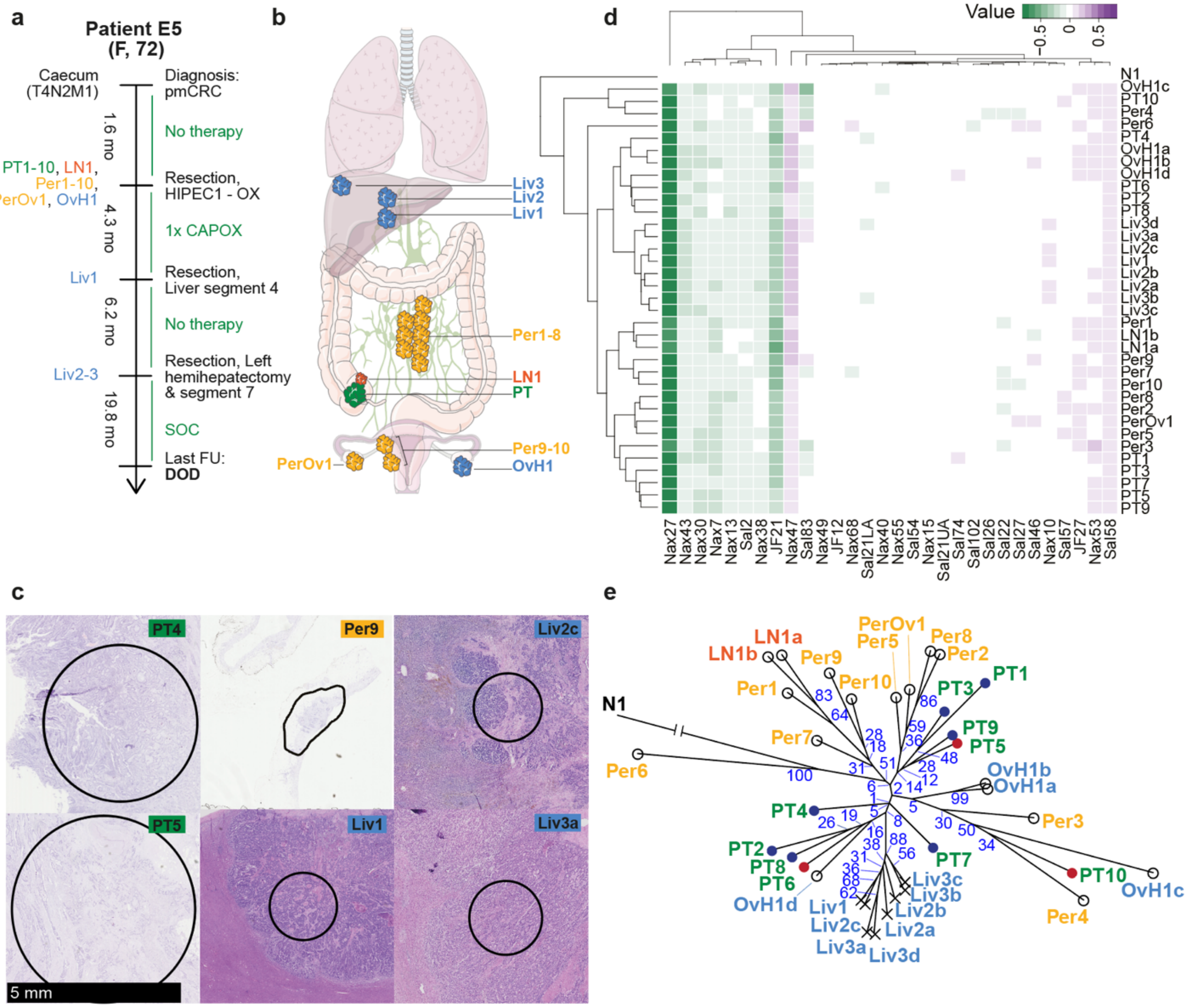


f

| | *k* | *l* | *m* | RDS |
|---|---|---|---|---|
| Peritoneum | 10 | 4 | 11 | 0.097 |
| Lymph node | 10 | 1 | 1 | n/a |
| Tumor deposit | 0 | 0 | 0 | n/a |
| Liver | 10 | 3 | 3 | $6.2x10^{-3}$ |

PT region depth
- ● Luminal/mucosal
- ● Deep-invading

Metastasis timing
- ○ Synchronous
- × Metachronous

**Supplementary Figure 3.** Additional information for patient E5.

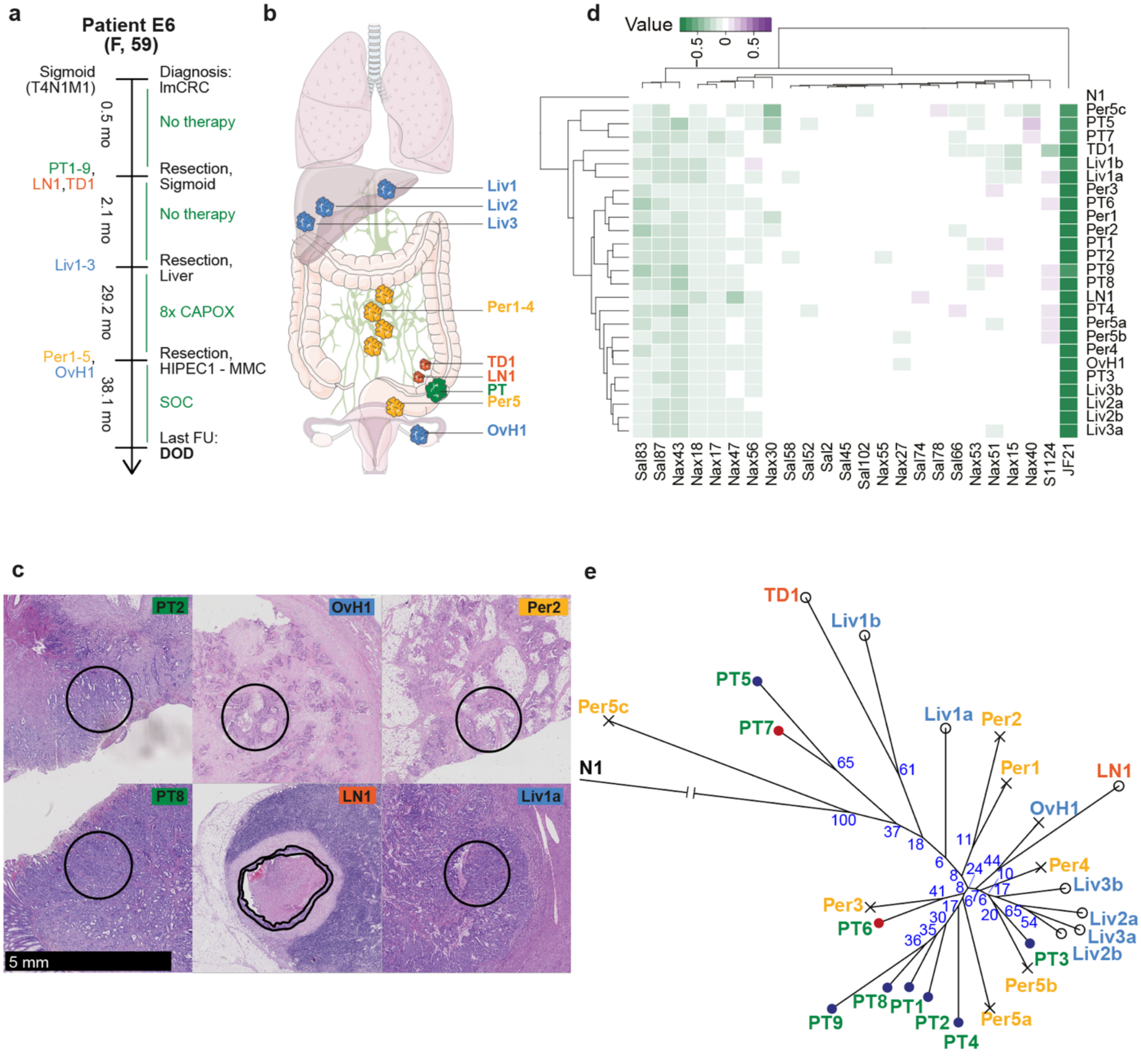


f

| | *k* | *l* | *m* | RDS |
|---|---|---|---|---|
| Peritoneum | 9 | 1 | 5 | 1 |
| Lymph node | 9 | 1 | 1 | n/a |
| Tumor deposit | 9 | 1 | 1 | n/a |
| Liver | 9 | 2 | 3 | 0.143 |

**Supplementary Figure 3.** Additional information for patient E6.

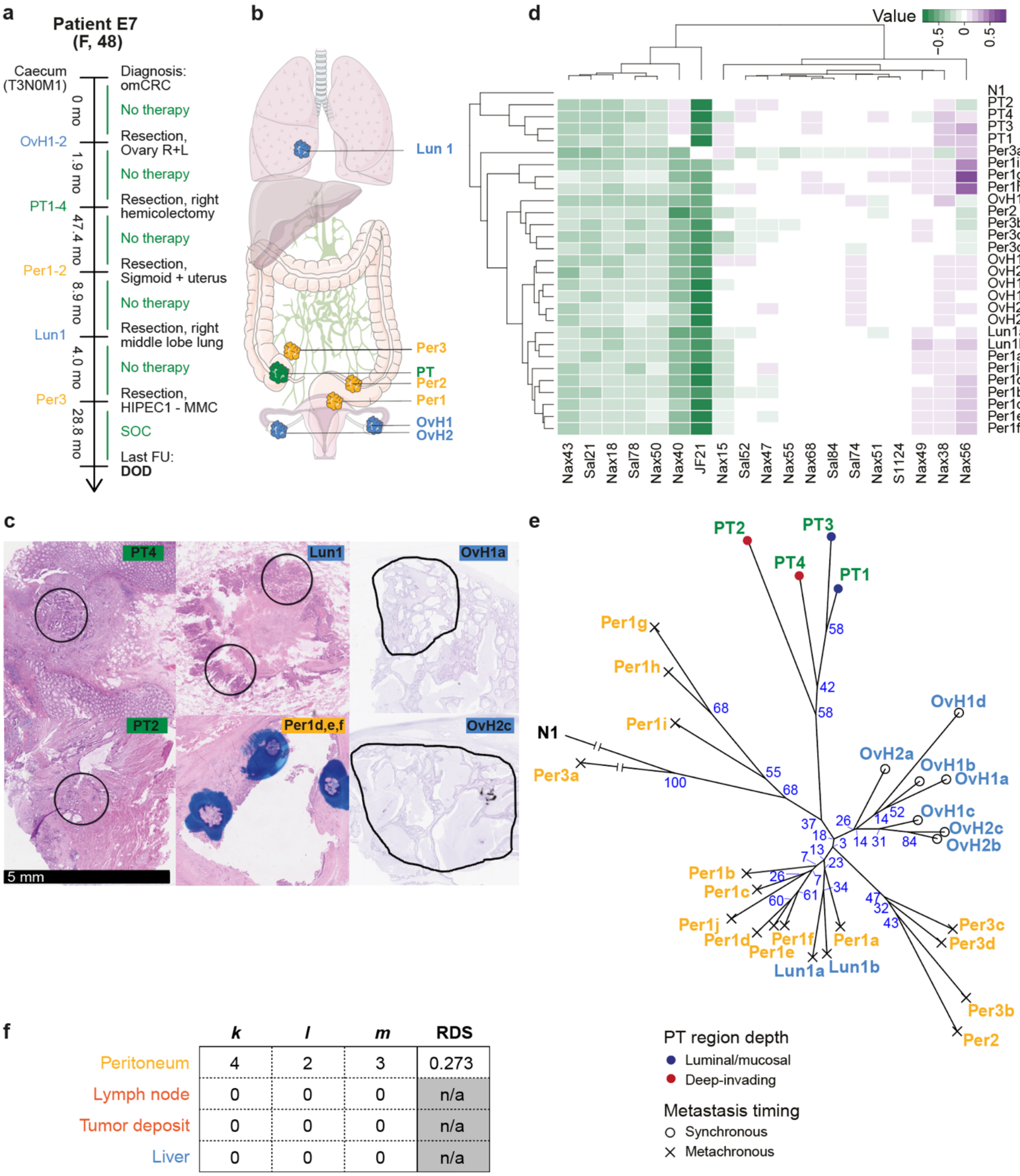


| | *k* | *l* | *m* | RDS |
|---|---|---|---|---|
| Peritoneum | 4 | 2 | 3 | 0.273 |
| Lymph node | 0 | 0 | 0 | n/a |
| Tumor deposit | 0 | 0 | 0 | n/a |
| Liver | 0 | 0 | 0 | n/a |

**Supplementary Figure 3.** Additional information for patient E7.

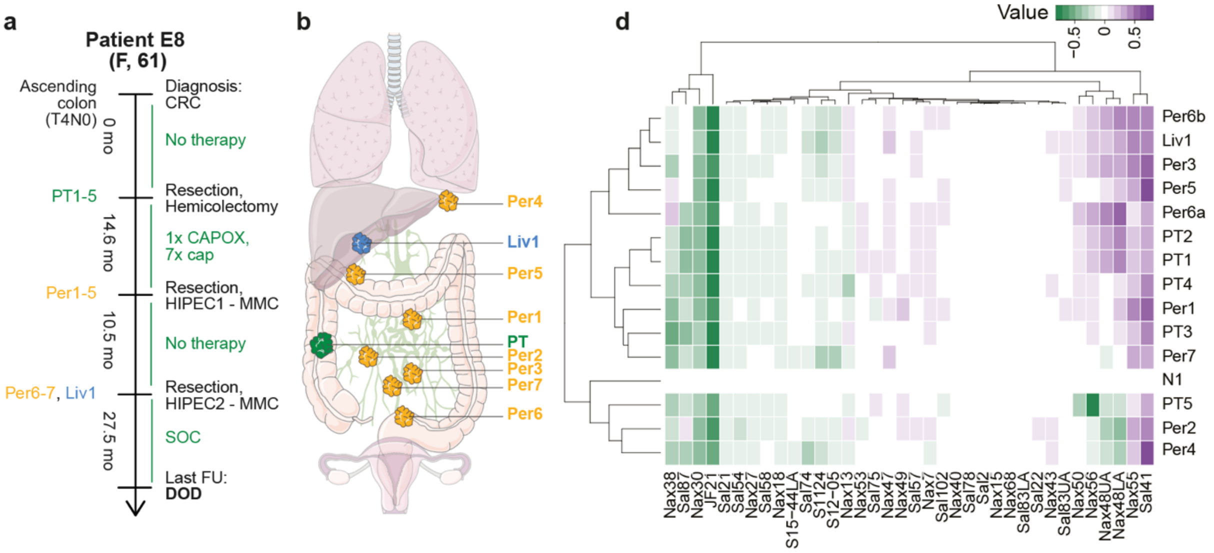


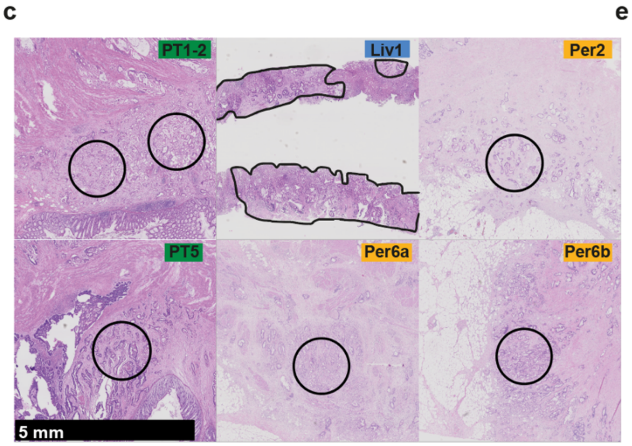


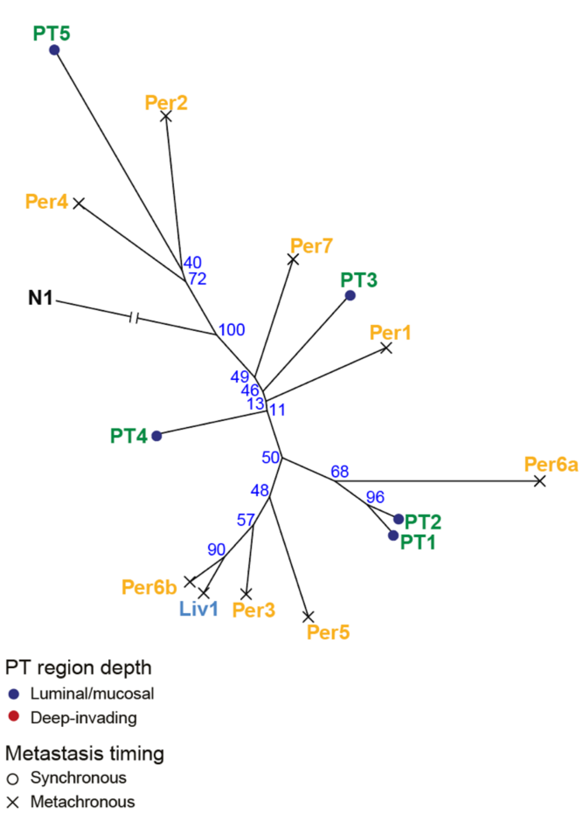


f

| | *k* | *l* | *m* | RDS |
|---|---|---|---|---|
| Peritoneum | 5 | 2 | 7 | 0.752 |
| Lymph node | 0 | 0 | 0 | n/a |
| Tumor deposit | 0 | 0 | 0 | n/a |
| Liver | 5 | 1 | 1 | n/a |

**Supplementary Figure 3.** Additional information for patient E8.

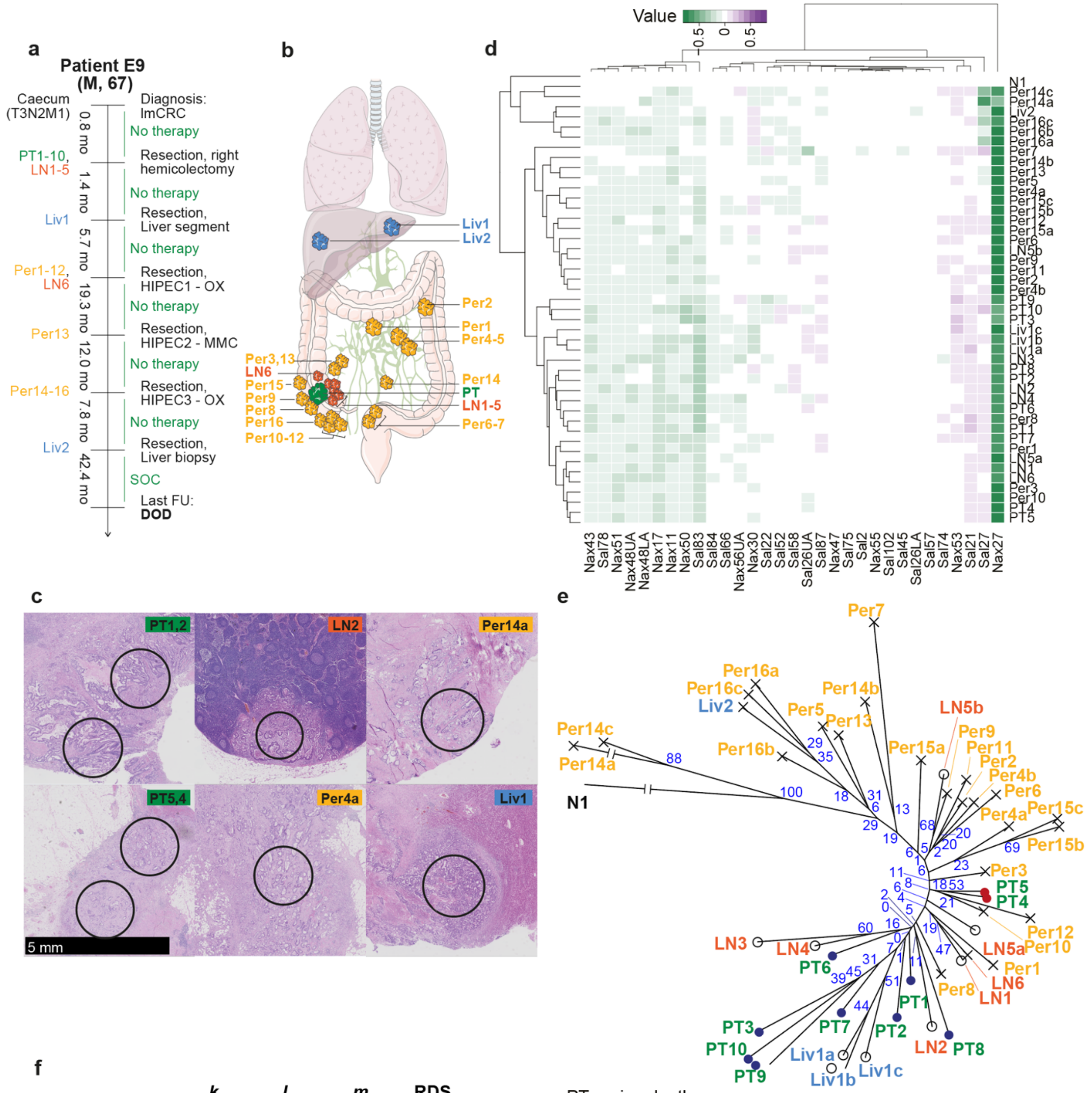


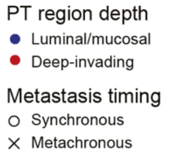


**f**

| | *k* | *l* | *m* | RDS |
|---|---|---|---|---|
| Peritoneum | 10 | 4 | 16 | 0.249 |
| Lymph node | 10 | 2 | 6 | 0.461 |
| Tumor deposit | 0 | 0 | 0 | n/a |
| Liver | 10 | 1 | 2 | 1 |

**Supplementary Figure 3.** Additional information for patient E9.

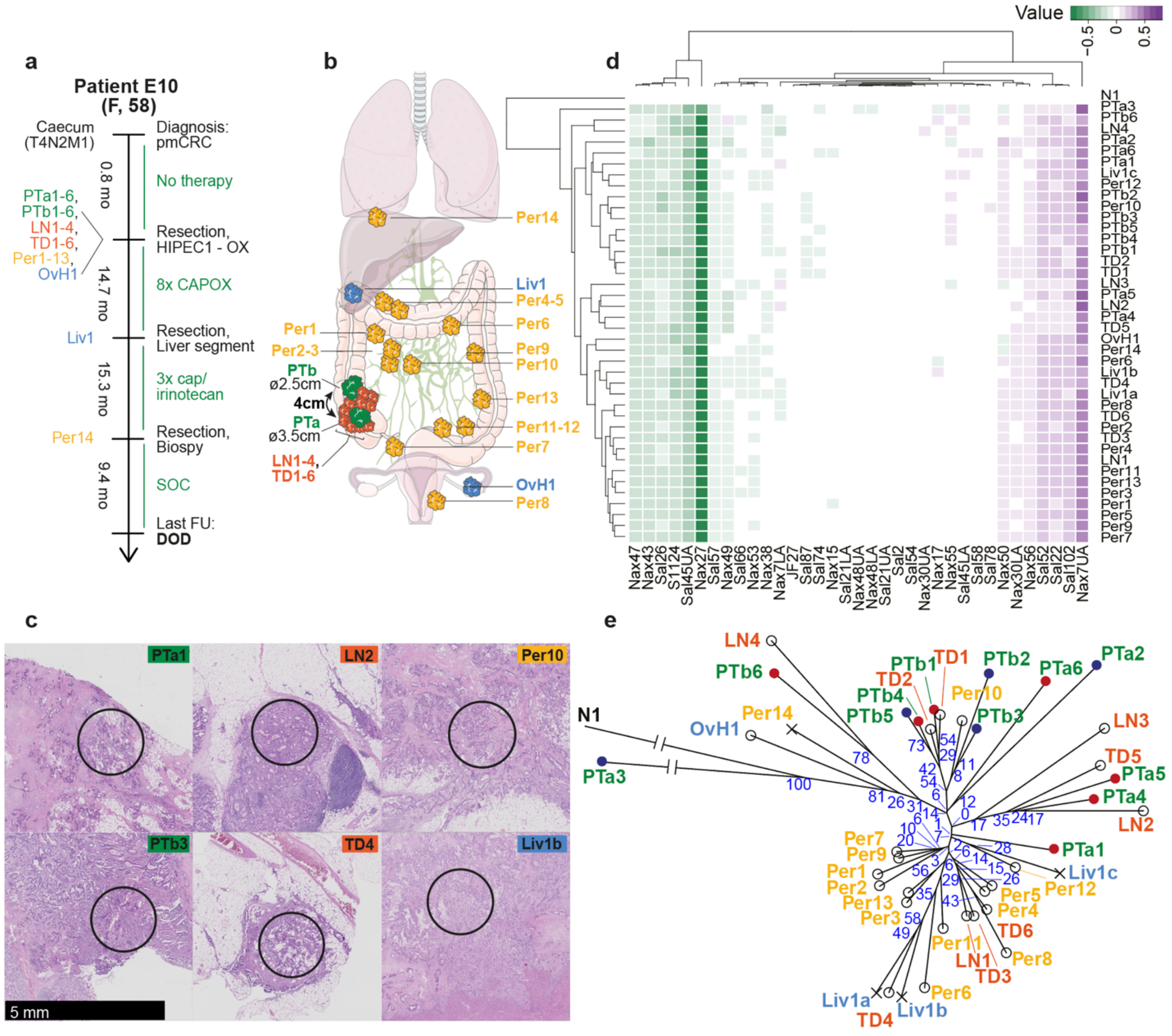


f

| | *k* | *l* | *m* | RDS |
|---|---|---|---|---|
| Peritoneum | 12 | 4 | 14 | 0.142 |
| Lymph node | 12 | 1 | 4 | 1 |
| Tumor deposit | 12 | 2 | 6 | 0.411 |
| Liver | 12 | 1 | 1 | n/a |

PT region depth
- ● Luminal/mucosal
- ● Deep-invading

Metastasis timing
- ○ Synchronous
- × Metachronous

**Supplementary Figure 3.** Additional information for patient E10.

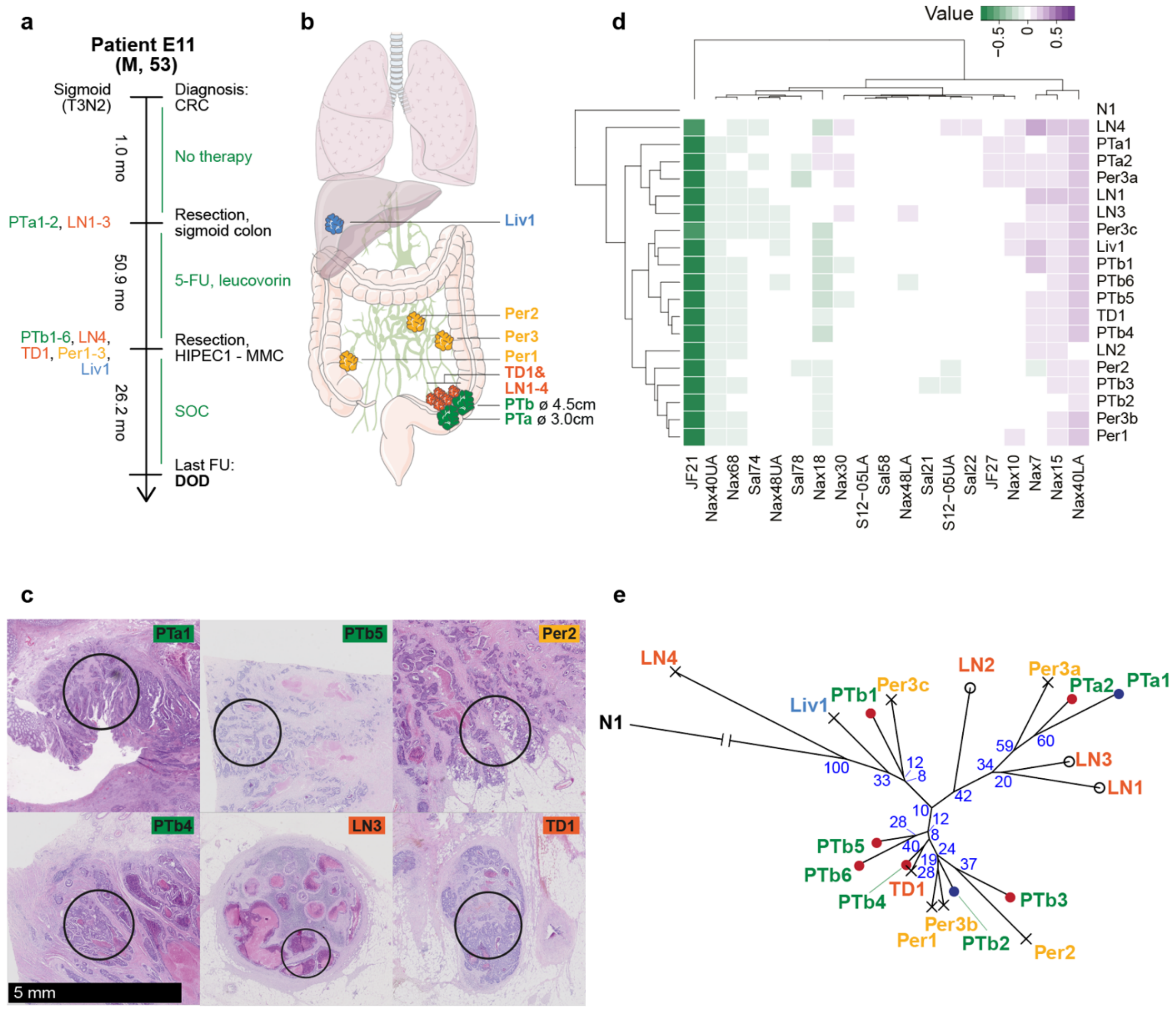


f

| | *k* | *l* | *m* | RDS |
|---|---|---|---|---|
| Peritoneum | 8 | 1 | 3 | 1 |
| Lymph node | 8 | 2 | 4 | 0.278 |
| Tumor deposit | 8 | 1 | 1 | n/a |
| Liver | 8 | 1 | 1 | n/a |

PT region depth
- ● Luminal/mucosal
- ● Deep-invading

Metastasis timing
- ○ Synchronous
- × Metachronous

**Supplementary Figure 3.** Additional information for patient E11.

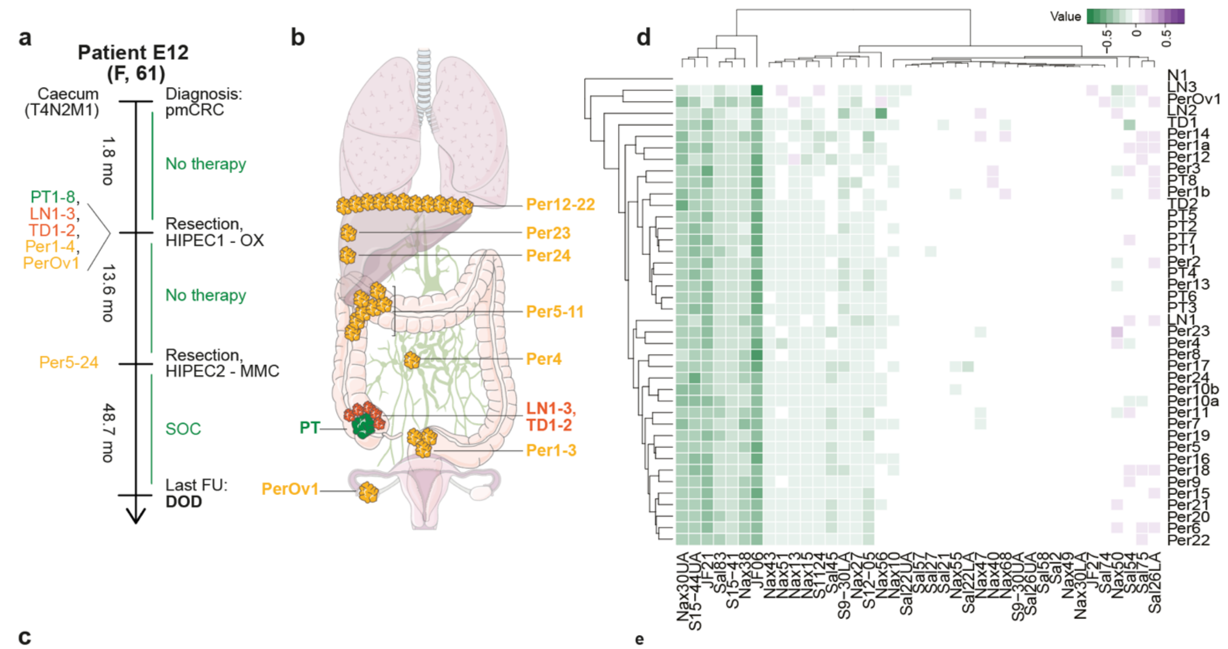


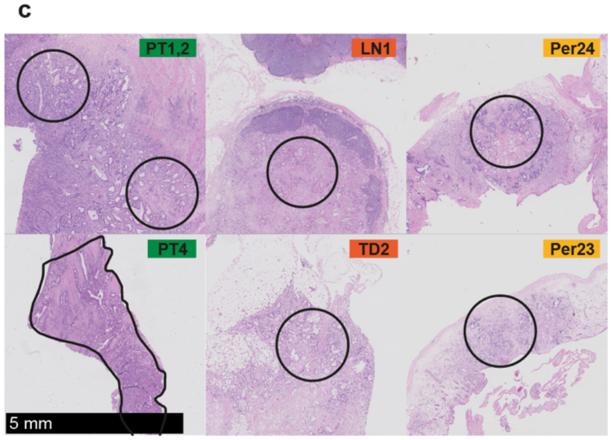


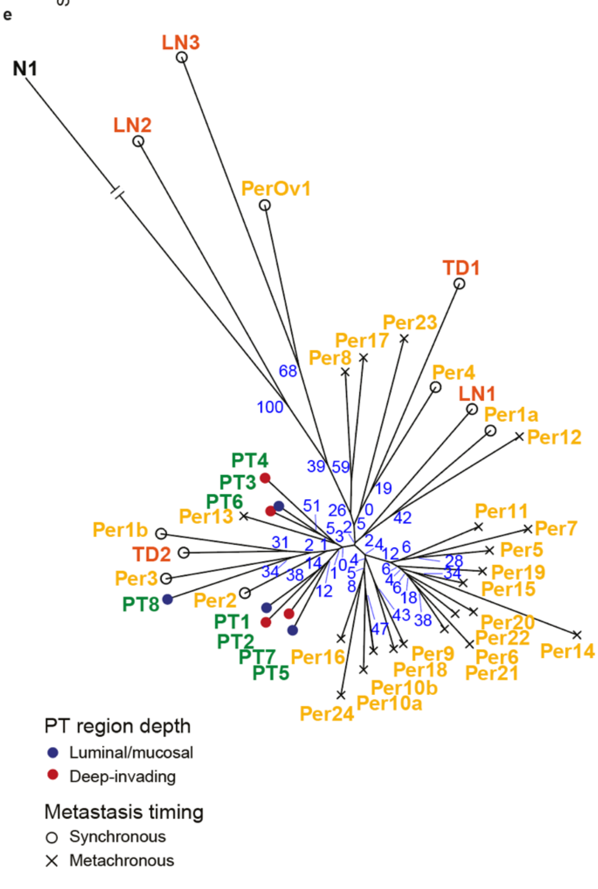


f

| | *k* | *l* | *m* | RDS |
|---|---|---|---|---|
| Peritoneum | 8 | 16 | 25 | 7.1x10$^{-4}$ |
| Lymph node | 8 | 1 | 3 | 1 |
| Tumor deposit | 8 | 1 | 2 | 1 |
| Liver | 0 | 0 | 0 | n/a |

**Supplementary Figure 3.** Additional information for patient E12.

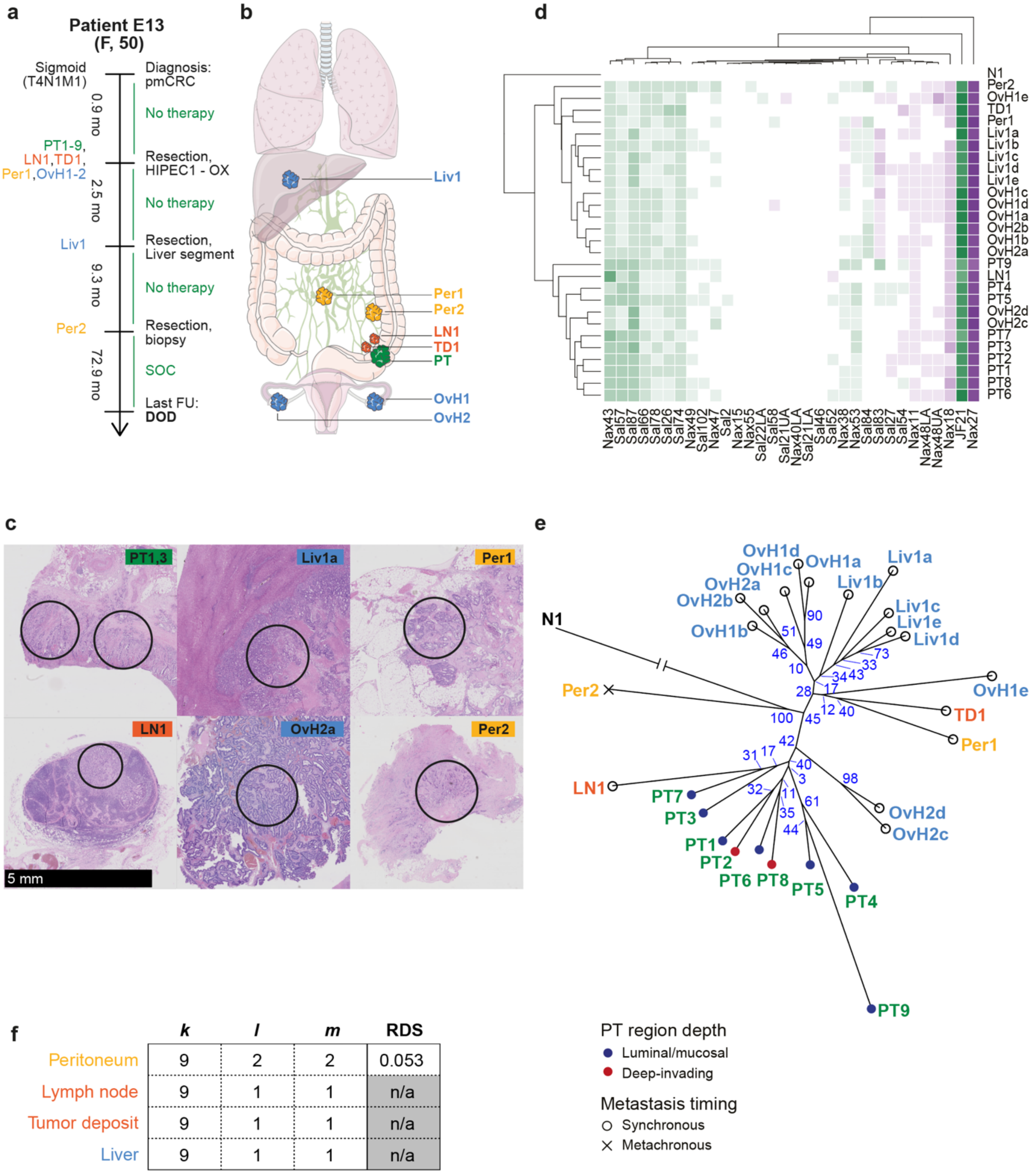


**f**

| | *k* | *l* | *m* | RDS |
|---|---|---|---|---|
| Peritoneum | 9 | 2 | 2 | 0.053 |
| Lymph node | 9 | 1 | 1 | n/a |
| Tumor deposit | 9 | 1 | 1 | n/a |
| Liver | 9 | 1 | 1 | n/a |

**Supplementary Figure 3.** Additional information for patient E13.

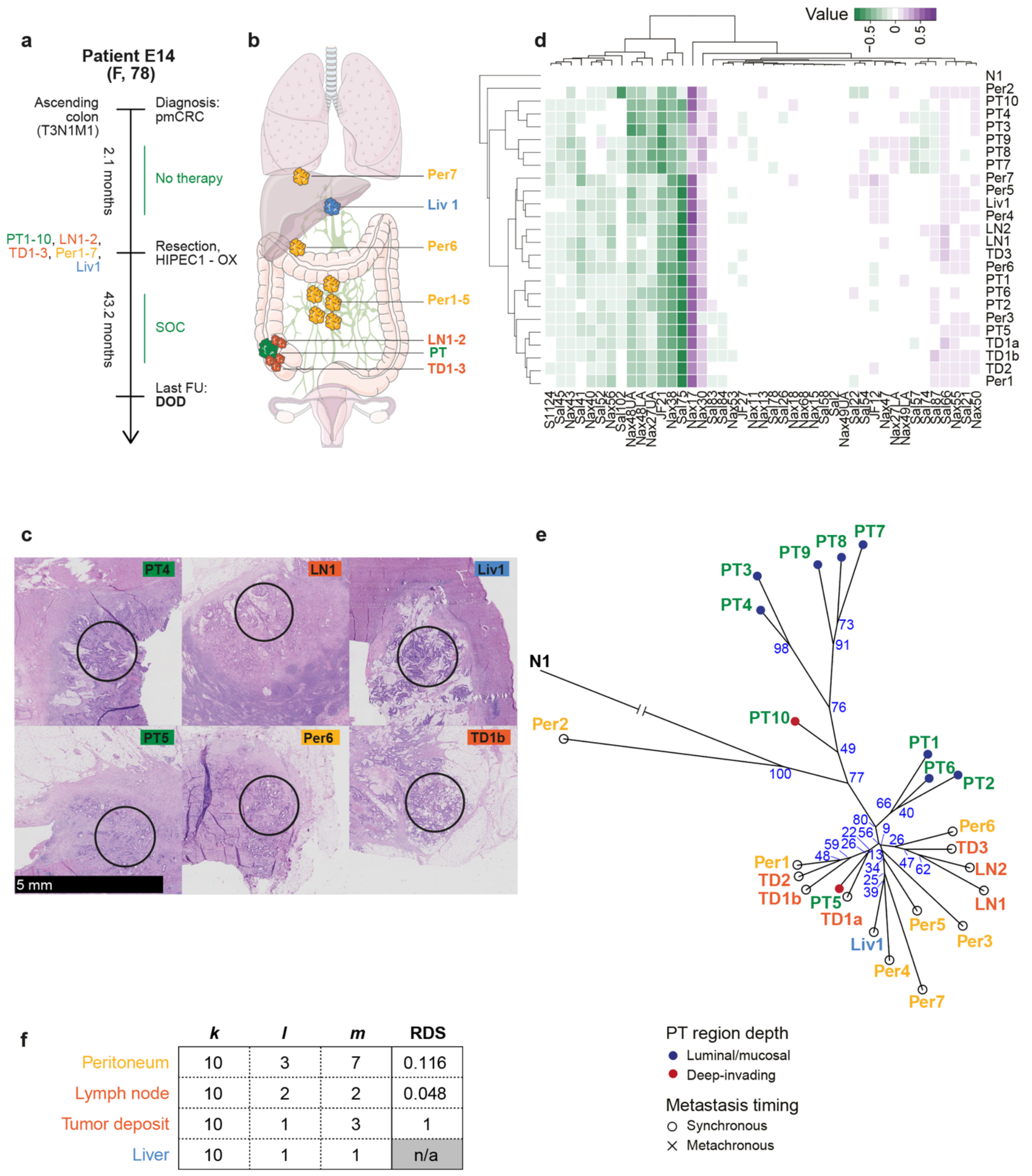


| | *k* | *l* | *m* | RDS |
|---|---|---|---|---|
| Peritoneum | 10 | 3 | 7 | 0.116 |
| Lymph node | 10 | 2 | 2 | 0.048 |
| Tumor deposit | 10 | 1 | 3 | 1 |
| Liver | 10 | 1 | 1 | n/a |

**Supplementary Figure 3.** Additional information for patient E14.

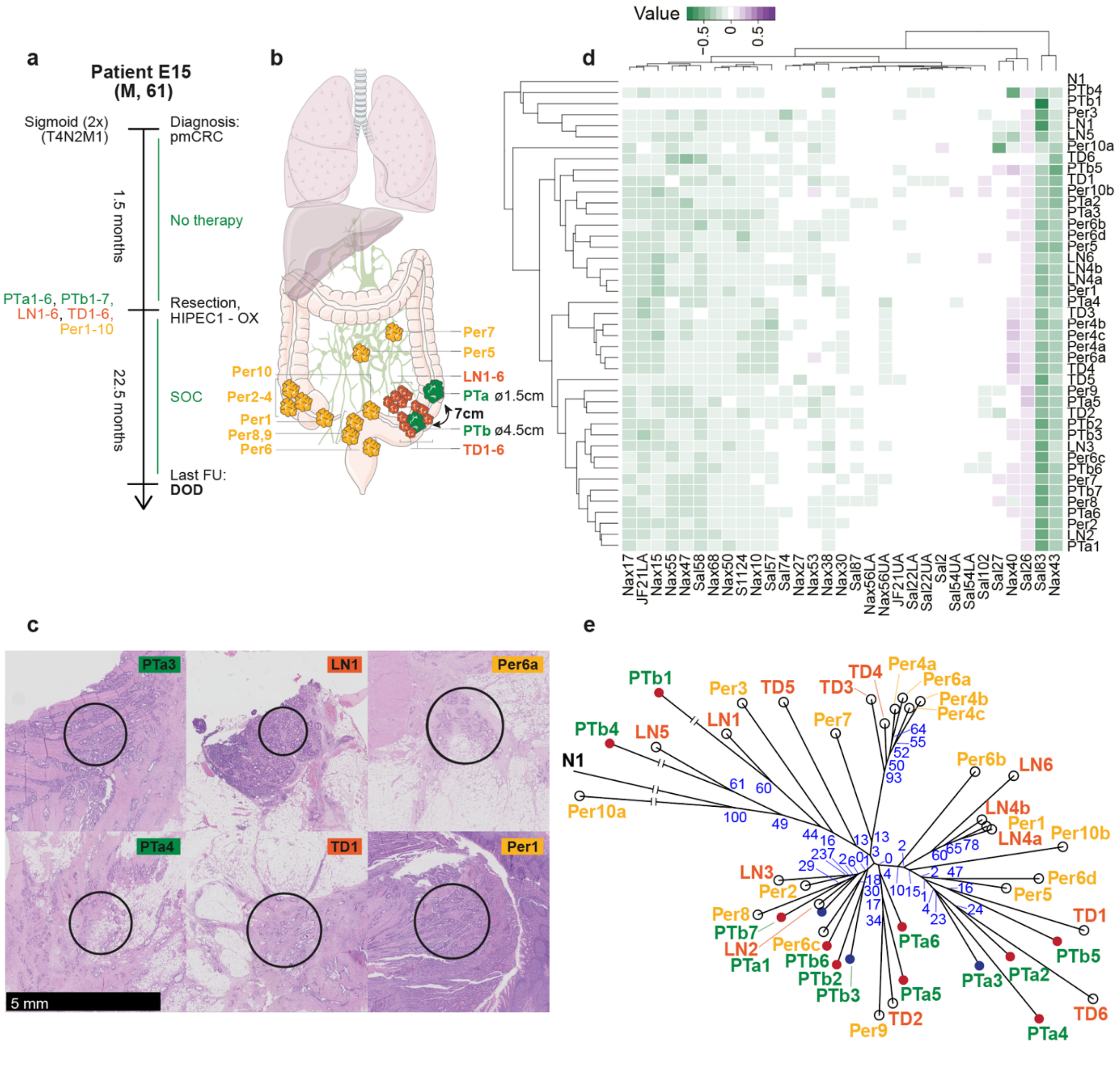


f

| | *k* | *l* | *m* | RDS |
|---|---|---|---|---|
| Peritoneum | 13 | 2 | 10 | 0.733 |
| Lymph node | 13 | 2 | 6 | 0.39 |
| Tumor deposit | 13 | 2 | 6 | 0.39 |
| Liver | 0 | 0 | 0 | n/a |

PT region depth
- ● Luminal/mucosal
- ● Deep-invading

Metastasis timing
- ○ Synchronous
- × Metachronous

**Supplementary Figure 3.** Additional information for patient E15.

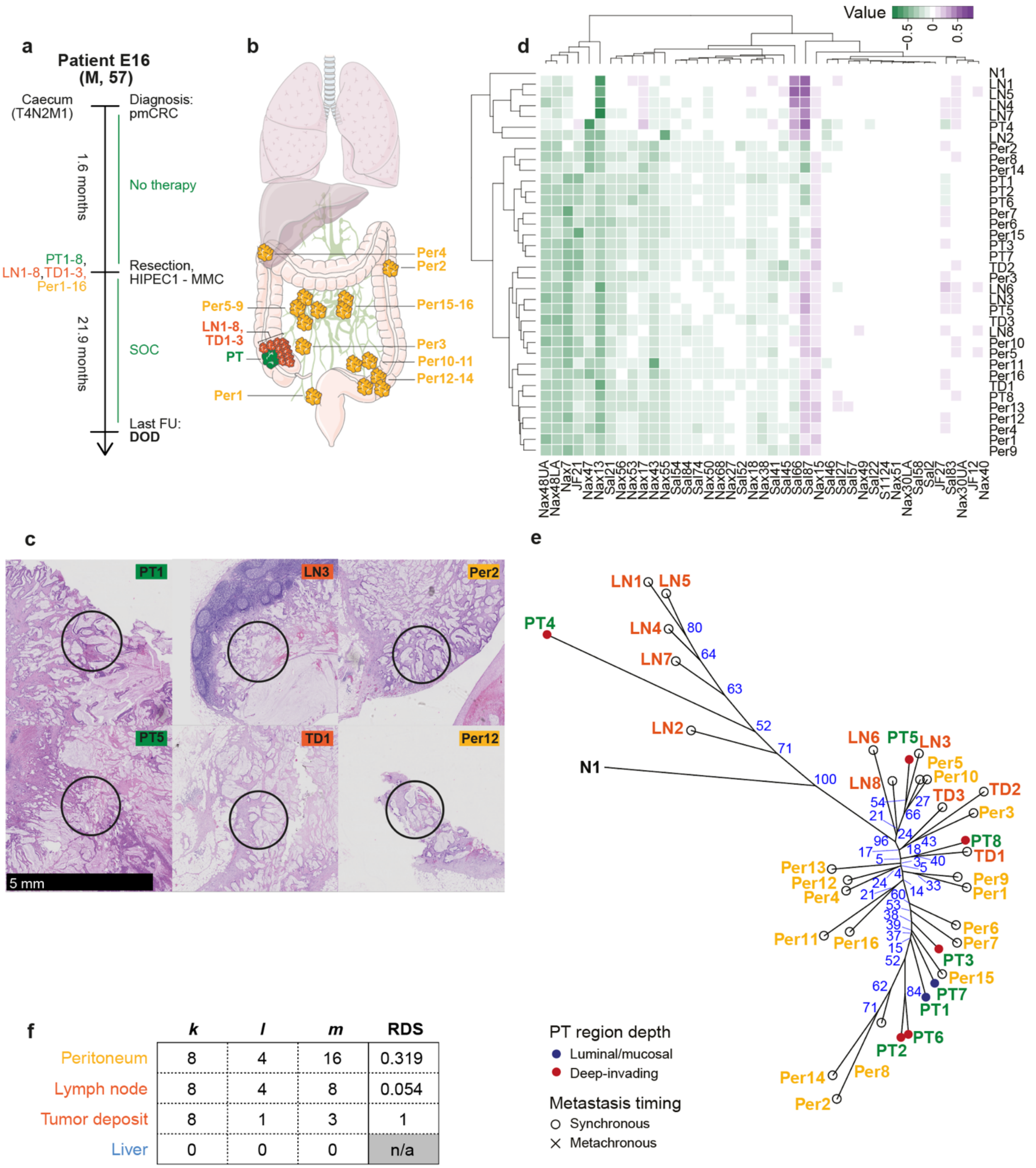


| | *k* | *l* | *m* | RDS |
|---|---|---|---|---|
| Peritoneum | 8 | 4 | 16 | 0.319 |
| Lymph node | 8 | 4 | 8 | 0.054 |
| Tumor deposit | 8 | 1 | 3 | 1 |
| Liver | 0 | 0 | 0 | n/a |

**Supplementary Figure 3.** Additional information for patient E16.

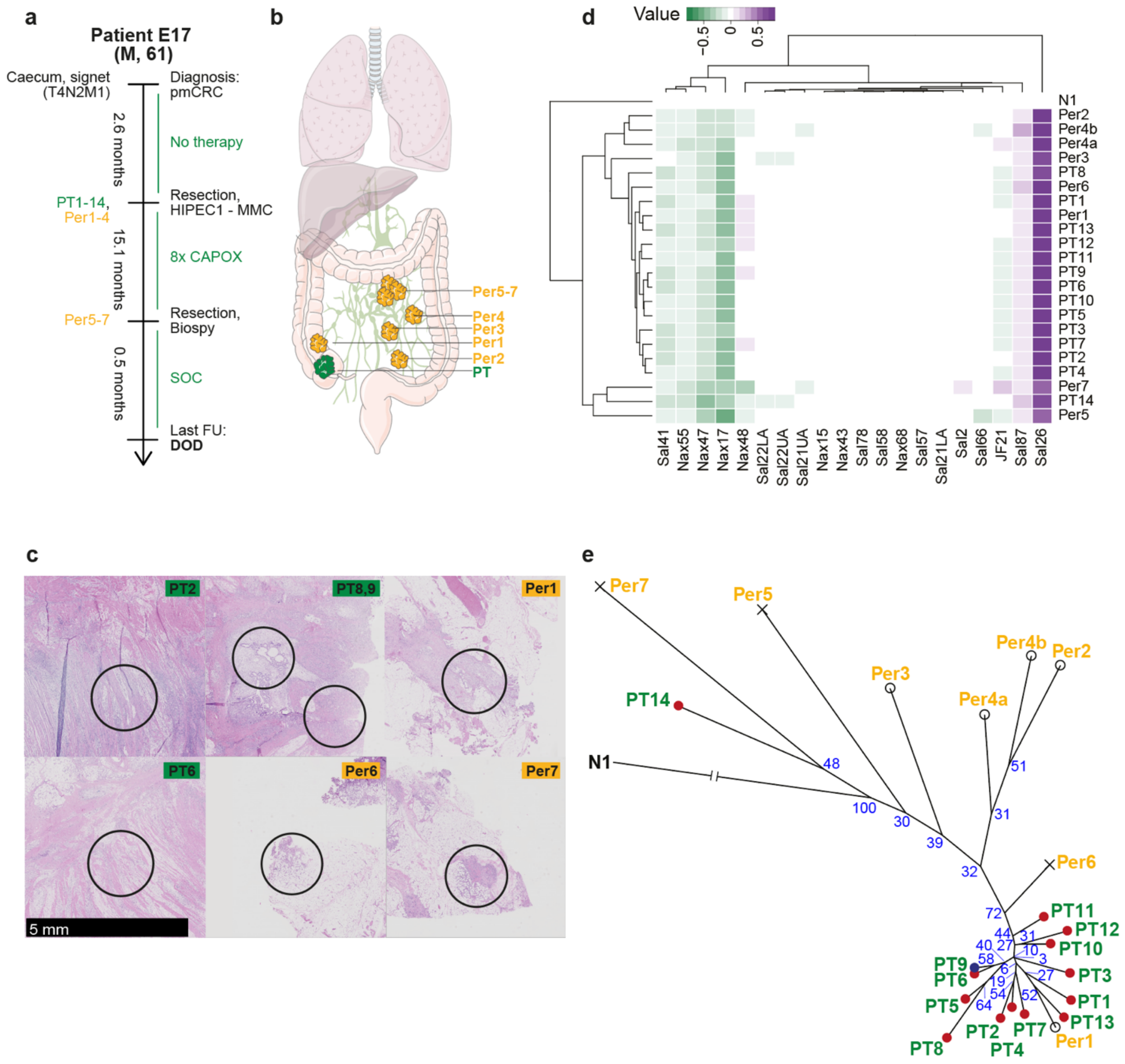


**f**

| | *k* | *l* | *m* | RDS |
|---|---|---|---|---|
| Peritoneum | 14 | 2 | 7 | 0.468 |
| Lymph node | 0 | 0 | 0 | n/a |
| Tumor deposit | 0 | 0 | 0 | n/a |
| Liver | 0 | 0 | 0 | n/a |

PT region depth
- Luminal/mucosal
- Deep-invading

Metastasis timing
- ○ Synchronous
- × Metachronous

**Supplementary Figure 3.** Additional information for patient E17.

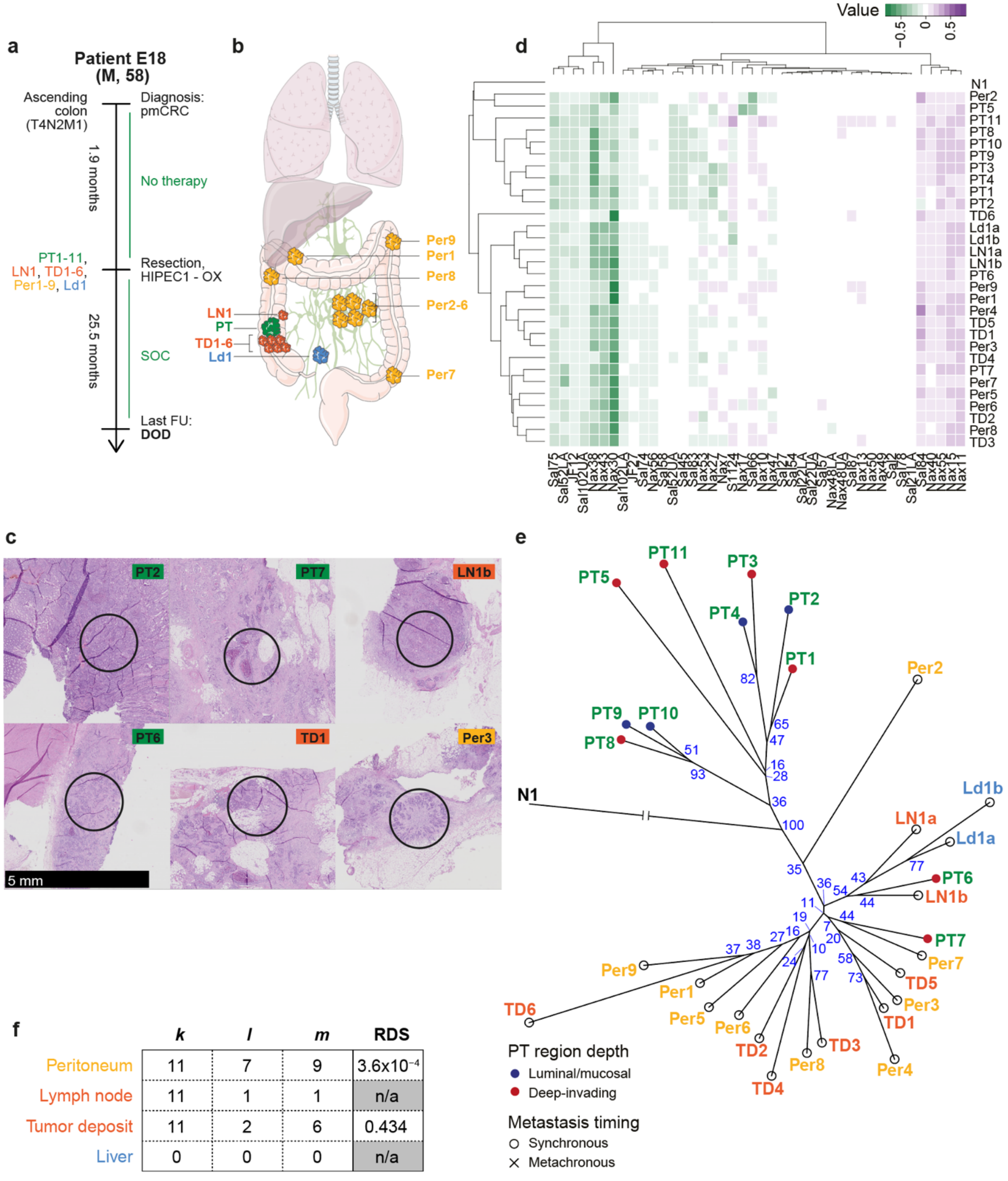


f

| | *k* | *l* | *m* | RDS |
|---|---|---|---|---|
| Peritoneum | 11 | 7 | 9 | $3.6x10^{-4}$ |
| Lymph node | 11 | 1 | 1 | n/a |
| Tumor deposit | 11 | 2 | 6 | 0.434 |
| Liver | 0 | 0 | 0 | n/a |

**Supplementary Figure 3.** Additional information for patient E18.

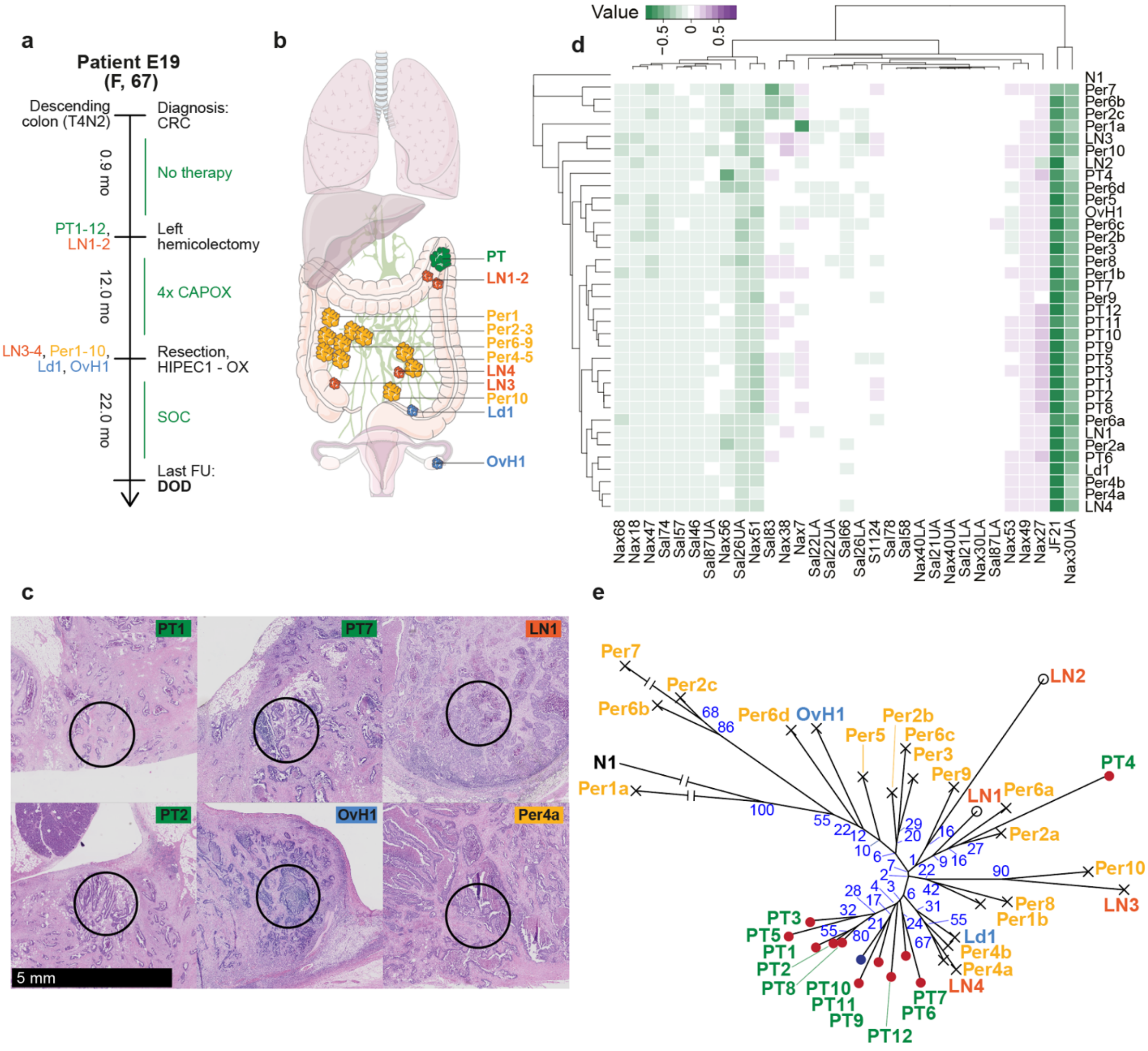


f

| | *k* | *l* | *m* | RDS |
|---|---|---|---|---|
| Peritoneum | 12 | 2 | 10 | 0.755 |
| Lymph node | 12 | 1 | 4 | 1 |
| Tumor deposit | 0 | 0 | 0 | n/a |
| Liver | 0 | 0 | 0 | n/a |

PT region depth
- ● Luminal/mucosal
- ● Deep-invading

Metastasis timing
- ○ Synchronous
- × Metachronous

**Supplementary Figure 3.** Additional information for patient E19.

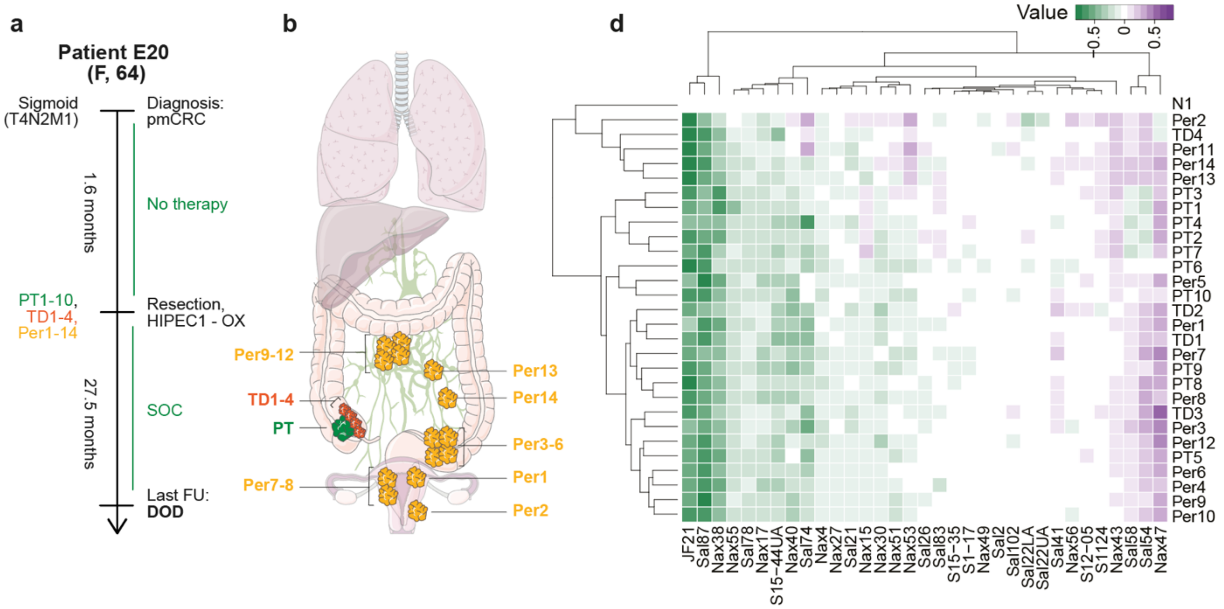


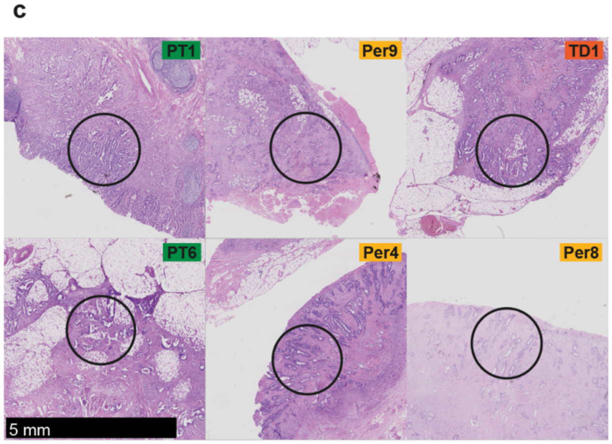


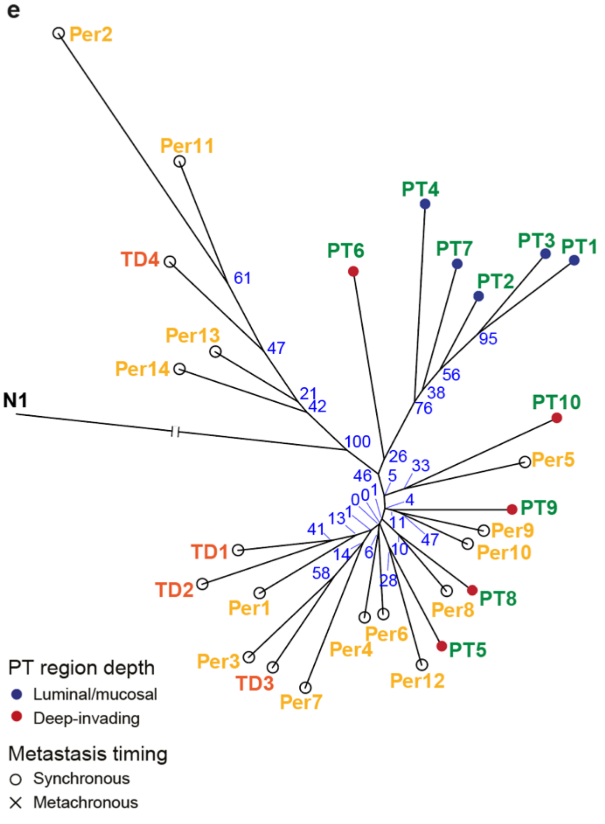


f

| | *k* | *l* | *m* | RDS |
|---|---|---|---|---|
| Peritoneum | 10 | 3 | 14 | 0.481 |
| Lymph node | 0 | 0 | 0 | n/a |
| Tumor deposit | 10 | 2 | 4 | 0.235 |
| Liver | 0 | 0 | 0 | n/a |

**Supplementary Figure 3.** Additional information for patient E20.

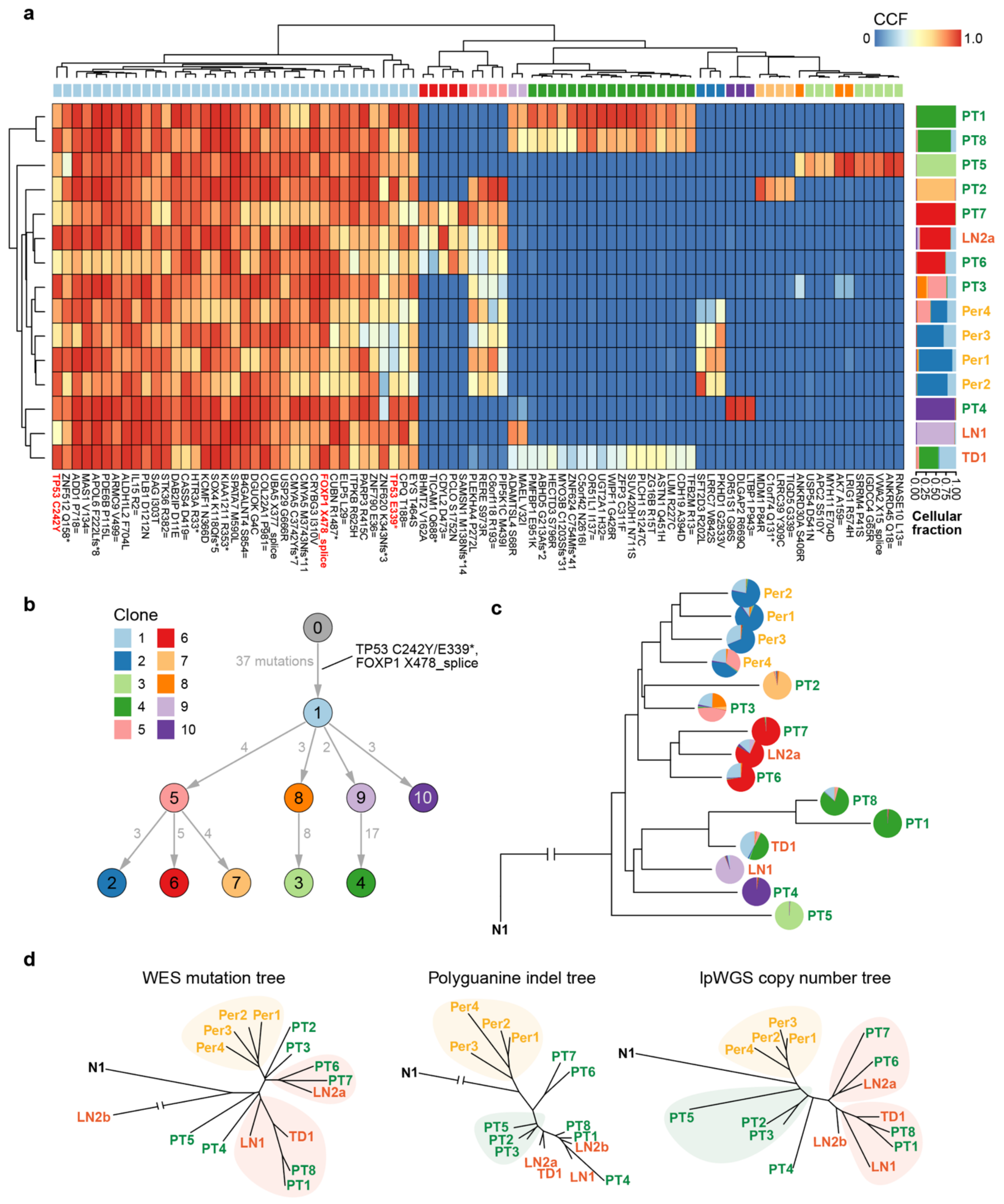


**Supplementary Figure 4. Clonal evolution in patient C157. a,** Heatmap of mutation cancer cell fractions (CCFs) from whole-exome sequencing (WES) data for patient C157. Red text, variants predicted to have oncogenic effects by OncoKB. Top track, clone to which each mutation is assigned (10 clones inferred across all tumor samples using PyClone-VI). Right track, the estimated proportion of each clone in each bulk tumor sample. Note, sample LN2b was an outlier in the WES data, with 3.2-fold the median number of clonal mutations (2.7-fold the median number of variants overall); we therefore excluded LN2b from this analysis to avoid potential false positives. **b,** Inferred clone phylogeny for C157 based on the mutations in (a), constructed with Orchard s. Clone 0 represents the patient's germline genotype. Grey text indicates the number of private mutations acquired during the evolution of each new clone. Variants predicted to be oncogenic in black. **c,** Sample phylogeny for patient C157 based on CCFs as in (a). The phylogeny was constructed from Euclidean distances between CCF vectors using neighbor joining. Pie charts indicate the proportion of each clone in the bulk sample as in (a). **d,** Comparison of phylogenies constructed based on CCFs from WES (left), polyguanine fingerprints (center), and somatic copy number alterations called using lpWGS (right).

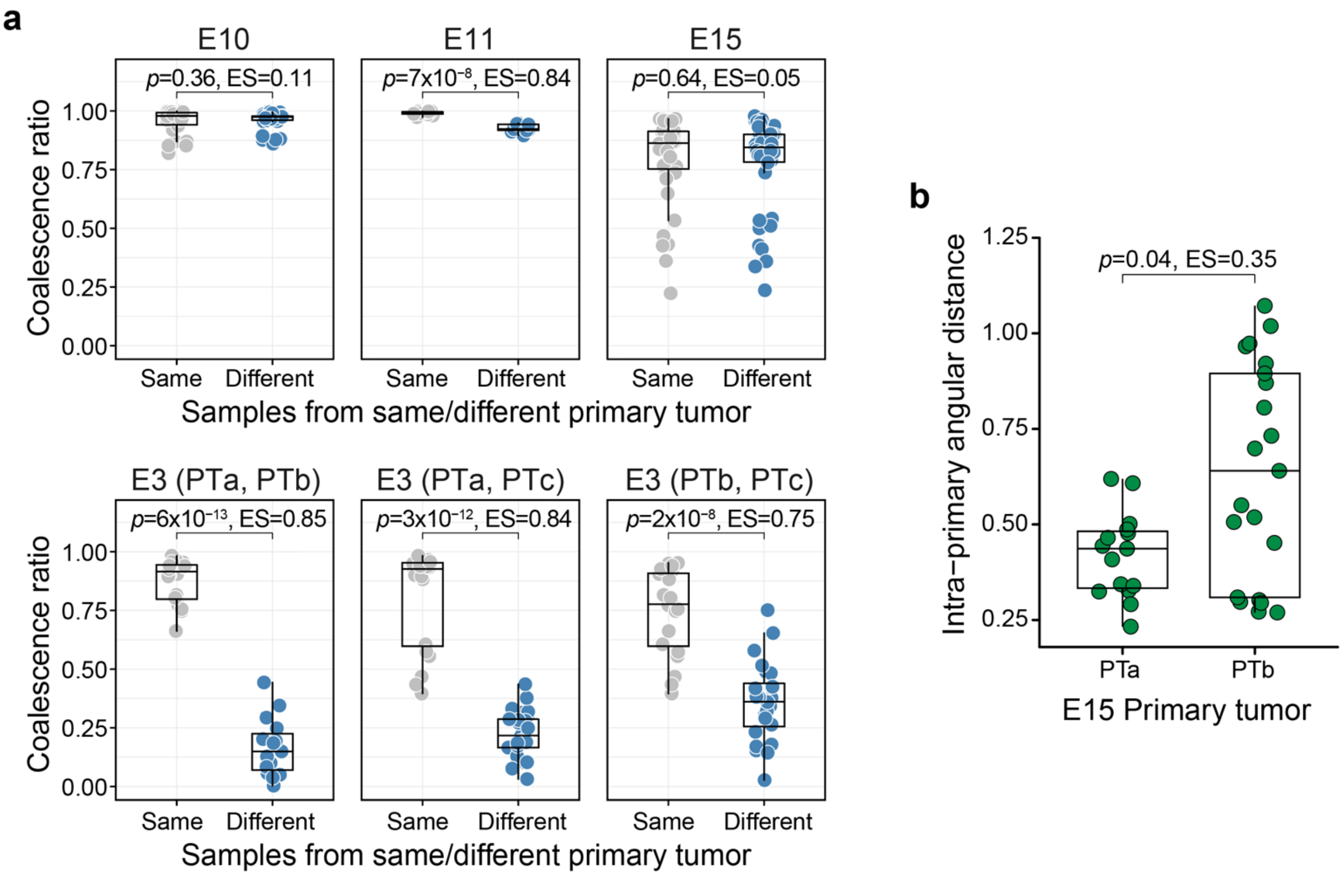


**Supplementary Figure 5. Polyguanine fingerprints distinguish independent vs. clonally related primary tumors. a,** For all patients diagnosed with multiple anatomically distinct primary tumors, coalescence ratios are shown for all possible pairs of samples. Gray dots indicate coalescence ratios for samples taken from the *same* tumor and blue dots indicate samples taken from *different* tumors. Coalescence ratios estimate the length of the shared ancestral lineage between two samples, divided by their total lineage length[15]. A coalescence ratio of 1 means that the two samples' ancestral lineages are identical, while 0 signifies no shared ancestral divisions. A large difference between coalescence ratios for samples from the same tumor (gray) vs. samples from different tumors (blue) suggests that the two tumors arose independently. Patient E3 had three separate primary tumors (PTa, PTb, PTc); each pairwise comparison is shown (bottom row). **b,** Comparison of intra-lesion heterogeneity between patient E15's two primary tumors. Pairwise angular distances among all combinations of samples within each primary tumor are shown.

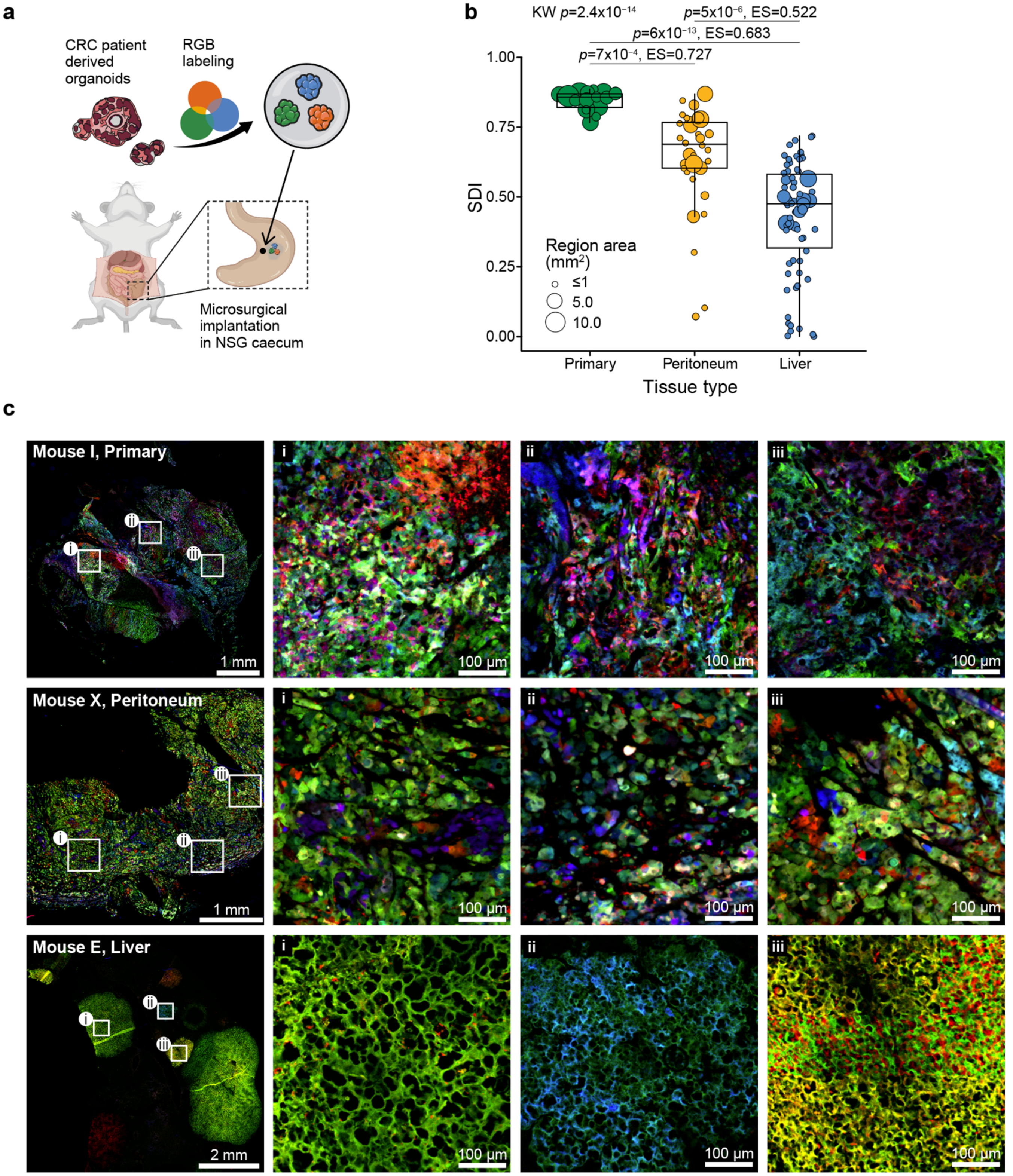
a
CRC patient derived organoids
RGB labeling
Microsurgical implantation in NSG caecum
b
KW p=2.4x10⁻¹⁴
p=5x10⁻⁶, ES=0.522
p=6x10⁻¹³, ES=0.683
p=7x10⁻⁴, ES=0.727
SDI
1.00
0.75
0.50
0.25
0.00
Region area (mm²)
≤1
5.0
10.0
Primary
Peritoneum
Liver
Tissue type
c
Mouse I, Primary
Mouse X, Peritoneum
Mouse E, Liver
i
ii
iii
1 mm
2 mm
100 µm

**Supplementary Figure 6. Intra-lesion heterogeneity visualized *in vivo* using optical barcoding. a,** Generation of orthotopic caecal tumors for multi-color lineage tracing in mice. Patient-derived organoids were transduced with multi-color lentiviral LeGO vectors and surgically implantated into the caeum of NSG mice (n=17 implanted primary tumors in total). **b,** Comparison of intra-lesion heterogeneity between mouse primary caecal tumors, peritoneal metastases, and liver metastases quantified by Simpson's diversity index (SDI) for discretized color species. Dot size represents the area of the corresponding region on the tissue section. **c,** Representative fluorescence microscopy images showing lineage heterogeneity in a surgically implanted mouse primary caecal tumor (top row), spontaneous peritoneal (center) and liver metastases (bottom). Left column shows tissue overview, highlighted regions (i, ii, and iii) are magnified in three right-most columns for each sample.

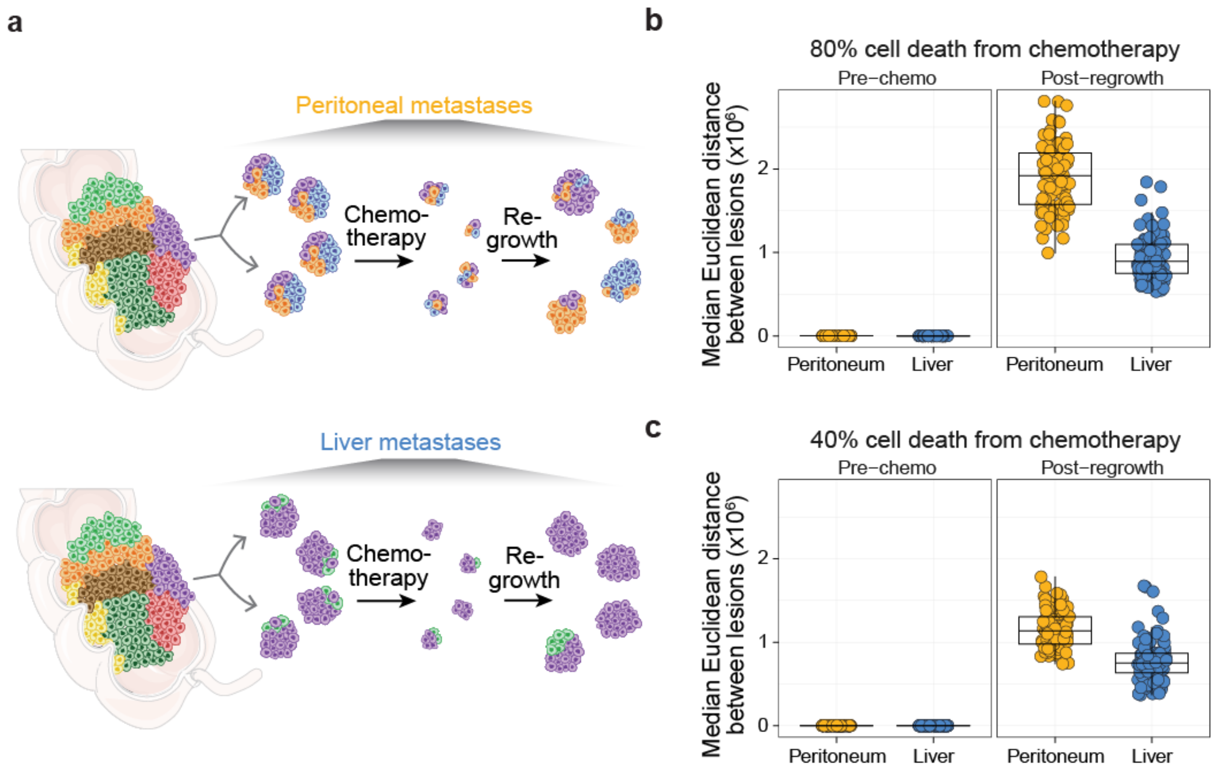


**Supplementary Figure 7. Simulated effects of chemotherapy on inter-lesion heterogeneity. a,** Schematic illustrating the hypothesized effect of chemotherapy on metastases containing high (peritoneal metastases, left) or low (liver metastases, right) levels of intra-lesion heterogeneity. After a fraction of cells in each lesion dies due to chemotherapy, lesions start re-growing according to a stochastic birth-death process and clone frequencies are remodeled by genetic drift. Inter-lesion divergence will be greater for metastases that harbored greater intra-lesion heterogeneity before treatment. **b,** Inter-lesion heterogeneity among metastases with high intra-lesion heterogeneity (peritoneal, yellow) and low heterogeneity (liver, blue) before treatment (left) and after 80% cancer cell death due to chemotherapy and subsequent regrowth (right). Data from 100 simulations is shown, please see methods for details on parameters. **c,** As in (b) but with 40% cancer cell death due to chemotherapy.

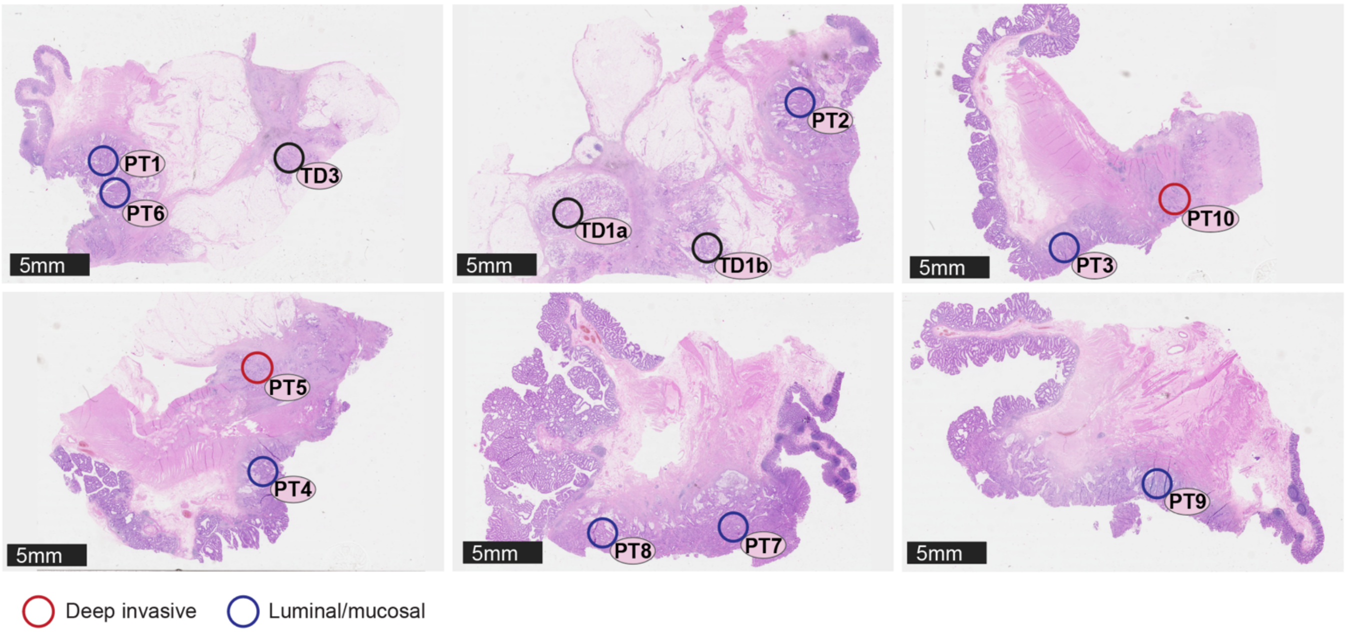


**Supplementary Figure 8. Hematoxylin and eosin stains of primary tumor regions from patient E14.** The locations of 10 primary tumor samples from patient E14 are shown (open circles). Deep-invading, red; luminal/mucosal, blue. Three tumor deposit regions were also sampled from the same FFPE blocks (locations shown in open black circles).

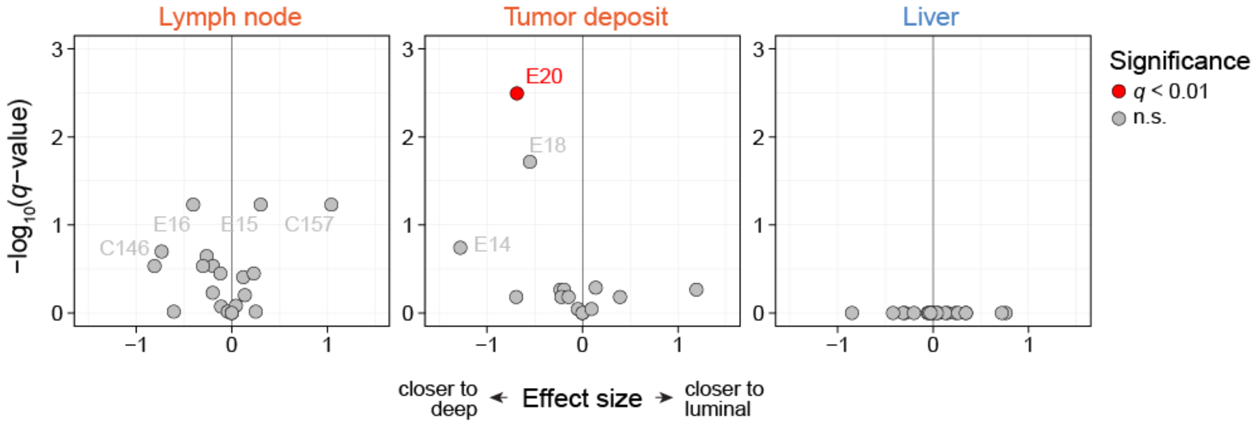


**Supplementary Figure 9. Association of lymph node metastases, tumor deposits and liver metastases with deep-invading vs. luminal primary tumor areas in each patient.** For each metastasis of a given type, we calculate the ratio of its angular distances to the closest deep-invading and closest luminal/mucosal region (lesion-depth ratio). This value is then averaged across all lesions to quantify their overall proximity to deep-invading vs. luminal regions. x-axis: $\log_2$-ratio of the *observed* average lesion-depth ratio to the *expected* average lesion-depth ratio (median of 10,000 permutations of primary tumor regions' invasion-depth labels within each patient). y-axis: $-\log_{10}$ *p*-values from two-sided permutation tests for each patient, with correction for multiple hypothesis testing (*q*-values). Patients with significant metastasis similarity to either deep-invading or luminal/mucosal regions are highlighted in red.

### Angular distance from microsatellite data

Cell samples' genotypes carry information on the evolutionary distances between the samples, which is our interest. However the genotypes are also influenced by sample purity, which we are not interested in. In this document we define the *angular distance* between cell samples in terms of microsatellite lengths, explain why angular distance is largely independent of purity, and relate angular distance to numbers of cell divisions between samples via a mathematical model of microsatellite evolution.

#### 1. Definition

To define the angular distance between samples, we first need to establish notation for the data. Consider that for a single sample of cancer cells $m$ microsatellites are sequenced. For the $i$th microsatellite, let $x_{i,j}$ denote the fraction of reads which have length $j$ (so $\sum_j x_{i,j} = 1$). Then let $x_i = \sum_j j x_{i,j}$ denote the mean length of the $i$th microsatellite and $x = (x_1, .., x_m)$ the vector of mean microsatellite lengths. Similarly, let $y = (y_1, .., y_m)$ and $z = (z_1, .., z_m)$ be the mean microsatellite lengths for another sample of cancer cells and a sample of normal (non-cancerous) cells respectively. So each sample is represented by a point in $m$-dimensional Euclidean space. We define the *angular distance* between the cancer samples $x$ and $y$ as the angle between $x$ and $y$ from the perspective of the normal sample $z$:

$$\begin{aligned} \text{angle}_z(x, y) \quad &:= \quad \arccos\left(\frac{(x-z)\cdot(y-z)}{|x-z||y-z|}\right) \qquad (1) \\ &= \quad \arccos\left(\frac{\sum_{i=1}^m (x_i - z_i)(y_i - z_i)}{\left(\sum_{i=1}^m (x_i - z_i)^2\right)^{1/2}\left(\sum_{i=1}^m (y_i - z_i)^2\right)^{1/2}}\right). \end{aligned}$$

In order to include normal samples as leaves on phylogenetic trees constructed by distance-based algorithms, we are also motivated to define the angular distance between the normal sample and cancer samples. We define the angular distance between the normal sample and any cancer sample as $\pi/3$. Although at first glance it may seem nonsensical to define the angle between two points in Euclidean space without the perspective of a reference point, the number $\pi/3$ is justified by an evolutionary model in the third section of this document.

#### 2. Purity

Now we 'prove' that angular distance is independent of purity. To begin, it is important to clearly state the assumptions that will underlie the theoretical argument - in order to establish a starting point for the line of reasoning, and also to bring to awareness potential limitations of the purity-independence claim. First, we assume that the impact of sequencing errors on mean microsatellite lengths can be neglected, so the observed genotypes $x, y, z$ represent the actual genotypes. Second, we assume that each microsatellite has a constant copy number among the normal and cancer samples. Third, we assume that the mean microsatellite lengths are constant among normal samples and among those normal cells which infiltrate cancer samples. The third assumption is motivated by our data which show that the variation in mean microsatellite lengths among normal samples is small compared to their variation among cancer samples.

These assumptions lead to a simple relationship between genotypes and purity. Suppose that the cancer sample with genotype $x$ has purity $p_x$, which means that it is composed of fraction $p_x$ cancer cells and fraction $1 - p_x$ normal cells. Denote the mean microsatellite lengths of specifically the cancer cells in that sample by $\tilde{x}$. Then the sample's genotype $x$ can be written as a linear combination of the cancer genotype $\tilde{x}$ and normal genotype $z$:

$$x = p_x \tilde{x} + (1 - p_x)z.$$

Similarly the genotype $y$ of the other cancer sample can be written as

$$y = p_y \tilde{y} + (1 - p_y)z,$$

where $\tilde{y}$ is the genotype of the cancer cells in the sample and $p_y$ is the purity. It follows that the angular distance between $x$ and $y$ is

$$\begin{aligned}
&\text{angle}_z(x, y) \\
&= \arccos\left(\frac{(x - z) \cdot (y - z)}{|x - z||y - z|}\right) \\
&= \arccos\left(\frac{p_x(\tilde{x} - z) \cdot p_y(\tilde{y} - z)}{|p_x(\tilde{x} - z)||p_y(\tilde{y} - z)|}\right) \\
&= \arccos\left(\frac{(\tilde{x} - z) \cdot (\tilde{y} - z)}{|(\tilde{x} - z)||(\tilde{y} - z)|}\right) \\
&= \text{angle}_z(\tilde{x}, \tilde{y}),
\end{aligned}$$

which is independent of the sample purities $p_x$ and $p_y$.

#### 3. Cell divisions

Towards drawing the relationship between angular distance and cell division numbers, we mathematically depict the genealogical structure of cells in a single human as a discrete, rooted, binary tree $\mathbb{T}$, where each node of the tree represents a cell.

Superimposed on this tree structure we consider a symmetric random walk model of microsatellite evolution: at cell division each of the $m$ microsatellites independently changes its length with some probability, with the mean increase in length equal to the mean decrease. For calculations, it will be helpful to provide explicit notation for these microsatellite mutations. Let $M_{a,i}$ for $a \in \mathbb{T}\backslash\{\text{root}\}$ and $i = 1,..,m$ be i.i.d. real-valued random variables with mean $\mathbb{E}M_{a,i} = 0$ and variance $\mathbb{E}M_{a,i}^2 = v$. Then writing $L_{\text{root},i}$, $i = 1,..,m$, for the microsatellite lengths of the tree's root, the microsatellite lengths of cell $b \in \mathbb{T}$ are given by

$$L_{b,i} = L_{\text{root},i} + \sum_{\text{root} \prec a \preceq b} M_{a,i},$$

where the summation is over all cells $a$ on the lineage between the root and $b$. A sample of cells is a subset of the tree $\mathbb{T}$. Consider a sample $X \subset \mathbb{T}$, and denote its mean microsatellite lengths by

$$x_i = \frac{1}{\#X} \sum_{b \in X} L_{b,i}.$$

Similarly, let $y_i$ and $z_i$ denote the mean microsatellite lengths of samples $Y, Z \subset \mathbb{T}$. Then the angular distance between $X$ and $Y$ with respect to $Z$ is given by

$$\arccos\left(\frac{\sum_{i=1}^m (x_i - z_i)(y_i - z_i)}{\left(\sum_{i=1}^m (x_i - z_i)^2\right)^{1/2}\left(\sum_{i=1}^m (y_i - z_i)^2\right)^{1/2}}\right) \tag{2}$$

matching the form of (1), which we think of as a random variable parameterised by the number of microsatellites $m$, the mutation rate $v$, and the genealogical structure of the samples $X, Y, Z$ with respect to the tree $\mathbb{T}$. Our task in this section is to calculate the probability distribution of the angular distance in terms of these parameters.

For the sake of simplicity, let's assume that the normal sample's genotype is well approximated by the root's genotype, that is we set $z_i = L_{\text{root},i}$, and let's further assume that the genotypes of each of the samples $X$ and $Y$ are well approximated by a mixture of the root genotype and the genotype of another cell: specifically, we set

$$x_i = p_x L_{a,i} + (1 - p_x)L_{\text{root},i} \quad \text{and} \quad y_i = p_y L_{b,i} + (1 - p_y)L_{\text{root},i}$$

for some purities $p_x, p_y \in (0, 1]$ and cells $a, b \in \mathbb{T}$.

Then the angular distance (2) becomes

$$\arccos\left(\frac{\sum_{i=1}^m (L_{a,i} - L_{\text{root},i})(L_{b,i} - L_{\text{root},i})}{\left(\sum_{i=1}^m (L_{a,i} - L_{\text{root},i})^2\right)^{1/2}\left(\sum_{i=1}^m (L_{b,i} - L_{\text{root},i})^2\right)^{1/2}}\right). \tag{3}$$

One component of (3) is

$$
\begin{aligned}
&(L_{a,i} - L_{\mathrm{root},i})(L_{b,i} - L_{\mathrm{root},i}) \\
&= \left( \sum_{\substack{c \in \mathbb{T}: \\ \mathrm{root} \prec c \preceq a}} M_{c,i} \right) \left( \sum_{\substack{c \in \mathbb{T}: \\ \mathrm{root} \prec c \preceq b}} M_{c,i} \right) \\
&= \sum_{\substack{c_1, c_2 \in \mathbb{T}: \\ \mathrm{root} \prec c_1 \preceq a \\ \mathrm{root} \prec c_2 \preceq b}} M_{c_1,i} M_{c_2,i},
\end{aligned}
$$

whose expected value is

$$
\begin{aligned}
&\mathbb{E}(L_{a,i} - L_{\mathrm{root},i})(L_{b,i} - L_{\mathrm{root},i}) \\
&= \sum_{\substack{c_1, c_2 \in \mathbb{T}: \\ \mathrm{root} \prec c_1 \preceq a \\ \mathrm{root} \prec c_2 \preceq b}} \mathbb{E}\left[ M_{c_1,i} M_{c_2,i} \right],
\end{aligned}
$$

thanks to linearity of expectation. But $\mathbb{E}\left[M_{c_1,i} M_{c_2,i}\right] = v$ if $c_1 = c_2$ and 0 otherwise. So

$$
\begin{aligned}
&\mathbb{E}(L_{a,i} - L_{\mathrm{root},i})(L_{b,i} - L_{\mathrm{root},i}) \\
&= v \times d(\mathrm{root}, a \vee b),
\end{aligned}
$$

where $a \vee b$ denotes the most recent common ancestor of $a$ and $b$ while $d(\mathrm{root}, a \vee b)$ denotes the number of cell divisons on the lineage between the root and $a \vee b$. Therefore by the law of large numbers, as the number of microsatellites $m$ tends to infinity

$$
\frac{1}{m} \sum_{i=1}^{m} (L_{a,i} - L_{\mathrm{root},i})(L_{b,i} - L_{\mathrm{root},i})
$$

converges almost surely to $v \times d(\mathrm{root}, a \vee b)$, and it follows that the angular distance (3) converges almost surely to

$$
\arccos\left( \frac{d(\mathrm{root}, a \vee b)}{d(\mathrm{root}, a)^{1/2} d(\mathrm{root}, b)^{1/2}} \right).
$$

This asymptotic view of angular distance has some appealing features. Most obviously, it is independent of the number of microsatellites and of the mutation rate, and it is of course independent of the purities $p_x$ and $p_y$ too. Its relationship to cell division numbers is also quite elegant. The arccosine's argument is the geometric mean of the ratio of the distance from root to most recent common ancestor relative to the distance from root to leaves. Specialised to an ultrametric tree, the distance from root to leaves is constant so the angular distance simplifies further to

$$
\begin{aligned}
\mathrm{angle}_z(x, y) &= \arccos\left( \frac{d(\mathrm{root}, a \vee b)}{d(\mathrm{root}, \mathrm{leaf})} \right) \\
&= \arccos\left( 1 - \frac{d(a, b)}{2 d(\mathrm{root}, \mathrm{leaf})} \right), \qquad (4)
\end{aligned}
$$

which shows that the angular distance is an increasing function of the distance between $a$ and $b$ normalised by the distance from root to leaves. In other words the angular distance, which is a distance between genotypes, is an increasing function of genealogical distance. This latter property might sound unsurprising but it is crucial, indicating that topology of a phylogenetic tree constructed via angular distance matches the true genealogical structure.

Finally we come to the justification for defining the angular distance between the normal and cancer samples as $\pi/3$. Assuming ultrametricity, choose any $a, b \in \text{leaves}(\mathbb{T})$ such that $d(a, b) = d(\text{root}, \text{leaf})$. According to (4) the angular distance between $a$ and $b$ is $\arccos(1/2) = \pi/3$. Therefore the angular distance between root and leaf is naturally defined as $\pi/3$.

To conclude, let's emphasise that this modelling discussion is extremely idealistic. It is based on numerous simplifying assumptions and should be thought of only as a cartoon of the true biology.

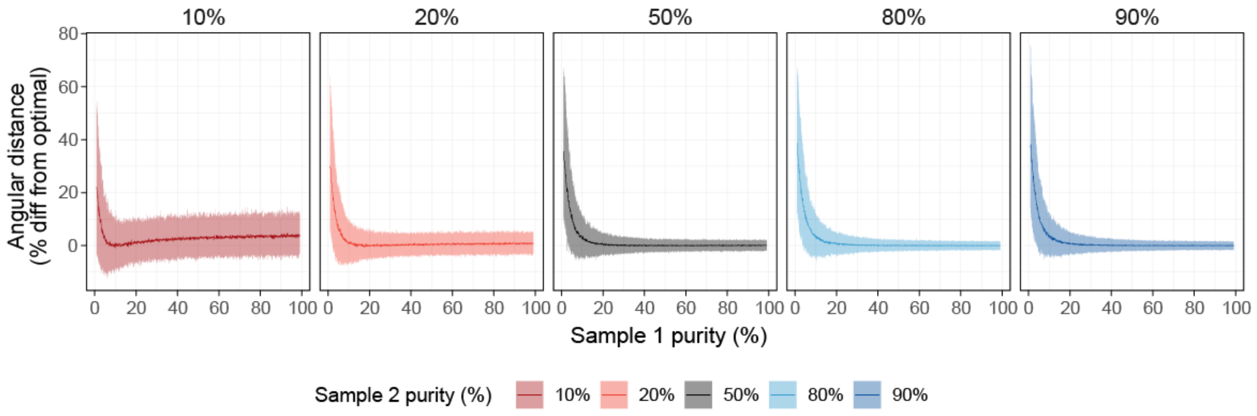


**Supplementary Note Figure 1. Simulated effects of tumor impurity on angular distance**
Angular distances between two simulated tumor samples with known purities are compared to their optimal values (perfect purity in both samples). For each pair of tumor sample purities, 200 simulations were executed and their angular distances under impure conditions were compared to the optimal condition of 100% purity. Y-axis: percent difference from optimal conditions. X-axis: purity of simulated tumor sample 1, with values ranging from 5% to 95%. Different colors indicate the purity of simulated tumor sample 2 (also labelled above each plot). Dark colors show the median percent difference of 200 simulations. Lighter color bands show 95% confidence intervals.
